# Spatial, transcriptomic and epigenomic analyses link dorsal horn neurons to chronic pain genetic predisposition

**DOI:** 10.1101/2022.04.01.486135

**Authors:** Cynthia M. Arokiaraj, Michael J. Leone, Michael Kleyman, Alexander Chamessian, BaDoi N. Phan, Bettega C. Lopes, Vijay Kiran Cherupally, Myung-Chul Noh, Kelly A. Corrigan, Deepika Yeramosu, Michael E. Franusich, Riya Podder, Sumitra Lele, Stephanie Shiers, Byungsoo Kang, Meaghan M Kennedy, Viola Chen, Ziheng Chen, Richard P. Dum, David A. Lewis, Yawar Qadri, Theodore J. Price, Andreas R. Pfenning, Rebecca P. Seal

## Abstract

Key mechanisms underlying chronic pain occur within the neural circuitry of the dorsal horn. Recent genome-wide association studies (GWAS) have identified genetic variants associated with the predisposition to chronic pain. However, most of these variants lie in regulatory non-coding regions that have so far not been linked to spinal cord function. Here, we take a multi-species approach to determine whether chronic pain variants impact regulatory elements of dorsal horn neurons. We first built a more comprehensive single cell atlas; filling gaps by generating a high-quality Rhesus macaque atlas and integrating it with human and mouse. With cellular-resolution spatial transcriptomics, we mapped the laminar distributions of the resulting species-conserved neuron subtypes, uncovering an unexpected organization. Lastly, we generated a mouse single-nucleus open chromatin atlas to partition the heritability of chronic pain traits. From this, we identified strong, selective associations between specific, conserved neuron subtypes and major forms of chronic pain.

## Introduction

The spinal cord dorsal horn is the gateway for somatosensory integration and transmission of normal sensory processing as well as for maladaptive states such as chronic pain^1,2^. It is organized into dorsoventral laminae (I-VI); that have roles defined in part by termination patterns of functionally distinct class(es) of primary sensory neurons^3–9^. The superficial laminae (I-II) receive input from sensory neurons that convey temperature, itch, crude touch and pain while the deeper laminae (III-V) receive input from sensory neurons that convey crude touch, and proprioception. The deeper laminae have also been implicated in sensorimotor transformations. Projection neurons reside in laminae I, and III-V and innervate many brain regions including the thalamus, parabrachial nucleus, dorsal column nuclei and cerebellum.

Until recent single-cell transcriptomic studies of mouse and adult human spinal cord, interneurons were assigned to cell type classifications and functionally studied based on the expression of one or two gene markers^10–15^. Though these single-cell datasets provide a much richer understanding of the molecular composition and organization of the dorsal horn, barriers to bridging rodent models and human pain disorders still exist. For example, transcriptomic analysis from human donors is complicated by factors such as their limited availability, control of experimental conditions, and reduced cell viability. As for mouse, the evolutionary distance from humans can result in significant molecular, physiological, and anatomical differences. Inclusion of non-human primates, which are evolutionarily closer to humans and hence valuable translational models, overcomes many of these limitations.

In parallel with the advances in the molecular characterizations of cell types, recent genome-wide association studies (GWAS) have identified dozens of loci confidently associated with the genetic predisposition to chronic pain^16^. Prior research in both human and animal models suggests that the genetic risk for chronic pain impacts the function of both primary sensory and dorsal horn neurons^12,17,18^. Previous studies have established some links between chronic pain-associated single-nucleotide polymorphisms (SNPs, single-nucleotide variants with at least 1% prevalence of each allele), and the SNP proximity to genes most expressed in specific cell types^17,18^. However, most of the genetic risk burden for chronic pain is from several common, small effect-size variants residing in non-coding genomic regions. Many of these variants likely disrupt the function of distal, cell-type-specific *cis*-regulatory enhancers^19–22^, which are abundant in open chromatin and can lie far away from the genes they regulate^21,22^. This genetic architecture is common for polygenic traits, including those with a neural basis^23,24^. Without direct measures of cell-type-specific regulatory regions, genetic analyses are limited to annotations via marker genes, bulk epigenomic measures, or expression quantitative trait loci (eQTLs), which are loci associated with changes in gene expression, an indirect measure of gene regulation. Notably, two recent analyses based on gene expression and eQTLs of chronic pain GWAS found no significant enrichment in bulk spinal cord RNA-seq^25,26^.

Thus, to more fully account for the spinal cord’s potential genetic risk of chronic pain disorders, a detailed epigenomic resource that provides functional annotations at the single-cell level is required. Such data can then be used to link human genetics to species-conserved cell types studied in rodent and primate pain models. In this work, we provide the first comprehensive connection between species-conserved neuron subtypes of the dorsal horn using a single nucleus epigenomic dataset we developed, and their genetic risk for chronic pain.

First, we generated a single-nucleus transcriptomic atlas of the young-adult Rhesus macaque dorsal horn and compared gene expression profiles to the publicly available mouse and human spinal cord datasets. We found high conservation of macaque cell type identities across all species, but less so for individual marker genes, leading us to identify new marker genes that better reflect conserved expression. Based on these conserved markers, we introduce a new nomenclature for these cell types that reflects better harmonization across datasets and species. We also compared the laminar patterns of these cell types in macaque, and mouse using two single-cell resolution assays, RNAscope and Xenium spatial transcriptomics, respectively^27,28^. Our analysis revealed an interesting relationship between the molecular uniqueness of a neuronal subtype and its laminar location, and it resolved a major discrepancy about the location and identity of an elusive cell type, Exc-PBX3/PDYN. Despite comparing disparate species and using significantly different technologies, we found highly consistent laminar distributions, across datasets providing multi-modal evidence of cross-species conservation.

We then assessed the conserved neuron subtypes for human chronic pain genetic risk using the mouse open chromatin dataset, with a particular focus on distal regulatory regions such as enhancers, which are found within open, accessible chromatin. A single cell atlas of open chromatin in the mouse dorsal horn was generated using the single-nucleus Assay for Transposase Accessible Chromatin (snATAC-seq)^19,20^. We labeled the nuclei using snRNA-seq integration, and then robustly mapped mouse open chromatin to human coordinates. Using a statistical method for partitioning heritability of traits by functional region, we show^29^ for the first time that the open chromatin regions of specific conserved neuron subtypes are indeed significantly enriched for human genetic variants linked to multiple chronic pain phenotypes. We also identified two genetic variants from the chronic pain GWAS that are particularly compelling because they intersect with open chromatin peaks of specific neuron-subtypes, indicating potential disruption of enhancer function by these variants in chronic pain disorders.

In summary, we provide a set of data resources and a framework for decoding the logic of dorsal horn somatosensory circuits in animal models with greater generalizability to human genetics, physiology, and disease. These resources include a Rhesus macaque transcriptomic atlas, a new cell type framework of cell conservation, a mouse single cell open chromatin dataset, and a mouse single cell spatial dataset. We leveraged these new resources to provide the highest resolution analysis to-date of evolutionary conservation, spatial organization, open chromatin, and genetic risk of the dorsal spinal cord. The work will enable the field to make further discoveries about dorsal horn anatomy and function with greater precision, cross-species generalizability, and medical translatability.

## Results

### Major Cell Types of the Macaque Dorsal Horn

We first sought to build a more detailed cellular map of the primate dorsal horn that would also fill gaps in the human map to further facilitate the translation from rodents and expand the resources of an important translational model system. We thus performed single-nuclear RNA sequencing (snRNA-seq) on lumbar tissue (L4-5) harvested from young adult (3-year-old) male Rhesus macaques (n=3) using the 10X Genomics Chromium platform. For the isolation of the dorsal horn nuclei, the spinal grey matter of each animal was dissected, transected at the central canal, and mechanically dissociated (**Figure 1A**). Deep sequencing of the single nucleus libraries produced an average of ∼50M reads per animal. To optimize alignment of the reads to the macaque genome (rheMac10), we employed a custom annotation method (**Methods**) that mapped 91% of the reads to the genome^29^. After quality control filtering, 12,243 nuclei were retained for further analyses. The three biological replicates were then co-normalized to remove cell-specific read sampling biases using Scran and subsequently integrated and batch-corrected with Scanorama (**Figure S1B**)^30,31^. Nuclei were assigned to their major cell type based on the expression of well-established marker genes. An average of 1,720 genes and 3,700 UMIs were detected per cell type (**Figures 1B-D** and **S1A,C**)^11^. The neuronal marker, RNA Binding Fox-1 Homolog 3 (*RBFOX3)*, identified 2,698 nuclei, comprising ∼22% of the total nuclei (**Figure 1C-D**). The largest cell class was oligodendrocytes (39%, MBP, **Figure 1C-D**), which could be further sub-clustered into two groups based on the expression of either quinoid dihydropteridine reductase (Oligo 1, *QDPR*), or Dpy-19 like C-mannosyltransferase 1 (*DPY19L1*) together with S100 calcium binding protein B (*S100B*) (**Figure 1C-D**). Astrocytes were also sub-clustered into two groups based on high (Astrocyte 1) or low (Astrocyte 2) expression of Glial fibrillary acidic protein (*GFAP*). Oligodendrocyte precursor cells, microglia, meninges, ependymal cells and Schwann cells all make up smaller individual clusters (**Figure 1C-D**).

**Figure 1.**
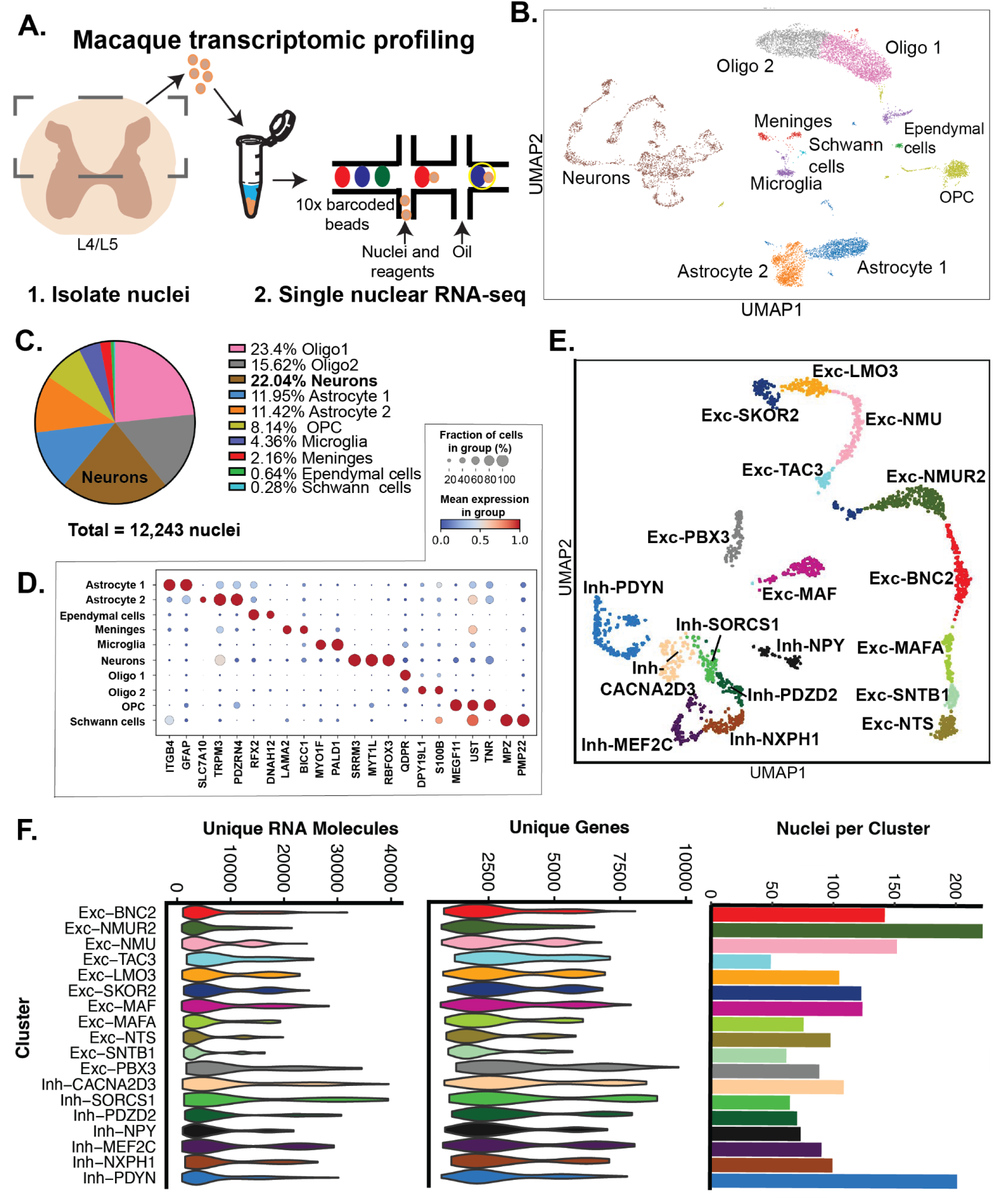
snRNA-seq and Cell Type Identities of the Macaque Dorsal Horn. **A,** Schematic overview of the single-nuclear RNA-sequencing (snRNA-seq) experimental workflow of the macaque lumbar dorsal horn. **B.** UMAP representation of major cell types identified by established marker genes. **C,** Pie chart representation of the contribution of each major cell type present in the macaque dorsal horn. **D,** Dotplot of the normalized mean expression of key marker genes used to identify each major cell type. The size of the circles depicts the percentage of cells within each cluster that express the gene. **E,** UMAP visualization of the 1,954 dorsal horn neuronal nuclei identified through Leiden clustering. Excitatory clusters are prefixed with **Exc** and inhibitory clusters with **Inh**. **Table 1** indicates naming and relationship to other cell types, and the rationale for naming is described in **Results**. **F,** Quality control and number of nuclei per cluster, Left: Violin plot of per-cell Unique RNA molecules by cluster, Middle: Violin plot of number of unique genes per cell found by cluster, Right: number of nuclei present in each cluster.

### Transcriptomically Defined Macaque Neuronal Clusters

We next sub-clustered the neuronal nuclei into 18 distinct clusters that could be broadly characterized as being either excitatory or inhibitory based on the known gene markers: vesicular glutamate transporter 2 (*SLC17A6*) for excitatory and vesicular inhibitory transporter (*SLC32A1*), as well as GABA synthesizing genes *i.e.* glutamate decarboxylase 1 and 2 (*GAD1 and GAD2*) for inhibitory (**Figure S2A**)^32^. Interestingly, two of these clusters were unusual in that they each contain both excitatory and inhibitory nuclei, but otherwise have indistinguishable gene expression profiles (**Figure S2A).** This is characteristic of clusters previously identified to be a part of the intermediate and ventral regions of the mouse spinal cord (MidVent)^11^. Because of the difficulty in further refining cell identities within this cluster, we exclude these cells from downstream snRNA-seq analysis but consider their spatial distributions later on (see **Results: Exc-PBX3/PDYN is a distinct dorsal horn cell type, while *Spp1* neurons previously called MidVent are prevalent in the deep dorsal horn)**.

Our final dataset, excluding the Spp1 domain, consists of 1,954 nuclei, with 64% classified as excitatory and 36% as inhibitory. Note, we further sub-clustered two inhibitory clusters (Inh-SORCS1/Inh-PDZD2 and Inh-MEF2C/Inh-NXPH1) based on interesting discrepancies we observed within these neuron subtypes between mouse and macaque (see Results: **Cross-Species Comparisons, Figure S2C**). Thus, from these nuclei, we derived 18 distinct cell types, 11 excitatory and 7 inhibitory (**Figure 1E).** Each neuron cluster has an average of 139 nuclei with a range of 49 to 222 nuclei per cluster, and each nucleus with an average 3,196 distinct genes detected and 8,010 UMIs (**Figure 1F**). Relationships between the 18 individual clusters were visualized using the Ward’s hierarchical clustering method (**Figures S2B**). To provide a unified nomenclature for the dorsal horn neuron subtypes that apply to mammals more broadly, we assigned and hereafter use new names that are based on conserved marker genes we identified from our cross-species integration of macaque, mouse, and human single cell datasets (described in more detail later) (**Table 1**).

### Macaque and Human Laminar Distribution of Marker Genes

The functional organization of the dorsal horn is to a degree imparted by its laminar organization. To determine how the transcriptomically-defined macaque clusters are distributed within the Rexed laminae of the lumbar dorsal horn, we identified macaque-specific marker genes that we then mapped in the dorsal horn using RNAscope^®^ (**Figure 2A)**. For some clusters, a combination of marker genes was used to achieve sufficient representation of the nuclei that belong to the cluster. RNAscope^®^ fluorescence *in situ* hybridization was performed with these combinations of marker genes on L4-5 spinal cord slices taken from both male and female macaques (**Figures 2B-F, S4, S5**)^27^.

**Figure 2.**
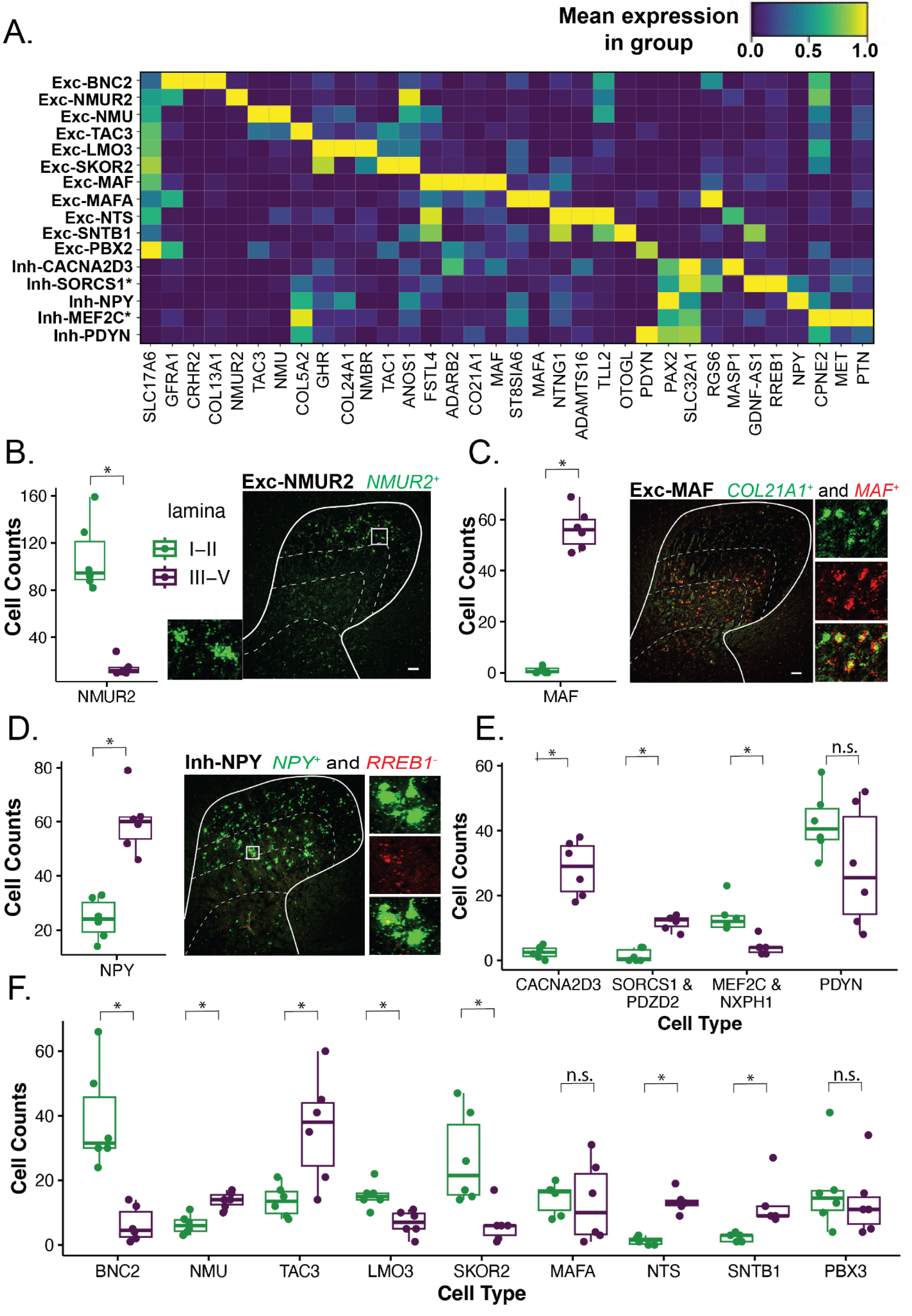
Macaque-specific markers of dorsal horn neuron subtypes and RNAscope In situ Validation. **A,** Heatmap of the normalized mean expression of macaque-specific marker genes (mentioned in Figure SX, dendrogram) for the neuronal subtypes. Inh-SORCS1* refers to the combined Inh-SORCS1/Inh-PDZD2 population, and Inh-MEF2C *refers to the combined Inh-MEF2C/Inh-NXPH1 population. **B-D,** RNAscope results for the selected neuron subtypes **B,** Exc-NMUR2 **C,** Exc-MAF, and **D,** Inh-NPY. Panels depict the *in situ* hybridization marker gene combinations used to detect each target cell type, with the color of the marker name indicating the fluorescence of the marker in the image. Dorsal horn images are taken at 10x with the smaller insets showing a magnified image (20x) of the individual gene(s) as well as the merged image. Laminar boundaries (dashed lines) are drawn between II/III, III/IV and IV/V. Scale bars in bottom right corners of panels = 100 μm. Box plots indicate the binned laminar distribution (shallow=I-II, deep = III-V) of cell counts by sample: n=5 or 6 spinal cord sections from N=2-3 macaques. *p<0.05.

Nearly all cluster markers showed expression patterns that were significantly enriched in either the superficial (I-II) or deep dorsal horn (III-V) laminae (**Figures 2B-F, S4-5**). For example, the Exc-NMUR2 cluster markers are located almost exclusively in the superficial laminae (**Figure 2B**) while markers for Exc-MAF are enriched in the deep dorsal horn (**Figure 2C**). The Inh-SORCS1/Inh-PDZD2 cluster expresses Neuropeptide Y (*NPY*) and *RREB1*, whereas the Inh-NPY cluster only expresses *NPY*. We therefore defined the Inh-NPY cluster as positive for *NPY* and negative for *RREB1* expression (**Figure 2D**). Overall, inhibitory populations had a significant association with either superficial or deep laminae with the exception of Inh-PDYN (**Figures 2E, S5**). No difference was observed for cluster markers with respect to sex.

We also examined the distribution of neuropeptides in the dorsal horn of macaques and humans. The dorsal horn is rich in neuropeptides that are involved in many aspects of somatosensory processing including NPY, CCK, GRP and TAC1. The laminar distribution of genes encoding neuropeptides has been well characterized in mice, but whether this laminar distribution is conserved in macaques and humans is not known. Using RNAscope^®^ we found a high conservation of the overall laminar distributions of neuropeptide genes between macaques and humans that are similar to those reported for mice (**Figure S6, Tables S1-2**). We also observed co-expression of certain neuropeptide genes such as *NMU* and *TAC3 (Tac2* in mice) in lamina II, which has also been reported for mice. One exception we observed was a higher percentage of *CCK* expressing cells in the superficial laminae of humans compared to what we observed in macaques and what has been published for mice^33–35^.

### Cross-species Comparisons of Macaque, Human and Mouse Dorsal Horn Neuronal Clusters

Two additional snRNA-seq atlases of the spinal cord were recently made publicly available: (1) a meta-analysis that unified six datasets from the mouse^12^, and (2) nuclei from human donors^17^. To determine the extent of molecular and cellular conservation between species, we compared the gene expression and cell types in our macaque dataset to the adult human dataset and the adult subset of the mouse atlas. First, we performed a focused comparison between macaque and mouse datasets. Details are discussed in **Methods.** Briefly, each neuronal nucleus from the macaque was compared to each nucleus from the mouse based on the top 100 marker genes from the mouse meta-analysis (**Figure S2C-D**, **Table S3**). Interestingly, the macaque Inh-SORCS1/Inh-PDZD2 and Inh-MEF2C/Inh-NXPH1 clusters show a poor correspondence to the mouse clusters (described in more detail below). Upon closer examination, however, we observed that macaque UMAPs of orthologous genes for two of the mouse family markers, *RORB* and *ADAMTS5*, show an interesting separation within the macaque clusters (**Figure S2C**), and that further sub-clustering revealed a strong correlation of Inh-SORCS1 and Inh-MEF2C with the mouse Rorb family and of Inh-PDZD2 and Inh-NXPH1 with the Adamts5 family (**Figure S2C-D**). For this reason, we separated them into four independent but closely related cell types.

We next integrated all three datasets, macaque, human and mouse, beyond the cluster level to visualize the similarity of single nuclei. The datasets were normalized and integrated simultaneously using Seurat and were found to be highly overlapping, with minimal experimental and species-specific differences **(Figure 3A, left: macaque, middle: human, right: mouse**). Regardless of species, cells expected to be of the same type based on our macaque-mouse comparison and a previous comparison between human and mouse, were aligned in the UMAP, suggesting that cell type signatures drive the visualization and clustering. This is more formally supported by silhouette scores for each cell type (**Figure S7)** and label transfer, which uses the high-dimensional relationships of nearest neighbors to determine identity. Label transfer between mouse and human reproduced the same cell type mapping previously identified by Yadav *et al.* (**Figure S8, compared to Figure 5C of** ^17^). Additionally, per-cell label transfer results are shown from macaque to mouse and to human in **Figure S9**, and subtype homologies are also summarized in **Table S4**. Nearly all mouse and human cell types were consistently mapped to a single macaque cell type, with the exceptions of Exc-TAC3 and Exc-PBX3. Exc-TAC3 is a smaller cluster closely related to Exc-NMU, the latter of which did label several mouse and human cell types. Mouse cells from Excit.20 and Excit.30 mapped to Exc-PBX3, but none of the human cells mapped to Exc-PBX3. However, because we used the subsets of the human and mouse datasets that were labeled as *dorsal horn*, and based on our own macaque clustering showing that Exc-PBX3 is closely related to Mid and Ventral cells, we theorized that Exc-PBX3 might better match to a cell type labeled within what others have called MidVent. We thus re-integrated mouse and macaque datasets but this time included all mouse neuron cell types, and found that Exc-PBX3 is closely aligned with the cell population Excit.25 from mouse (**Figure S10**), a MidVent cluster homologous to the Ex-M-2 population in humans^17^. Overall, our results strongly support the delineation and integration of robust and consistent neuronal cell populations across all three species.

**Figure 3.**
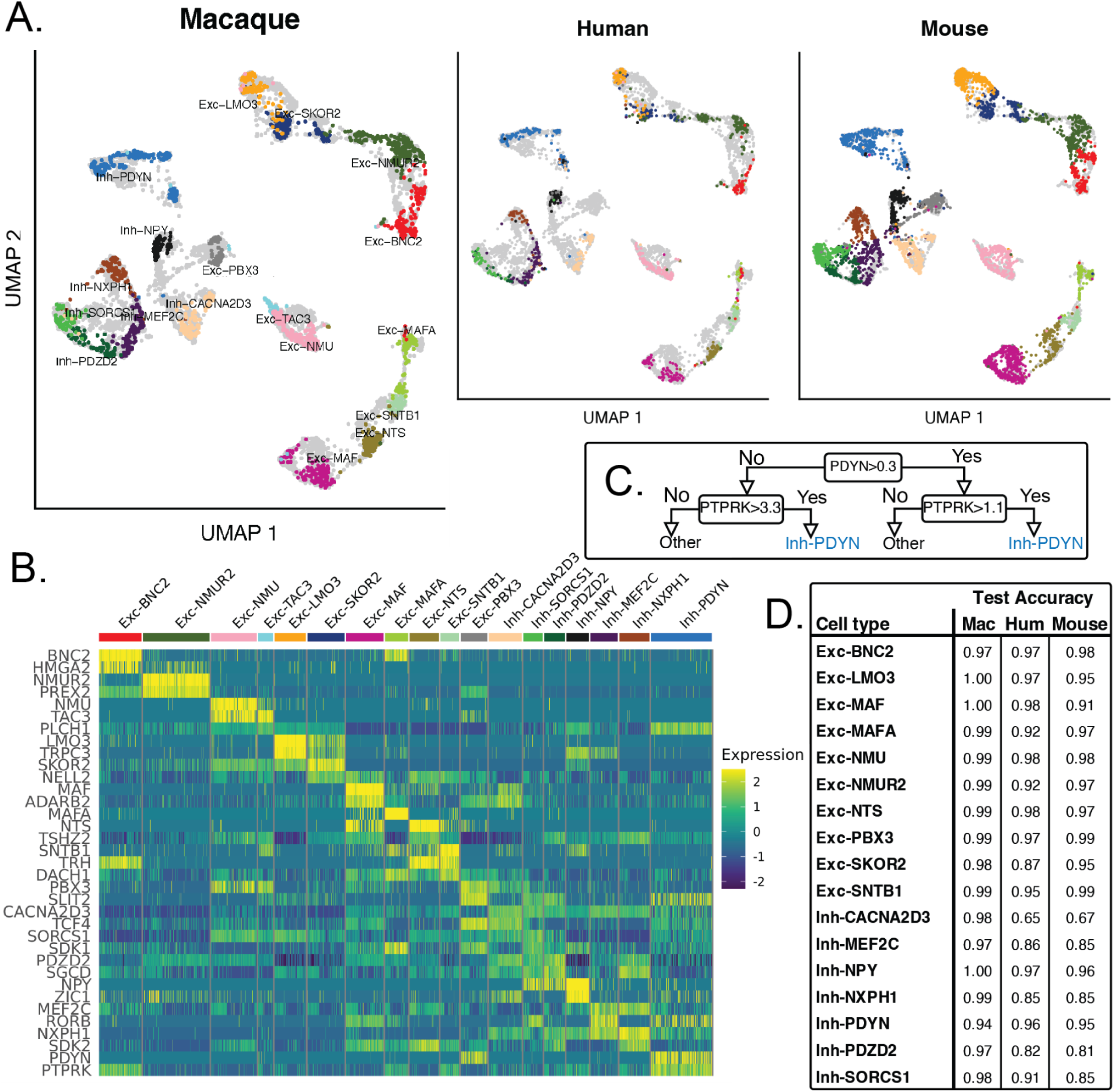
snRNA-seq integration of adult macaque, human, and mouse dorsal horn datasets; identification and validation of new conserved marker genes. **A,** UMAP representations of dorsal horn neuron subtypes of each adult species in integrated coordinates (Macaque, left; Human, middle; mouse, right). Overlayed, colored dots represent nuclei from the species titled, light grey background dots represent nuclei from the other species. The neuron subtype color scheme in **panel B** applies to UMAPs. **B,** Scaled gene expression of selected conserved marker genes. Rows are individual marker genes and columns are each neuron subtype. Each colored tick represents the expression of the given marker for an individual nucleus. The full correspondence between sets of marker genes and each cell type is given in **Table 1**. **C.** Schematic of an example decision tree to validate the choice of conserved marker genes for a given neuron subtype, in this case Inh-PDYN. The decision tree learns expression cutoffs that best separate the target cell type from off-target cells, trained using 80% of the macaque nuclei. **D,** Decision tree test accuracies are reported for each cell type and across the test sets of the three species: the remaining 20% of the macaque nuclei, and all of the human and mouse nuclei. Mac=Macaque. Hum=Human. Additional metrics are provided in Table S9-12.

To facilitate reference to and study of these integrated cross species neuronal subtypes, we sought to identify marker genes that demonstrate consistent cross-species expression levels for each cell type. First, we identified per-species differentially expressed genes using Seurat (**Tables S5-7**) and differentially expressed genes from the integrated dataset (**Table S8**) to guide the choice of candidate markers. We next selected two to three genes per cell type that would separate neuron subtypes well based on visualization (**Figures 3B, S11**). To quantify this separability, we trained decision trees (**Figures 3C, S13**) to optimally distinguish cell types with just the selected genes. We trained our models using 80% of the macaque data (*training set*) and evaluated performance on the heldout 20% of macaque data as well as the full data for human and mouse (*test sets*). We performed this analysis on scaled, imputed^36^ RNA expression data because this better reflects dataset-independent relative differences and better approximates gene expression across cells. As expected, test performance was highest for macaque (**Figure 3D**), yet we found overall strong model performance across all species for identifying the target cell type (**Tables S9-12**).

Given our quantitative findings, we named the cell types based on these cross-species marker genes (**Table 1**), provided nicknames for them based on the first most specific marker gene, and created visualizations of marker gene expression in macaque (**Figure 3B**), human and mouse (**Figure S11**). This nomenclature is therefore used throughout the paper.

### Species-specific differences in Gene Expression

We then used the atlas to analyze the cell type expression of several marker genes commonly used to label dorsal horn neuron populations for functional studies in mice. Some of these genes had low absolute expression in multiple datasets, which would then be artificially inflated by imputation. Thus, we visualized un-imputed expression in our macaque dataset, versus in the adult mouse^11,12^ and human datasets^17^, grouping cells based on our conserved cell type annotations (**Figure S12**). In this comparison again, most, but not all genes show species conserved cell type enrichment (see **Discussion).**

Identification and laminar distribution of conserved dorsal horn neurons in highly multiplexed *in situ* spatial transcriptomics of mouse spinal cord We next examined the laminar distributions of the integrated neuron subtypes we identified in mouse dorsal horn *in situ* and compared it to what we observed in macaque. We used the Xenium *in situ* spatial transcriptomics assay, which sensitively measures the presence and position of hundreds of distinct genes in a single experiment at single-cell resolution. To test if we could recover the conserved neuronal subtypes in Xenium, we performed a simulation using the mouse snRNA-seq dataset with the 248 genes available in the mouse brain panel^28^. With the high per-gene sensitivity (simulated by imputation^36^) that is likely achievable with Xenium, we were able to distinguish the conserved neuronal subtypes (**Figure S14**). We next assayed adult mouse spinal cord slices with the mouse brain panel. Cell labeling was performed using the software Seurat^37^, taking a spatial-agnostic approach. We first integrated the full Xenium dataset with the full adult mouse snRNA-seq dataset, which yielded distinct, identifiable glial clusters and probable neuronal clusters (**Figure S14A-B**). The resulting representation of cell types after labeling is shown in **Figure 4A**. **Figure 4B** shows a representative slice with the cells labeled as in **Figure 3A**. The location of specific cell types faithfully match expected locations based on prior anatomical studies: for example, Schwann cells lie in putative peripheral nerves, leptomeningeal cells line the spinal cord edge, oligodendrocytes and OPCs are most abundant in white matter, while neurons are limited to grey matter with the highest density in the dorsal horn.

**Figure 4.**
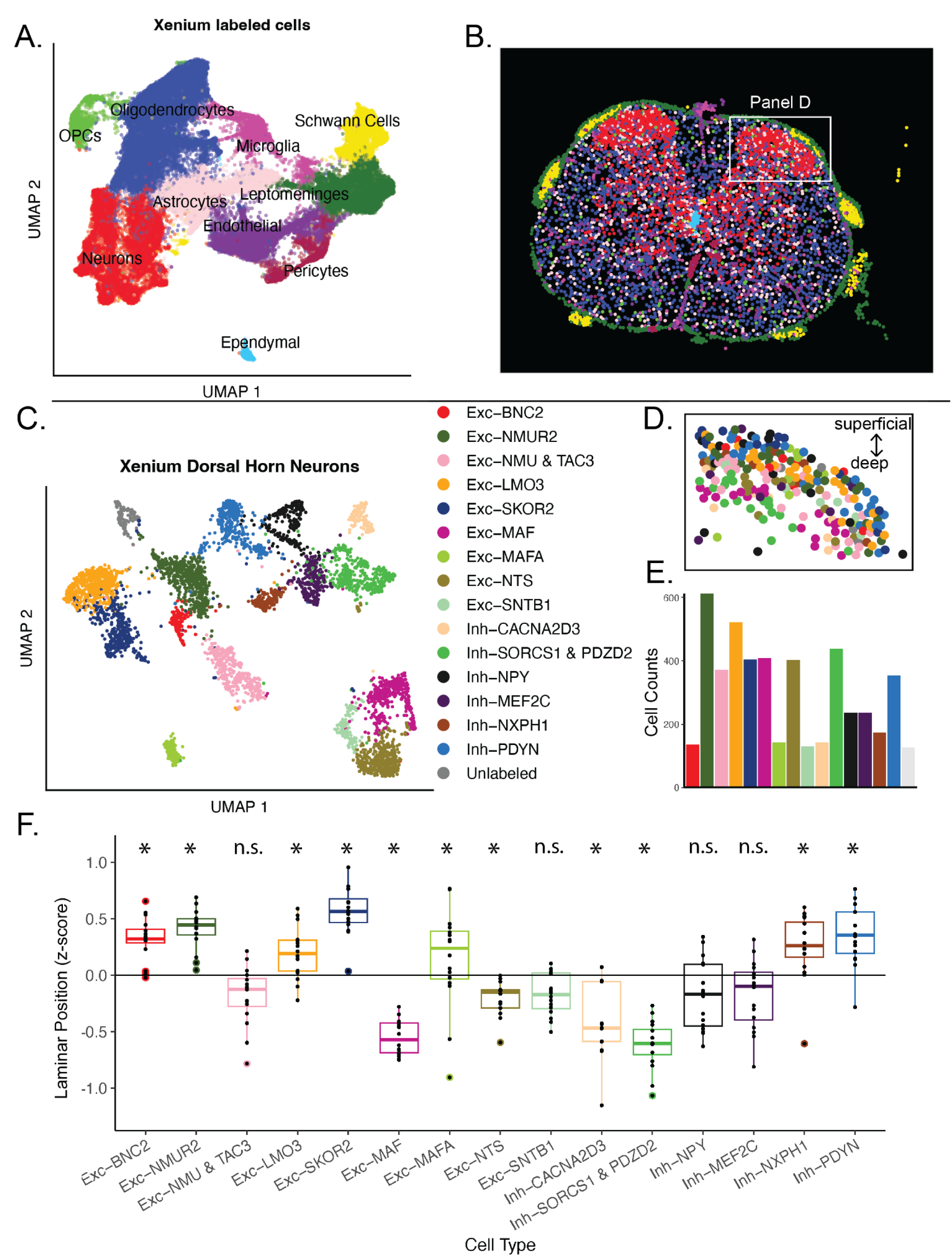
Dorsal horn neuron cell types identified in mouse Xenium® Spatial Transcriptomics assay and calculation of laminar distributions. **A,** UMAP representation of major cell types identified in Xenium assay based on integration with adult mouse snRNA-seq dataset. **B,** Spatial distribution of major cell types in spinal cord cross-section. Individual dots represent individual cells segmented by Xenium. Dot color indicates cell type; the UMAP in panel A indicates the major cell type for a given color. Box indicates location of representative dorsal horn section, shown in Panel D. **C,** UMAP representation of dorsal horn neuron subtypes identified in Xenium assay based on integration with adult mouse snRNA-seq dataset with conserved labels. Inh-SORCS1 and Inh-PDZD2 are combined as a population as individual cell types were not readily distinguishable in Xenium. One cluster indicated as Unlabeled did not correspond to any snRNA-seq dorsal horn cell type. **D,** Cell type distribution of boxed region shown in Panel A. Superficial position defined as higher vertical coordinate in Xenium. Some trends are visually apparent, such as deeper location of Exc-MAF cells (purple). **E,** Bar chart of cell counts per neuron subtype in Xenium. **F,** Box plot of laminar shallowness per cell type. z-score of laminar shallowness was the shallowness of an individual cell relative to the average y-coordinate of its slice and normalized by the variability in y-coordinates of the slice. Individual points of each box plot represent the average z-score for the cells of that cell type in one slice. * significant p-value indicating difference from zero (either significantly deep or shallow), after false discovery rate correction of 0.05.

We then integrated the putative neuron clusters with the species-conserved dorsal horn neuron subtypes identified with snRNA-seq. Most clusters aligned (**Figure S14C-D**), but some putative neuronal clusters lacked a match, cells likely belonging to the population referred to as MidVent. Integrating the datasets after removing this large cell group led to a clear correspondence between each of the Xenium identified clusters and the conserved dorsal horn neurons labeled in the Sathyamurthy snRNA-seq dataset (**Figure S14E-F**). One exception to this was a lack of separation of the closely related subtypes, Inh-SORCS1 and Inh-PDZD2, even if they were clustered at higher resolutions. The Xenium neurons assigned to conserved neuron subtypes are shown in **Figure 4C**. Neuron subtype counts are shown in **Figure 4E**, along with glial cell counts **(Figure S16A)** and the numbers of genes and transcripts for both neurons and glia **(Figure S16B-E)**. Although several conserved marker genes from **Figure 3D** were absent from the Xenium panel, several were present: Pdyn, Plch1, Nts, Sntb1, Pdzd2, Rorb, and Sdk2; and all showed specificity for their corresponding neuron subtypes (**Figure S21**). Additionally, while parvalbumin (PVALB) was not detected in the snRNA-seq datasets we analyzed, it was clearly expressed in the Xenium dataset by the Inh-CACNA2D3 subtype (**Figure S18A**), and less robustly by Inh-SORCS1, Inh-PDZD2, and Exc-MAF subtypes. PVALB inhibitory neurons are critical to the development and maintenance of mechanical hypersensitivity after injury, and their more precise assignment to particular inhibitory cell types will facilitate mechanistic studies^38,39^. The expression of PVALB in the Exc-MAF subtype is consistent with previous studies showing overlap of this gene with *Cck*^38^.

We then examined the laminar patterns of the Xenium-derived dorsal horn neurons labeled as species-conserved neuron subtypes. As shown in **Figure 4D**, each subtype shows a restricted laminar distribution. Exc-MAF cells are in lamina III, whereas Exc-NMUR2 cells reside in lamina II. We hypothesized that with a formal analysis, each labeled subtype in the mouse would show similar superficial or deep dorsal horn distributions consistent with RNAscope analysis in macaque (see **Figures 2B-F, S4-5**). We thus computed a per-cell normalized measure of laminar position relative to other neurons in each of the sixteen dorsal horn slices (z-score). As seen in **Figure 4F**, most neuron subtypes are in either the superficial or deep dorsal horn. Both the macaque RNAscope and mouse Xenium assays show the same significant patterns: Exc-BNC2, Exc-NMUR2, Exc-SKORS2, Exc-NELL2, and Inh-NXPH1 are all superficial; while Exc-MAF, Exc-NTS, Inh-CACNA2D3, and Inh-SORCS1/Inh-PDZD2 are deep. For the remaining subtypes, one of the two methods showed non-significant enrichment, but the trends were consistent with each other. Finally, data obtained with Xenium from a second mouse again showed separable neuron subtypes (**Figure S18A-B, D**), that had similar laminar distributions and PVALB was again enriched in the Inh-CACNA2D3 subtype (**Figure S18C**).

### Exc-PBX3/PDYN is a distinct dorsal horn cell type, while Spp1 neurons previously called MidVent are prevalent in the deep dorsal horn

One unexpected observation from our spatial transcriptomics analysis is that the neuron subtypes were all located within laminae I-III (**Figure 4B)**, We therefore went back to the large population of neurons that were previously not matched and were not clearly distinguishable as discrete clusters, initially assigned as MidVent (**Figure S19A-B**). To further characterize these neurons in the Xenium mouse *in situ* dataset, we performed clustering of the neurons not assigned to the conserved neuron subtypes (**Figure S19C-D**). We integrated them with the MidVent clusters of the adult mouse snRNA-seq. Consistent with our previous analyses, both Xenium clustering and Xenium-snRNA co-clustering yielded poor separation (**Figure S20A**). Label transfer between snRNA-seq and Xenium also failed (not shown; most cells erroneously labeled as motoneurons). Similar to the analysis in **Figure 4F**, we then tested whether these closely related Xenium sub-clusters show specific laminar patterns (**Figure S20B**). From this analysis, we observe that several of these sub-clusters are located dorsal to the central canal. Interestingly, one of these subclusters, xen.9. is found in the superficial dorsal horn and is well-aligned with Excit.25 previously designated in the mouse RNA-seq data (**FigS19C-D**) and Exc-PBX3/PDYN in macaque. It was previously shown to be homologous to Ex-M-2 in humans^17^.

We then further differentiated xen.9 (putative Exc-PBX3/PDYN) and the remaining “MidVent” neurons by finding their top marker genes available in Xenium. We found the top marker for “MidVent” is Spp1 (**Figure 5C**). Because these cells are prevalent not just ventrally but also within the dorsal horn, we re-named them the Spp1 family. Second, we confirmed that Pdyn was a highly specific marker for only sub-cluster xen.9 (**Figure 5C-E**), and not expressed in Spp1 or excitatory dorsal horn subtypes. Given the congruence with prior snRNA-seq findings (**Figure 2**, see also Ex-M-2 expression of PDYN in Yadav *et. al.* Figure 3D), and the prior literature about excitatory prodynorphinergic neurons^40,41^, we named xen.9 the Exc-PBX3/PDYN cell type. A single-cell visualization of all neuron subtypes of the dorsal horn is depicted in (**Figure 5A-B)**.

**Figure 5.**
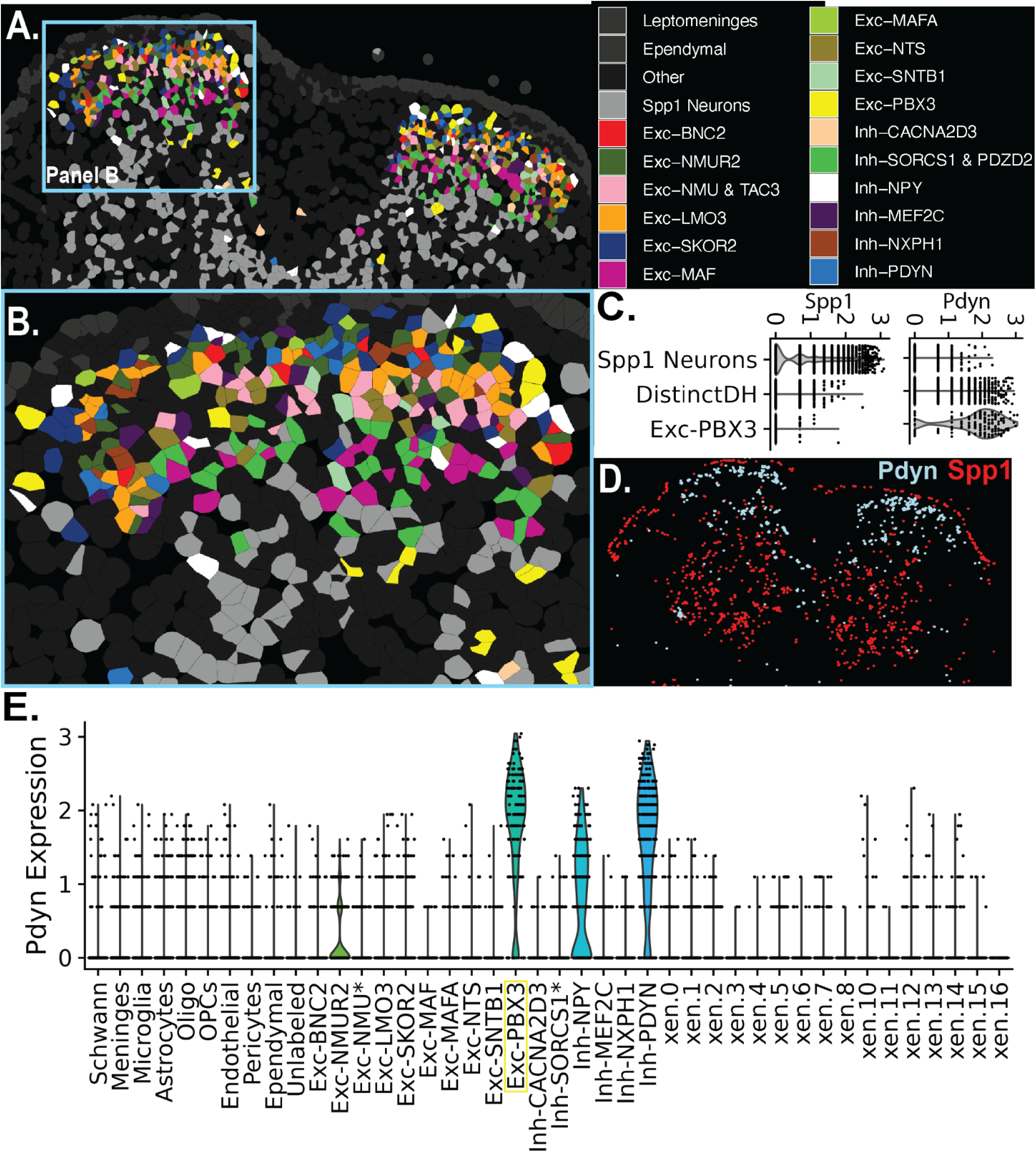
Single-cell spatial visualization of dorsal horn neuron subtypes in mouse. Exc-PBX3 is a superficial dorsal subtype identifiable by Pdyn expression. **A.** Distinct neural subtypes previously labeled based on RNA integration are shown with cell segmentations and individual colors. Leptomeninges and other glia are shown in shades of dark grey to demarcate the meningeal border and white matter of the spinal cord. The Spp1 family of neurons, previously called “MidVent”, are shown in light grey. **B.** Distinct neuron subtypes are shown to be concentrated in the superficial dorsal horn. **C.** Violin plot showing the expression of Spp1 (left) and Pdyn (right) among Spp1 neurons, Exc-PBX3, and distinct neural subtypes besides Exc-PBX3 (DistinctDH). These categories account for all dorsal horn neurons. **D.** Visualization of Spp1 (red) and Pdyn (light blue) transcript distribution in the spinal cord. **E.** Violin plot of Pdyn expression across, glial cell types, distinct dorsal horn subtypes, and subclusters of the Spp1 family (indicated by xen.). In addition to the excitatory marker Slc17a6 (not shown), Pdyn expression distinguishes Exc-PBX3 from all other cell types and subclusters.

### Single-nucleus Open Chromatin of Dorsal Horn Conserved Neuron Subtypes

Cell-type-specific gene expression patterns are driven by the activity of *cis*-regulatory elements such as enhancers. Previous studies of common, polygenic neuronal diseases have found that genetic variants found through genome-wide association studies (GWAS) are often linked to cell-type-specific *cis*-regulatory elements^42–44^. To investigate the relationship between chronic pain and dorsal horn regulatory elements, we harvested single nuclei from lumbar segments L3-5 of two pools of 10 mice each (20 total, 5 Female/5 Male per pool, 7-8 weeks old), and performed single-nucleus assay for transposase-accessible chromatin sequencing (snATAC-seq, **Figure 6A**). Following sequencing, we pre-processed this dataset using the software ArchR^45^, removing nuclei that did not pass metrics for sufficiently high TSS enrichment (3.5) or unique fragments (log10(3.5)); we also removed potential doublets based on ArchR’s simulated doublet method. Overall, the data are of high quality based on several metrics such as TSS enrichment, nucleosome periodicity, number of fragments (**Figures S22-24**), and successful sample-wise batch correction was evident (**Figure S25**). The vast majority of nuclei have a TSS fold enrichment > 10, indicating high experimental quality^46^. We next performed un-biased clustering of cells, and then we applied ATAC-RNA integration, which compares the accessibility at, and surrounding, gene transcription start sites as a proxy for gene expression in snATAC-seq, with the actual gene expression levels measured from nuclei in the snRNA-seq reference dataset. Assigning snATAC-seq nuclei to the most similar snRNA-seq nuclei provided a labeling of neurons versus glial cell classes, which we verified by also visualizing the gene scores for established marker genes of cell classes (**Table S13**). We identified 74,437 glial cells and 19,073 neuronal cells in total (**Figure 6B**). We then sub-clustered the snATAC-seq neurons and again used ATAC-RNA integration to label them with conserved subtype labels. RNA-ATAC co-embedding of neurons is shown in **Figure S26**. Integration yielded distinct, well-separated neuron subtypes, recapitulating the separation seen in snRNA-seq data (**Figure 6C**), with the exception that Exc-PBX3 and Exc-NTS were not found in sufficient numbers for downstream analysis. With the nuclei now labeled, we generated cell-type-specific open chromatin peaks for glia classes and for species-conserved neuron subtypes.

**Figure 6.**
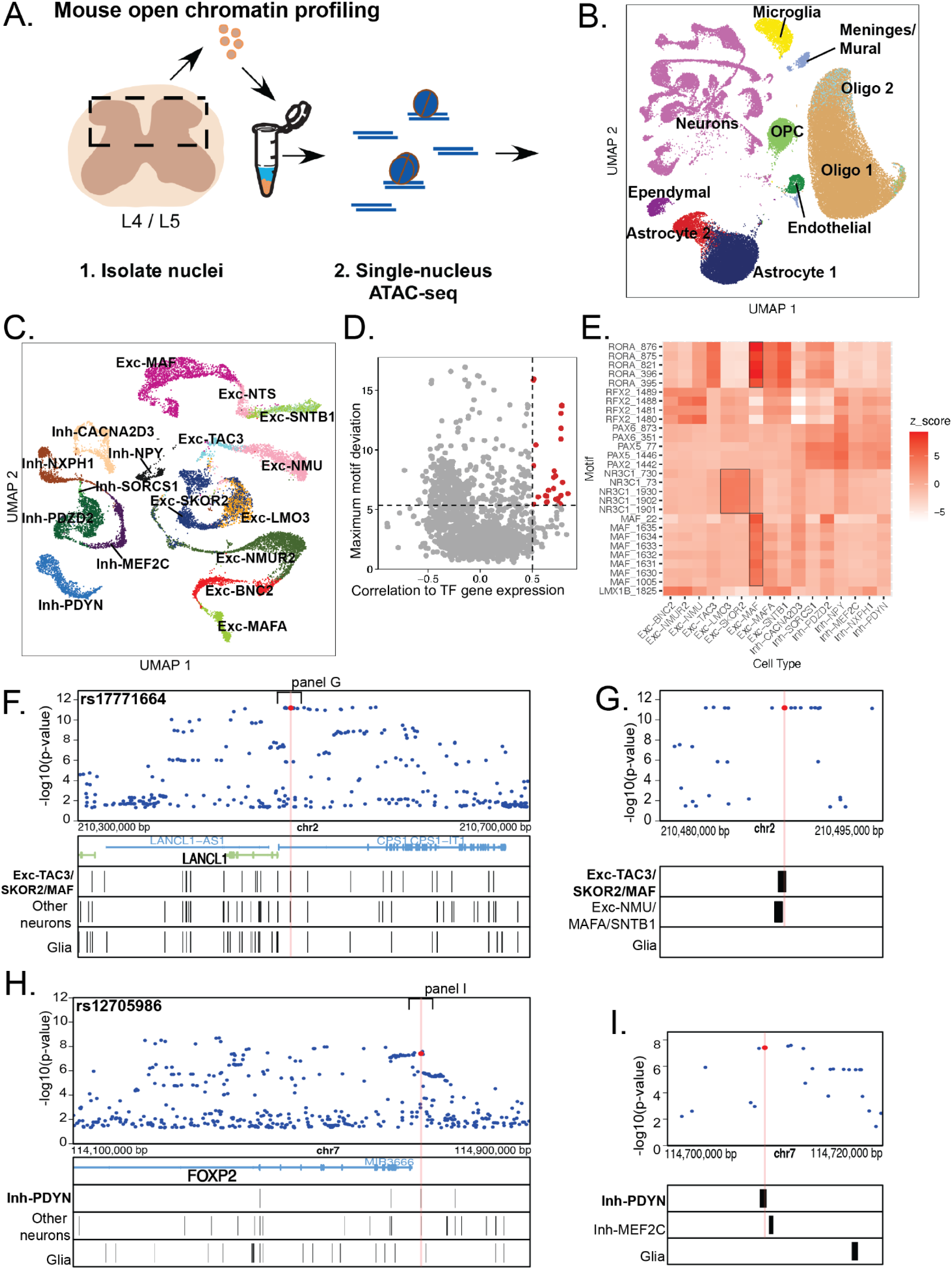
Mouse snATAC-seq open chromatin: Dorsal horn neuron neuron subtype identification, transcription factor motifs, and candidate regulatory SNPs. **A,** Schematic overview of the single-nucleus ATAC-seq experimental workflow of the mouse lumbar dorsal horn (L4/5). **B.** UMAP representation of snATAC nuclei and major cell types after label transfer using RNA-ATAC integration and confirmation using gene scores of marker genes. **C.** UMAP representation of dorsal horn neuron subtype nuclei after neuron-specific re-clustering and label transfer using RNA-ATAC integration. Cell subtype color scheme same as Figure 3**,4,5. D.** Identification of positive transcription factor (TF) regulators. Motifs are considered TF regulators if they satisfy (a) in the top quartile of motif deviation, a measure of cell-type specificity (y-axis), and (b) their presence is correlated (>0.5, Pearson correlation) with the imputed gene expression of their corresponding TF (x-axis). **E.** Heatmap showing the normalized motif deviation (z-score) of each TF regulator by neuron subtype. Black boxes highlight the MAF and RORA motifs specific to Exc-MAF, and NR3C1 motifs (glucocorticoid receptor) specific to Exc-LMO3 and Exc-SKOR2. TF = transcription factor. **F.-I.** Example GWAS SNPs near genes linked to the spinal cord and chronic pain within subtype-specific mouse open chromatin peaks. Top section of each panel are Manhattan plots of significant SNPs (red dot, with rsID above) and nearby SNPs (blue dots) from Kupari *et al*. The bottom half of each panel are the following annotations from top to bottom (1) the nearby genes (Gene), (2) the open chromatin peaks of the highlighted neuron subtype, and (3) peaks of other dorsal horn subtypes (Other). The translucent red rectangle highlights the SNP falling within the peak of interest. **F.-G.** A LANCL1 SNP falling within Exc-TAC3/SKOR2/MAF peaks. **H.-I.** A FOXP2 SNP falling within an Inh-Pdyn peak. SNP - single-nucleotide polymorphism. eQTL = expression quantitative trait locus.chr2,7 = chromosome 2,7. KB = kilobases range of locus.

Chromatin accessibility and cis-regulatory activity are driven by the binding of transcription factors (TFs) with specific affinity for unique combinations of DNA motifs. Previous snRNA-seq studies found that dorsal horn neuron subtypes express characteristic TFs^17^, but were not able to connect them to their motifs without open chromatin data. Given our joint ATAC-RNA profiles of dorsal horn subtypes, we investigated whether TFs are specific in both gene expression and motif enrichment among the dorsal horn subtypes. We measured motif enrichment using the chromVAR software within ArchR. Higher deviation scores are a measure of higher subtype specificity of motif appearance. Given that families of TFs have similar motifs, it can be difficult to robustly link specific TF genes with distinct motifs. We thus calculated the correlation between gene expression of TFs (from the RNA integration) with their putative motif deviations; a higher correlation indicates a higher likelihood of a true biological correspondence between a TF-motif pair. We defined positive TFs regulators as having a maximal motif deviation above the 75th percentile (higher specificity) as well as correlation > 0.5 of per-nucleus motif enrichment with expression level of the TF gene (**Figure 6D**). As shown in **Figure 6E**, we found expected regulators such as MAF and RORA for the Exc-MAF cells. We also found a novel potential regulator, NR3C1 (the glucocorticoid receptor) for Exc-LMO3 and Exc-SKOR2. Given the limitations of RNA expression imputed via RNA-ATAC integration, we also identified candidates with very high deviations (above 95th percentile but with a relaxed gene correlation condition of > 0, **Figure S27**) to nominate candidate motifs for further investigation as potential drivers of neuron identity. For example, we found that ASCL2, MYOD1, and several TAL1 motifs were specific to Inh-CACNA2D3.

### Genetic risk variants for chronic pain are enriched in mouse neuron subtype-specific open chromatin

Recent chronic pain GWAS have revealed several candidate genetic risk loci for general chronic pain as well as localized conditions: head, face, neck and shoulder, back, hip, stomach and abdominal, and knee.

For other common polygenic diseases, risk variants tend to fall within non-coding regions of the genome, and we hypothesized this is the case for chronic pain. As an initial estimate, we calculated the proportion of genome-wide significant SNPS falling within coding or non-coding regions. We found that 15 of 389 (4%, Kupari *et. al.*) and 248 of 4,094 (6%, Khoury *et. al.*) were in exons, while the remaining 96% and 94%, respectively, of variants fell within non-coding regions.

We then sought out cases when potentially causal SNPs from the chronic pain GWAS overlapped with subtype-specific open chromatin. First, we mapped GWAS SNP and open chromatin coordinates to the same human coordinates (hg38)^47,48^ (56% successfully mapped). We limited our search to the most significant loci from the GWAS, in particular those belonging to a credible set of potentially causal SNPs based on Bayesian fine-mapping estimates (LocusZoom^49^). We found notable SNPs near two genes of interest, LANCL1 and FOXP2 (**Figure 6F-I**). On human chromosome 2, we found the SNP rs17771664 (an expression quantitative trait locus for LANCL1), from ^18^, “Number of Pain Sites”, which fell within a peak shared by Exc-TAC3, Exc-SKOR2, and Exc-MAF (**Figure 6F)**. We observed that this SNP is among the most significant of its LD block (**Figure S28A**) and belongs to the credible set of LocusZoom SNPs that were statistically calculated as likely causal (**Figure S28B**). As shown in **Figure 6G**, a peak belonging to Exc-NMU/MAFA/SNTB1 is also nearby this SNP of interest. Similarly, on human chromosome 7, we found the SNP rs12705986 nearby the gene FOXP2, also satisfying our criteria for LD and belonging to the fine-mapping credible set (**Figure S28C-D**) and falling within a peak specific to Inh-PDYN (**Figure 7H)**. As seen in **Figure 7I**, a peak specific to Inh-MEF2C is also nearby this SNP. These SNPs are only a couple examples of the many that exist within this dataset. Because these data show links to open chromatin of specific dorsal horn neuron subtypes, we hypothesized that our open chromatin dataset is enriched for chronic pain variants on a larger scale, and thus establishes a link between chronic pain genetics and the spinal cord that has previously been missed in studies focused on marker genes.

**Figure 7.**
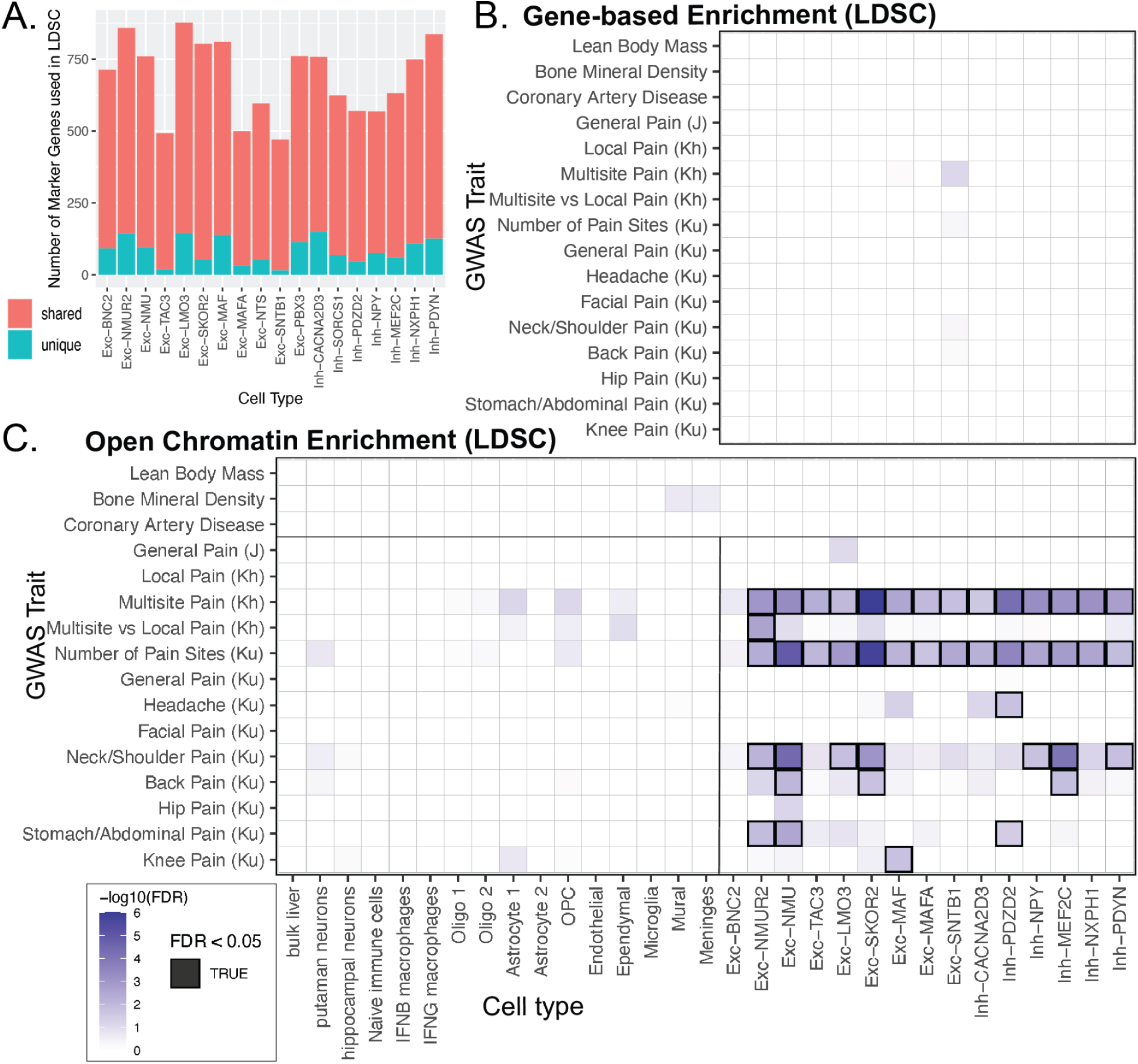
Open chromatin profiles rather than marker genes of distinct dorsal horn neuron subtypes are enriched for human SNP markers of several chronic pain disorders. **A.** Bar plot showing the proportion of marker genes for each neuron subtype that are either strictly specific to that neuron subtype (unique, blue) or a marker gene for at least one other neuron subtype (shared, red). **B.** Gene-based stratified LDSC based on marker gene enrichment only. The intronic and flanking non-coding regions of each neuron subtype (x-axis of **panel C**) were aggregated. The background for all subtypes was the intronic and flanking regions of all coding genes. Enrichment of GWAS SNPs per cell type versus the background was estimated by stratified LDSC. Per legend (shared by **Panel C**, lower left), darker purple indicates greater enrichment for the particular trait (y-axis) and neuron subtype (x-axis). Top three traits were non-neuronal negative controls. Significance based on FDR <0.05 is indicated by a black box around the square. **C.** Stratified LDSC of subtype-specific open chromatin. Foregrounds for each neuron subtype were the specific reproducible peaks from snATAC-seq. The background for each subtype was the union of foregrounds merged with a large mouse DNase hypersensitivity site dataset (see Methods). Legend and interpretation same as **panel B**. In addition to neuron subtypes, enrichment was estimated for glial cell types from the same dataset, as well as bulk liver, putamen and hippocampal brain neurons, and naïve, IFN-B, and IFN-G immune cells from separate datasets (details in Methods). J= Johnson et al, Kh = al Khoury et al, Ku = Kupari et al. LDSC = Linkage-disequilibrium score regression. FDR = false discovery rate. GWAS = genome-wide association study.

First, as a comparison, we investigated if variants are associated with non-coding regions nearby marker genes (**Figure 7B**, *Gene-Based Enrichment*). Taking the marker genes previously established for each cell type (**Tables S5-7**), we first observed that most marker genes are not strictly specific (blue bars) to a single neuron subtype (**Figure 7A**, blue bars), but rather most are shared (red bars). We next identified the intronic and flanking regions adjacent to each of these genes, in order to potentially capture regulatory elements. To estimate the enrichment of significant SNPs from GWAS, we employed a stratified linkage disequilibrium score regression (s-LDSC) approach (see Statistical analyses)^43^. We performed s-LDSC analysis for chronic pain traits from three major GWAS studies as well as for three negative control traits (lean body mass, bone mineral density, and coronary artery disease)^18,26,50^. After correcting for false discovery rate, we found that none of the cell types were significantly enriched for any of the traits (**Figure 7B**). We hypothesized that this result reflects a limitation of our approach rather than a true biological non-association, partly on marker gene observations (**Figure 7A**) and constructing windows around genes is not a direct way of capturing regulatory elements, especially distal enhancers. We therefore hypothesized that our (human orthologs of) direct measurements of single-nucleus open chromatin could be functionally relevant to chronic pain and performed stratified LDSC using the open chromatin annotations (**Figure 7C**, *Open Chromatin Enrichment*).

As comparators to the dorsal horn neurons and glia, we also analyzed open chromatin from bulk liver ATAC-seq^51^, neurons from the hippocampus and putamen^52^, and macrophages^53^, the latter given the potential immunogenic origin of some chronic pain disorders. As shown in **Figure 7C**, none of the negative control traits, or traits in non-dorsal horn cells or in glial cell types show enrichment.

In contrast, among the dorsal horn neurons, all but one subtype (Exc-BNC2) are enriched for at least one chronic pain trait. The traits with the most associations are multi-site pain and higher number of pain sites. Several cell types (Exc-NMU, Exc-NMUR2, Exc-SKOR2, Inh-PDZD2, Inh-MEF2C) show associations with at least four traits. Differences in trait heritability or number of marginally significant SNPs for each trait are provided as potential factors for why some traits tend to be more enriched (**Table S14**).

## Discussion

Understanding the molecular and spatial organization of the dorsal spinal cord is essential for defining functional circuits, determining how incoming somatosensory information is integrated and transformed, as well as for understanding the origins of chronic pain and how to treat it. In this study, we generated a framework for the development of chronic pain, in which genetic variants for chronic pain lie within regulatory elements of specific dorsal horn neuron subtypes. This finding is consistent with previous work showing dorsal horn subtypes have a causal role in persistent pain^34,54,55^. The implication of our findings here is that these variants work separately or together to produce functional changes in how these neuron subtypes respond to injury, predisposing people to the development of chronic pain. This model of altered functioning of regulatory regions of dorsal horn subtypes is consistent with what has been reported for other diseases with a genetic component; in fact, 80-90% of complex disease-associated genetic variants disrupt non-coding regulatory regions, primarily enhancers^21^. Thus, in this work, we provide insight into the predisposition for chronic pain that lies at the intersection of human genetics, regulatory genomics, and cell-type-specificity, as a critical step for understanding the cellular mechanisms of how genetics influences chronic pain disposition. In future work, alterations in gene regulatory networks within these subtypes can be studied to identify dorsal horn mechanisms underlying chronic pain.

Our comparative analyses between macaque, human, and mouse show that the underlying molecular and spatial organization is largely conserved with snRNA-seq integration demonstrating a high degree of consistency among cell types and across species (**Figures 3-4, S8-9**); although the number of defined clusters differed across studies, specific sets of mouse and human clusters reliably mapped to the macaque clusters with few exceptions, namely Exc-TAC3 and Exc-PBX3. In macaque, the Exc-TAC3 is a rare population with high similarity to the Exc-NMU subtype. Because we defined the macaque clusters more conservatively than prior spinal cord atlases (**Figure 1**), the difference in cell proportions observed between species may reflect experimental differences or a true divergence between the Exc-TAC3 and Exc-NMU subtypes in macaques.

Our analysis using single cell spatial transcriptomics showed for the first time the precise location of the single cell RNA-seq defined neuron subtypes. Surprisingly, the most molecularly distinct subtypes are all restricted to the superficial laminae I-II and lamina III while the much less distinct clusters, that we initially defined as MidVent (now named SPP1), are in laminae IV-V and into the ventral cord. This finding, while surprising, is consistent with the functional roles of these regions of the dorsal horn, with the superficial laminae responsible for exquisite processing of many different modalities: cold, heat, itch, touch and pain, while the more ventral laminae process information related to touch and sensorimotor transformations^56^. The less distinct molecular composition of neurons in the deep dorsal horn may also apply to the many populations of projection neurons that reside there^57^. Intriguingly, many SPP1 sub-clusters express Tacr1, which is associated with projection populations^58^. However, these are relatively rare populations and thus are not resolvable in the current work. Future studies can take advantage of our newly provided datasets, coupled with retrograde labeling and spatial transcriptomics, to molecularly and anatomically characterize these important rare populations in both rodents and primates.

Our findings also resolve discrepancies for the PBX3/PDYN subtype reported among prior cell atlases. In previous studies, the mouse and human orthologs of Exc-PBX3/PDYN were assigned to “MidVent’’ based on transcriptomic similarity to other neurons in that region of the cord, consistent with what we observed for macaque. However, in mice, the subtype had been assigned by RNAscope to layers IV-VI (Russ et al), while in humans, using Visium (a lower resolution spatial platform than Xenium), it appears to be more superficial (Yadav), as we observed here. We thus establish in this study that Exc-PBX3/PDYN contain the excitatory prodynorphinergic neurons of the superficial dorsal horn^40,41^, a finding consistent with other published work using two probe *in situ* hybridizations in mouse^55^ and with our own RNAscope analysis in macaque and human (**Figures S5-6**).

The genetics of chronic pain has previously been studied in relation to marker genes within the spinal cord, but to our knowledge not in relation to the more direct measure of regulatory elements. We thus developed a single-nucleus open chromatin resource for the dorsal horn to investigate dorsal horn regulatory genomics and its relationship to chronic pain. First, we find that chromatin accessibility patterns are signatures of cellular identity (**Figures 6C, S26**), consistent with the established biology that cell-type specializations are determined by gene regulatory elements. We also identified several transcription factor regulators of particular cell types, including those previously suspected based on gene expression alone (*e.g.* MAF and RORA for Exc-MAF) and on novel candidate regulators (*e.g.* TAL1 and NR3C1). Motifs for NR3C1, the glucocorticoid receptor, were specifically abundant in the open chromatin of Exc-LMO3/Exc-SKOR2, two closely related cell types in the mouse Sox5 family. NR3C1 is the well-known target of cortisol, which is released during periods of chronic stress as part of the hypothalamic-pituitary-adrenal (HPA) axis^59^.

Several studies link NR3C1 expression to spinal modulation of pain^60–64^. Our findings suggest that Exc-LMO3/Exc-SKOR2 could mediate the effects of chronic stress on pain phenotypes, whereby NR3C1 activation by cortisol release leads to changes in Exc-LMO3/Exc-SKOR2 gene expression and potentially neural firing, although experimental follow-up is needed to confirm these hypotheses-generating results.

By analyzing accessible chromatin rather than genes, our work establishes the spinal cord as a tissue harboring significant genetic risk for chronic pain. Our combination of genomic analyses shows that unlike glia and comparative tissues such as macrophages, open chromatin of several dorsal horn neuron subtypes are significantly enriched for variants of chronic pain traits (**Figure 7C**), which our gene-based analysis did not capture (**Figure 7B**). Prior eQTL-based analysis of chronic pain GWAS also did not find enrichment in spinal cord^26^, showing that our open chromatin resource fills a critical gap in the characterization of tissue-specific mechanisms of chronic pain risk, and links risk to specific neuronal subtypes. Although the ability to find human phenotype enrichment in orthologs of mouse open chromatin may be surprising, tissue- and cell-type-specific transcription factor binding sites and open chromatin regions are often functionally conserved, even when individual nucleotide conservation is not^51,65,66^. This degree of conservation explains how complex trait-associated genetic variants can show similar enrichment in human cell type-specific open chromatin regions and orthologous open chromatin regions in mouse^67^.

We note the existence of neuron-subtype enrichment depending on the anatomical location of pain; for example, SNPs from knee pain, a type of pain that likely has a strong inflammatory component, were strongly associated with Exc-MAF. This finding is consistent with experiments showing neurons in the Exc-MAF subtype are critical for conveying mechanical hypersensitivity in mouse models of inflammatory pain^34^. Neuron subtypes for stomach/abdominal pain, which are visceral, were strongly associated with other populations, including Exc-NMUR2, Exc-NMU and Inh-PDZD2. Studies of visceral pain circuitry in mice have not yet identified which dorsal horn neurons are important but these subtypes which overlap with somatostatin, PKCγ and PVALB neurons have been implicated in other types of persistent pain^34,54,55^. Headache SNPs are also enriched in open chromatin of Inh-PDZD2. This latter finding could be due to the innervation of upper cervical regions of the spinal cord by cranial muscles and dura or because of functional and anatomical similarities between spinal and medullary dorsal horns. Nevertheless, differences in the neuron subtypes that show strong associations for each type of pain reported is consistent with studies showing that chronic pain mechanisms and circuits differ by the nature of the injury^34,68,69^. It also fits that the number of neuron subtypes we identified as being highly enriched in open chromatin SNPs from patients with head/neck and multisite pain, which are likely to be combinations of inflammatory and neuropathic etiologies, was much greater than for the other types of pain. This finding is also consistent with the large number of dorsal horn cell populations that have been implicated in persistent pain in rodent functional studies, including neuron subtypes implicated here^70,71^. Again, these results suggest further studies are warranted to understand precisely how the specific neuron subtypes are involved in these types of chronic pain.

We also took note of two genes, LANCL1 and FOXP2, that are nearby regulatory regions with highly significant SNPs, reported in the Kupari et al study. LANCL1 is a neuron-enriched antioxidant that is upregulated with neural activity, neurotrophic factors and the stress response^72^. It is thought to play an important role in suppressing reactive oxygen species^72^. Reactive oxygen species have been implicated in central sensitization mechanisms underlying both neuropathic and inflammatory pain and ROS scavengers have been put forth as potential treatments^73^. Our results point to Exc-TAC3, SKOR2 and MAF as being particularly salient. The second gene, FOXP2 is a transcription factor that has been implicated in development of V1 interneurons in the ventral horn^74^, but is also expressed in both excitatory and inhibitory dorsal horn neuronal subtypes across species. Its role in the dorsal horn and potential role in chronic pain, particularly with respect to the Inh-PDYN subtype, which is known to be important for persistent mechanical hypersensitivity in inflammatory and neuropathic conditions^55^, remains to be explored.

Our study and resources have some limitations. Limitations in clustering strategies have already been discussed above. Our analysis was focused on the dorsal horn, especially the distinct neuron subtypes that reside in laminae I-III, an area critical for acute and persistent forms of pain. To encapsulate the Spp1 family, the spatial transcriptomic analysis was extended beyond the dorsal horn, but ventral populations such as motoneurons^73^ which are beyond the scope of this paper, were not analyzed. Furthermore, there were some limitations of the Xenium spatial transcriptomics assay, as not all conserved marker genes are in the available pre-designed mouse brain panel. We do nevertheless provide strong evidence for reliable localization of the neuron subtypes, including analyzing the available conserved marker genes. There is also a limitation to the snATAC-seq and s-LDSC approach taken because precisely identifying tissue- and cell-type-specific enhancers is challenging. We used distal peaks from snATAC-seq as a proxy for enhancer activity. Also, peaks shared across non-disjoint subsets of neural subtypes may partially confound subtype-specific findings. But limiting choices to peaks found in only one cell type would lead to uncontrolled type I error^75^. Finally, the brain regions and immune cell databases analyzed here did not show enrichments for chronic pain variants, but future studies may identify other tissues and cell types with heritable contributions to chronic pain.

Overall, our findings demonstrate that this mouse open chromatin resource, in conjunction with the conserved cell type framework, can be used to generate novel, testable hypotheses about the origin and pathophysiology of chronic pain. Having anchored the relationships between cell identity, genetic risk, regulatory genomics, and pathophysiology in humans and in the animal models typically studied in biomedical research, we help provide the foundation for studying the complex genetics and neural mechanisms underlying chronic pain disorders, and for developing a new generation of targeted therapeutics. There are now several FDA approved gene therapies including for inherited blindness and for spinal muscular atrophy^76–78^, demonstrating the potential safety and efficacy of these strategies. The development of single cell genomics and epigenomics enables further advances in gene therapy, namely tissue- and cell-type-specific targeting, which has complementary applications both for mechanistic studies of neural circuits, and for gene therapy. Targeting dorsal horn subtypes using promoters has shown some success^79,80^, but a particularly compelling strategy is to drive expression of therapeutic transgenes using cell-type-specific enhancers. Previous work has shown the feasibility of this approach in the brain^81–84^, with most successful approaches being those based on enhancer identification from chromatin accessibility^81–83^. Although our snATAC-seq dataset is from mice, extensive research shows that many cell-type-specific enhancers are robustly conserved across species^51,65^, and candidate enhancers identified in mouse would have therapeutic potential for humans. Thus, we are providing the spinal biology community with key resources to make progress on cell-type-specific targeting strategies for disorders such as chronic pain.

## Acknowledgements

The authors thank Ian Cumming at the Duke Human Vaccine Institute Flow Cytometry Core Facility, Nicolas Devos of the Duke Sequencing and Genomic Technologies Shared Resource and Dr. Priyabrata Halder for help with spinal cord harvests. We thank Ryan Patterson and Dr. Ariel Levine for access to and early discussions of the published mouse snRNA.seq dataset. Additionally, we thank Michael Pavel Vannice, Samyuktha Lokanandi and Andrea L. Goliaszewski for help with *in situ* hybridization quantification, Suh Jin Lee for the spinal cord diagram in Figure 1, and Isabel Bleimester as well as Monica Dayao for comments on the manuscript. We also thank the organ donors and their families for the donation of spinal cord for research purposes and Dr. Jeffrey Reese and Anna Cervantes for spinal cord recovery. We thank Dr. Janette Lamb and the University of Pittsburgh Genetics and Genomics Core for generation of single cell ATAC libraries and the UPMC Genomics Core for single cell library sequencing. This research was supported in part by the University of Pittsburgh Center for Research Computing through the resources provided. Funding was provided by NIH extramural support to R.P.S. (NS104964, NS133364, and NS109792), C.M.A. (NS111791), D.A.L (MH051234), T.J.P. (NS111929) and A.R.P (NS133364) and NIH Intramural support to A.J.L. (1ZIANS003153).

## Contributions

C.M.A. and R.P.S. conceived of the study. M.J.L., M.K, B.N.P., V.C, R.P., S.L., D.Y, M.F, B.K, M.M.K.N, V.K.C., Z.C. and A.R.P. performed the bioinformatic analyses and data curation. A.C and C.M.L. performed nuclei isolation and libraries for macaque snRNA-seq. B.C.L. harvested nuclei for mouse snATAC-seq.

C.M.A performed the macaque spinal cord *in situ* hybridization and S.S. performed the human spinal cord *in situ* hybridization. M.C.N. and K.A.C. performed the mouse dorsal horn Xenium experiment. C.M.A. created the R shiny macaque dorsal horn web browser. D.A.L. provided the Rhesus macaques and R.D. harvested macaque tissue. C.M.A, M.J.L, A.R.P and R.P.S. statistically analyzed the data. R.P.S, A.R.P.

T.J.P and Y.Q. provided resources and intellectual input. R.P.S, C.M.A, and M.J.L wrote the manuscript with significant contributions from M.K., B.N.P., A.R.P., and A.C.

## Resource Availability Lead Contact

Further information and requests for resources and reagents should be directed to and will be fulfilled by the Lead Contact, Rebecca P. Seal.

## Materials Availability

This study did not generate any new materials.

## Data and Code Availability

The code and Jupyter notebooks used for the analysis of the macaque snRNA-seq has been made available at Github repository https://github.com/pfenninglab/dorsal_horn_snrnaseq. The code and notebooks for the analysis of three-species integration, conserved marker identification and decision trees, xenium analysis, and mouse snATAC-seq analysis are available at https://github.com/pfenninglab/dorsalhorn_conserved_rna_atac_xen.

The raw and processed datasets for macaque snRNA-seq, mouse Xenium, and mouse snATAC are available through the GEO SuperSeries (GSE253954).

Additionally, the macaque dataset can be accessed through an R shiny application (https://seallab.shinyapps.io/macaquedh/). This Shiny application was created using a Shiny Cell package^81^.

## Methods

### Macaque snRNA-seq Samples

Lumbar spinal cord samples for snRNA-seq were obtained from *Macaca mulatta* provided by Dr. David Lewis at the University of Pittsburgh. Monkeys were housed in groups in the same social setting. All animals were deeply anesthetized with ketamine and pentobarbital and perfused transcardially with ice-cold artificial cerebrospinal fluid. All housing and experimental procedures were conducted in accordance with the guidelines of the US Department of Agriculture and the National Institutes of Health Guide for the Care and Use of Laboratory Animals and with the approval of the University of Pittsburgh Institutional Animal Care and Use Committee. No prior manipulations to the spinal cord were performed in these macaques. Three male macaques (3-years old) were used for the single nuclear RNA-sequencing. Male and female macaque lumbar spinal cord samples (3-5 years old) were utilized for in situ hybridization studies. Though we did not observe sex differences for any of the genes using in situ hybridization, there is not enough statistical power to confidently assess if there are sex differences due to the limited availability of macaque tissue.

### Human Spinal Cord Samples

All human spinal cord procurement procedures were approved by the Institutional Review Boards at the University of Texas at Dallas. Donor information is provided in **Supplementary Table 5**. The human spinal cords were gradually embedded in OCT in a cryomold by adding small volumes of OCT over dry ice to avoid thawing. All tissues were sectioned at 20 µm onto SuperFrost Plus charged slides using a cryostat. Sections were only briefly thawed in order to adhere to the slide but were immediately returned to the -20°C cryostat chamber until completion of sectioning. Due to the limited availability of human donor tissue, sex differences at the gene level were not assessed.

### Nuclear Dissociation and Isolation

Snap-frozen lumbar spinal cord segments were removed from -80°C storage and placed into separate Petri dishes containing a cold slurry of dissection buffer consisting of 1x Phosphate Buffered Saline (PBS, ThermoFisher Scientific; AM9625), 10% Dithiothreitol (DTT, Sigma; 43816-50ML).

From each animal, transverse sections (∼50-75 mg) were made using a sterile razor blade. Three sections were selected for further dissection and the rest were discarded. The sections were cut in the coronal plane at the middle of the spinal cord to obtain dorsal and ventral halves. Any residual meningeal membranes were removed. The three dorsal halves were retained and the ventral halves discarded.

Nuclei were isolated according to Martelotto with modifications^73^. The dorsal halves for each animal were transferred into separate Dounce homogenizers (Sigma) containing 1 mL of ice-cold homogenization buffer consisting of EZ Nuclei Lysis Buffer (Sigma; NUC101-1KT) with 0.5% RNasin Plus (Promega; N2615), 0.5% SUPERase-In (ThermoFisher; AM2696) and 1mM DTT. The samples were homogenized on ice using 20 strokes of Pestle A followed by 20 strokes of Pestle B. Any residual meningeal membrane was removed before switching pestles. The homogenate was filtered through a 50 µm filter (Sysmex; 04-004-2327) into a 2 mL microcentrifuge tube (Eppendorf; 022431048). An additional 0.5 mL of homogenization buffer was used to wash the Dounce homogenizer and filter. The sample was then placed on ice while the remaining samples were processed. The sample was centrifuged at 500g at 4°C for 5 min to obtain a crude pellet containing spinal nuclei. The supernatant was removed and discarded, being careful to not disturb the pellet. The pellet was resuspended in 1.5 mL of Homogenization Buffer and allowed to sit on ice for 5 mins. The samples were again centrifuged at 500x g, 4°C for 5 min. The supernatant was removed and the pellet was resuspended in 1 mL of Nuclei Resuspension Buffer (NRB) consisting of 1x PBS, 1% Bovine Serum Albumin (BSA, Sigma; 2905-5GM) and 1% SUPERas-In followed by centrifugation at 500g, 4°C for 5 mins. This wash step was repeated twice more for a total of 3 washes. The final pellet was resuspended in 0.5 mL of NRB containing 6 µM 4′,6-diamidino-2-phenylindole, dihydrochloride (DAPI, ThermoFisher; D1306). The suspension was filtered through a 20 µm filter (Sysmex; 04-004-2325) into a polypropylene tube and kept on ice.

Fluorescence Activated Nuclear Sorting (FANS) was performed to purify nuclei from debris on a FACSAria II (BD). Gates were set to isolate DAPI+ singlet nuclei based on forward scatter and side scatter as well as fluorescence intensity. The instrument was set to 40 pounds per square inch (psi) of pressure and a 70 *μm* nozzle was used, with sterile PBS sheath fluid. Nuclei were sorted into a 1.5 ml microcentrifuge tube containing 15 *μl* of NRB at 4°C. For each sample, 18,000 events were sorted into the collection tube. The sorted nuclei and NRB total volume were approximately 45 µl, allowing for loading the entire suspension into the Chromium Single Cell 3’ v3 solution (10x Genomics) without further manipulation. The 10x libraries were processed according to the manufacturer’s instructions. Completed libraries were run on the Novaseq 6000 (Illumina).

### Macaque snRNA-seq Alignment

Macaque snRNA-seq: Illumina sequencing bcl files were converted to fastq format using the *cellranger mkfastq* command line tool. Fastq reads were then aligned to the rheMac10 reference genome and quantified into a raw UMI count matrix using the STAR aligner with the -solo option^74^ against a custom transcriptome annotation that we previously developed was also given as input to the STAR aligner^22^.

### Macaque snRNA-seq droplet Filtering

Macaque snRNA-seq: Each biological replicate underwent a separate but standardized quality control (qc) filtering, whereby empty droplets were detected using the *defaultDrops()* function, which is a quantile filtering approach based on number of UMIs, with default parameters in the DropletUtils package^75^. Doublet filtering was also performed using the *cxds_bcds_hybrid()* function, which uses a hybrid approach utilizing artificial generated doublets and unlikely coexpression pairs to create a doublet score in the SCDS package^76^. Droplets with doublet scores above 1.0 were removed. After both rounds of filtering, 12,243 cells were left across 3 replicates. Both functions were run with default parameters.

### Macaque snRNA-seq Normalization

Macaque snRNA-seq: The filtered digital gene expression matrices were then loaded into Scanpy to perform the pre-clustering needed for Scran normalization^23,25^. Each biological replicate expression count matrix was then normalized with log counts per million (CPM), and denoised using iterative pca with 50 components. The pre-clustering was done via the Leiden community detection algorithm using Scanpy’s *leiden()* function^77^. All 3 biological replicates were then co-normalized using the *computeSumFactors()* function in Scran and *normalize()* function in Scuttle with the pre-clusters as input^78^. The size factor normalization that was used was chosen for its ability to handle sparse single cell data which may have cell types with different expression values.

### Unbiased Clustering of Macaque snRNA-seq

Post-normalization clustering was performed again using Scanpy’s Leiden function^77^. The clusters were classified into oligodendrocytes 1 and oligodendrocytes 2, neurons, astrocyte 1 and astrocyte 2, oligodendrocyte precursor cells, microglia, meninges, ependymal cells and Schwann cells based on the expression of established marker genes (shown in Figure 1C). Neuronal clusters were then manually selected using the marker gene *RBFOX3*, which left a total of 2,698 cells across replicates. Data integration across biological replicates and neuron filtered gene expression matrices were performed using the *integrate_scanpy()* function in Scanorama, which uses a panoramic stitching algorithm to integrate datasets by producing a batch corrected cell-cell distance graph. Leiden community detection was then performed with a resolution of 1.0 on the batch corrected cell-cell distance graph to get an initial integrated clustering^77^. It is important to note that while the Scanorama batch corrected data was used for clustering and integration, the ‘un-corrected’ data was used for marker gene and differential expression analysis to reduce possible bias added during batch correction. Our initial clustering was organized into a dendrogram using Ward hierarchical clustering. Further refinement of these clusters was performed by splitting clusters that had both excitatory and inhibitory markers (*SLC17A6* and *SLC32A1,* respectively). We computed a mid-ventral and dorsal score for each cluster based on the average z-score scaled expression of genes that have a high discriminative ability (AUROC) between the spinal cord regions as computed by previous studies^16^. Two of the macaque clusters were thus grouped into a single cluster labeled MidVentral. This cluster is excluded from our analysis of the dorsal horn populations as it lies outside our region of interest. The list of dorsal horn clusters was initially 11 excitatory clusters and 5 inhibitory clusters, and then extended to 7 inhibitory clusters. Inh-SORCS1/Inh-PDZD2 and Inh-MEF2C/Inh-NXPH1 were split afterwards based on discrepancies between mouse and macaque (see **Methods: Comparison of Macaque and Mouse Expression**).

### Selecting Macaque-specific snRNA-seq Marker Genes

A combination of methods was used for the marker gene selection for the clusters. A binary matrix was constructed by thresholding the normalized gene expression values at a threshold of 0.2 CPM. The binary matrix was then used to compute the precision and sensitivity of each gene for each cluster. In addition, the top differentially expressed genes were computed using the *rank_gene_groups()* function in Scanpy, and Wilcoxon rank-sum (Mann-Whitney-U) test. From this list, we removed genes that were co-expressed in non-neuronal cells.

### Comparison of Macaque and Mouse snRNA-seq Expression Patterns

To identify orthologous genes across species, each mouse gene symbol was matched to the corresponding ENSEMBL ID using BioMart (v101). Then, BioMart was used to identify the orthologous human gene, filtering for one-to-one orthologs to avoid potential false positive matches between species. The macaque genome was annotated with orthologous human genes as previously described, motivated by previous successful approaches to annotate more complete gene structures using orthology to human^22,79^. The resulting datasets frame for macaque and mouse contained 15,157 one-to-one orthologs.

Orthologous genes of two of the mouse family markers, *RORB* and *ADAMTS5*, showed separation in the macaque UMAP, but were not independently clustered. Inh-SORCS1/Inh-PDZD2 and Inh-MEF2C/Inh-NXPH1 clusters were re-clustered using Leiden clustering (resolution 0.3)^77^. Three of the identified clusters represented *RORB* populations while the remaining two populations represented *ADAMTS5*. We thus split Inh-SORCS1/Inh-PDZD2 and Inh-MEF2C/Inh-NXPH1 clusters into two sub-clusters each, with Inh-SORCS1 and Inh-MEF2C representing the RORB population and Inh-PDZD2 and Inh-NXPH1 representing ADAMTS5 populations.

We identified the top 100 most enriched and most-depleted makers for each of the mouse cell classes (Excitatory, Inhibitory, and MidVentral). Mouse and macaque markers for comparison were selected using the Wilcoxon rank sum approach in the *rank_genes_groups()* function^25^. The relative levels of those markers, weighted by their cluster enrichment scores, created a score that we could use to annotate each cell in the macaque population for each of the major cell classes. These enrichment scores corresponded strongly with the individual marker-based annotation of the macaque cell classes, and were used to identify significant shifts across the population using a t-test. The t-statistic confirms the broad differences in the distribution of the enrichment scores, with the exception of cluster Exc-PBX3, which has features of MidVentral neurons.

The same procedure was used to score each macaque cell for its cluster identity relative to its cell class (Excitatory or Inhibitory). We used the 50 most enriched and depleted markers, rather than 100, because the strength of enrichment declined more quickly down the ranked list for subtler differences between subtypes of excitatory and inhibitory neurons. Again, the annotation of cell clusters based on a t-test showed a strong match to annotation based on the most confident individual mouse markers.

To find genes specialized in a particular species, the mouse cells used were limited to the Sathyamurthy adult mouse snRNA-seq data^16,34^. Rather than the processed data, where low variance genes are removed, we used an unfiltered version of the mouse dataset. This allowed us to identify potential examples where the gene is specialized in macaque, but not mouse. Log2-fold differences were calculated using the procedure described above for both the mouse and macaque for each cell class relative to the others and for each family relative to the cell class. The family-level comparison was chosen to maximize the number of cells available for a rigorous identification in the differences in markers. We manually filtered out genes that exhibited a strong difference across species based on log2 fold change, but had very low abundance and were not significant. The marker genes of orthologous populations of cells were correlated with each other in their overall pattern (Pearson’s R ranging from 0.058 to 0.21; p-value from 1*10^-9^ to 7*10^-142^). These correlations provide further support for our assignment of cell type families. The broad range reflected the relative abundances of the cell types rather than the lack of a strong match. To determine species-specific markers, we required a gene to have an adjusted p-value of < 10^-5^ and an absolute log2 fold greater than 2. In addition, we required that the orthologous gene have a log2 fold difference of less than 0 in the other species. This highly stringent procedure is more selective for high-likelihood candidates, but may miss other candidates.

### Integration of Macaque and Mouse Dorsal Horn snRNA-Seq

Raw count matrices for macaque and adult mouse populations were read into R for analysis with the Seurat package^80^. We removed the nuclei annotated as MidVentral to focus on the dorsal horn biology. As recommended in the LIGER package, the datasets from the mouse and three macaques were each scaled, separated and integrated using the *RunOptimizeALS()* function^28^. We performed a grid search across a small number of parameter values for lambda (5,7,10) and k (15,20,30,40). The parameters lamba=7, k=30 were chosen as it produced the fewest number of cells in mouse only or macaque only cells with default parameters. The dataset was further normalized according to species and sample and visualized using the UMAP visualization with default parameters.

### Integration of Macaque, Human, and Mouse snRNA-seq

We used the dorsal horn subsets of our macaque, the Satyamurthy adult mouse, and the Yadav human datasets. Using the software package Seurat v4.2, we first normalized each dataset with SCTransform. Then we identified the genes that are variable in all datasets using *SelectIntegrationFeatures()*. Next, we identified cross-dataset anchors for cells across the pairwise combinations of the datasets using FindIntegrationAnchors, which is based on the algorithm described in . We then ran IntegrateData which uses the anchors from the previous step to construct a transformation matrix, which represents a weighted difference between the expression matrix of anchor pairs, then subtracts this transformation matrix from the original expression matrix, a process conceptually similar to batch correction and applicable to cross-species integration. The resulting combined data is in a shared integrated space allowing for direct comparisons between cells. We then performed three label transfers: mouse-to-human (to compare to Yadav et al results), as well as mouse-to-macaque and human-to-macaque (to establish the mouse and human orthologs of our macaque cell types, respectively). This was done by applying Seurat’s *FindTransferAnchors()* and *TransferData()* between the macaque, human, and mouse subsets of the integrated dataset. We then calculated silhouette scores with the package cluster v2.1 to assess confidence in label transfer of individual cells and of cell types.

Separately, we repeated the above integration steps in Seurat, but between only the mouse and macaque datasets and including the MidVentral cells. The purpose was to identify mouse orthologs of Exc-PBX3 if they belonged to the MidVentral domain. After performing integration, we observed that Exc-PBX3 most closely corresponds to Excit.25 of the mouse cells.

### Identifying Conserved Marker Gene Candidates and Visualizing Marker Genes

First, we assigned names to mouse and human dorsal horn cells based on label transfer to macaque, to establish consistency in the resolution of neuron subtypes. Next, we found cell-type-specific marker genes for each of the three species datasets using Seurat’s *FindAllMarkers()*, which identifies differentially expressed genes between groups of cells using a Wilcoxon Rank Sum test. We also ran *FindAllMarkers()* on the cross-species integrated dataset, which found differentially expressed genes after experiment-specific and species-specific signals were computationally removed via integration. Although finding differential expressed genes of the integrated dataset does not have a clear statistical interpretation, we found that the top marker genes from this procedure were effective at distinguishing cell types (see **Decision Tree Evaluation of Conserved Marker Genes**), and our goal was to select a small number of candidates from these lists. We selected top candidates from marker gene sets first based on their rank in log-fold change, and then by visualizing using violin plots and scatter plots (Seurat) how well combinations of marker genes distinguish cell types.

Marker genes for specific cell types were visualized in figures using heatmaps in Seurat, which shows the expression of individual cells belonging to a cell type with vertical ticks. We visualized scaled, imputed expression (using the ALRA method). For historical marker genes, we show un-scaled, un-imputed expression because some of the genes have low overall expression which would result in misleadingly inflated scaled and imputed signals.

### Decision Tree Evaluation of Conserved Marker Genes

After identifying conserved marker gene candidates, we quantitatively evaluated the cross-species separability of cell types using marker gene measures of expression. Because of the high rate of technical dropout in snRNA-seq for individual genes, yet the breadth of transcriptomic information available in snRNA-seq datasets which can be leveraged to infer missing expression patterns, we imputed expression signals using ALRA in order to more closely emulate experimental techniques such as RNAscope or Xenium spatial transcriptomics that have higher sensitivity measurements of specific genes. We also used scaled expression to assess relative differences in gene expression and to minimize other effects such as sequencing depth. We then extracted the cell-by-gene matrices of the macaque, human, and mouse datasets from their Seurat projects, and for a given learning problem (identifying a given cell type versus the others), we introduced binary labels for each cell whether they belonged to the target cell type (1) or another cell type (0). For each conserved cell type and using their set of two-to-three conserved marker candidates, we trained decision tree models based on 80% of the macaque cells using the rpart package. The models were then tested on the remaining 20% of the macaque cells, as well as all of the human and mouse cells. Thus, decision trees were trained based on the differences in a single species, macaque, and then evaluated for all species. We evaluated the performance of the models with the following metrics: Accuracy, Recall/Sensitivity, Specificity, Precision/Positive Predictive Value, Negative Predictive Value (**Tables S9-12**). We created visualizations of the decision tree classification boundaries for two example cell types (Exc-BNC2 and Inh-PDYN) using base R (**Figure S13**).

### RNAscope^®^ in situ hybridization

Fresh frozen lumbar spinal cord tissue samples were harvested from 3-year-old male and female macaques perfused with aCSF. The tissue was immediately placed in OCT and frozen on dry ice. L4-L6 lumbar spinal cord was sectioned using a cryostat at 20 µm thickness onto Superfrost-charged slides and stored in -80⁰C until the start of the assay. In situ hybridization was performed according to the Multiplex v2 Fluorescent (Advanced Cell Diagnostics) protocol for fresh frozen tissue after fixing the slides with cold 4% paraformaldehyde (PFA) for thirty minutes. The probes were designed and purchased from Advanced Cell Diagnostics. Signal amplification was carried out using the TSA Fluorescin, Cyanine 3 and Cyanine 5 reagents from Akoya Biosciences at 1:1500. All sections were co-stained for dapi. For in situ hybridization experiments conducted with human lumbar spinal cord, samples were fixed with cold 4% paraformaldehyde (PFA) for 15 minutes. The Multiplex v2 Fluorescent (Advanced Cell Diagnostics) protocol for fresh frozen tissue was followed with a 2-minute protease IV digestion. The Fluorescin, Cyanine 3 and Cyanine 5 reagents from Akoya Biosciences were used for probe visualization.

Combinations:

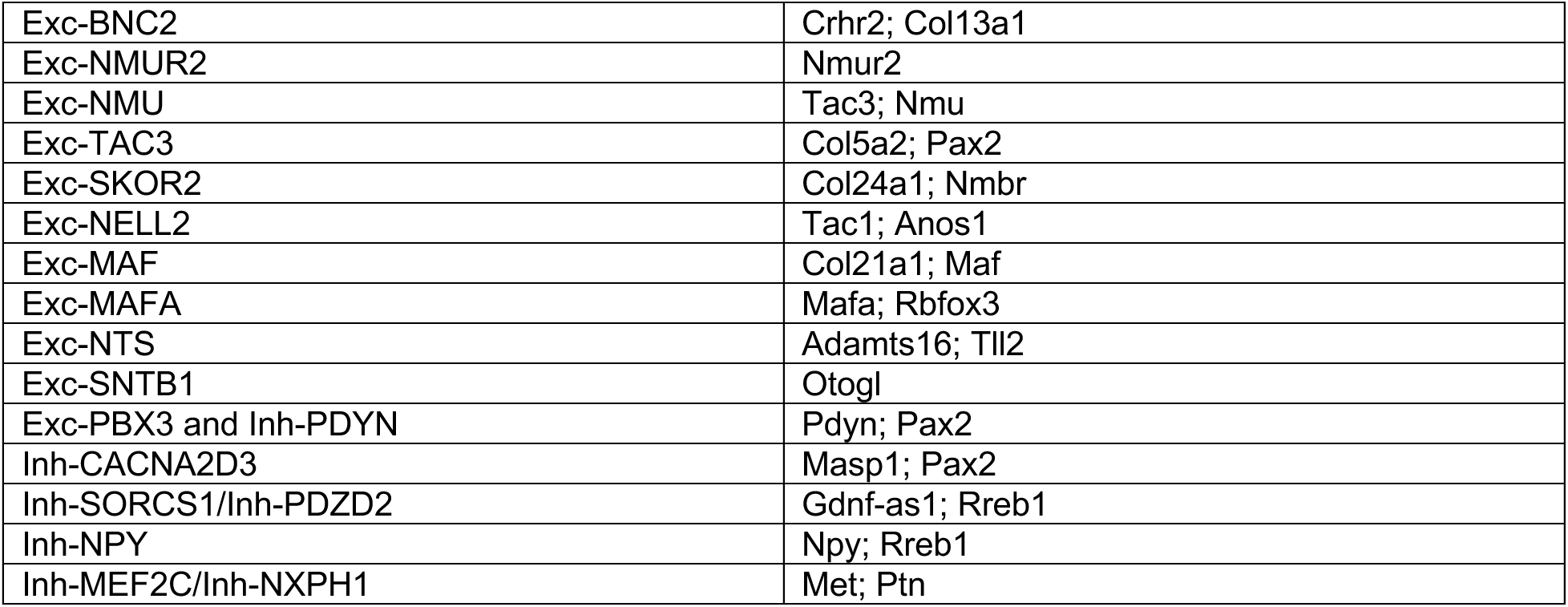

### Human and Macaque Image Acquisition and Quantification

For macaque spinal cord experiments, representative images were acquired at 10X magnification using the Nikon A1R and Nikon’s NIS-Elements imaging software and processed with ImageJ. Images taken for quantitative analysis were acquired using Nikon Eclipse 800. Laminar boundaries for the images were drawn with the Canvas X software using Atlas of the Spinal Cord: Mouse, Rat, Rhesus, Marmoset and Human as a reference. The substantia gelatinosa of the primate dorsal horn is also easily recognizable due to its translucent nature and hence was used to demarcate the boundary between laminae II and III.

For human spinal cord experiments, multiple 10x images were acquired of the dorsal horn starting from the substantia gelatinosa to lamina 10. The acquisition parameters were set based on guidelines for the FV3000 provided by Olympus. The 10x images were stitched together manually using anatomical landmarks (particularly lipofuscin) that were common between images.

For the quantitative analysis, three or more closely spaced puncta were counted as a positive cell. To account for the presence of lipofuscin in the macaque/human spinal cord tissue, the 488 channel was left blank i.e., no probe or fluorophore was added, and this was used for background subtraction.

### Experimental Procedure of Xenium Spatial Transcriptomics Assay of Mouse Spinal Cord

The first mouse (**Figure 4**) was a 10-week-old C57Bl/6 male that underwent sterile saline injection. The mouse was put under anesthesia at 4% isoflurane and then maintained at 2% isoflurane for the duration of the injection. A hamilton syringe was used to inject 5uL of saline intradermally into the left calf. The mouse was left under 1.25% isoflurane for 30 minutes after the injection. The mouse was IP injected with a solution of 100 mg/kg ketamine and 20 mg/kg xylazine one hour after the saline injection. It was then perfused using ice-cold RNAse-free PBS. The spinal cord was dissected, approximately one hour and 15 minutes after the injection of saline.

The second mouse (**Figure S18**) was a 10-week-old C57Bl/6 female, and put under at 4% isoflurane and then maintained at 1-1.5% isoflurane. The mouse’s left paw had a 0.07g von Frey filament applied for 3 seconds and was brushed, alternating between the two over the course of 5 minutes. The mouse was IP injected with a solution of 100 mg/kg ketamine and 20 mg/kg xylazine one hour after stimulus was completed. It was then perfused using ice-cold RNAse-free PBS. The spinal cord was dissected approximately one hour after stimulus.

For both mice, lumbar region was quickly isolated and embedded following 10x Genomics’ Xenium *in situ* for Fresh Frozen Tissues Preparation Guide: https://www.10xgenomics.com/support/in-situ-gene-expression/documentation/steps/tissue-prep-fresh-frozen/xenium-in-situ-spatial-profiling-for-fresh-frozen-%E2%80%93-tissue-preparation-guide. The tissue was then stored at -80 degrees Celsius before being cryosectioned at 20µm based on the Xenium guidelines. The cryosections were non-consecutive in order to capture information about distinct cells in each section. The slide was then stored again at -80 degrees Celsius until the Xenium run was completed per the protocols from 10x Genomics.

### Analysis of Xenium Spatial Transcriptomics of Mouse Spinal Cord

We used Seurat v5 to load the full field of view of the Xenium run, which consisted of eight non-consecutive spinal cord sections (description of experiment above). First, we removed cells with zero gene counts. Next, we performed un-biased clustering of the full set of Xenium cells using the Seurat functions *SCTransform()*, *RunPCA()*, *RunUMAP()*, *FindNeighbors()*, and *FindClusters()*, which normalizes expression signals and performs dimensionality reduction using PCA, calculates k-nearest neighbors for cells and constructs a shared nearest-neighbor graph, then uses this graph to optimize community detection (*i.e.* clustering) using the smart local moving algorithm. Next, we normalized the Satymathurthy adult mouse dataset (all cells including glia and other) with SCTransform, subsetted the available features to those within the Xenium assay, and integrated the Xenium and Satyamurthy datasets. We assigned to the Xenium clusters the label of the nearest cluster from Satyamurthy et al within the shared integration space based on UMAP visualization. A Xenium cluster was assigned the label ‘Neurons’ if the nearest Satyamurthy cluster was any of the neuron subtypes. We then visualized the locations of the Xenium cell types within the spinal cord sections, verifying the known spatial patterns of major cell classes such as high neuron density in grey matter, high oligodendrocyte density in white matter, etc.

Next, we subsetted to Xenium cells now labeled Neurons, and re-clustered the neurons only with the same steps described above. Then we integrated with the Satyamurthy dorsal horn subset of neurons. We found that a large region of Xenium cells did not align closely with any of the dorsal horn neurons, besides Exc-PBX3, which we previously showed as being likely MidVentral. We concluded that these Xenium clusters were all MidVentral/Ventral, subsetted them out, then repeated the same clustering and integration steps with just the Xenium putative dorsal horn neurons and Satyamurthy dorsal horn neurons. We found that now all Xenium clusters aligned closely to a specific labeled neuron subtype, with the exceptions that (a) one Xenium cluster was not clearly Exc-NMUR2 or Exc-SKOR2 and was left unlabeled, and (b) the single Xenium cluster 2 corresponded to the combined Inh-SORCS1 and Inh-PDZD2 cell types even after trying several resolutions (0.5 to 1.3) of the clustering algorithm. Inh-SORCS1 and Inh-PDZD2 were previously established as closely related cell types and cluster 2 was labeled Inh-SORCS1/Inh-PDZD2.

Once neuron subtype labels were assigned to Xenium dorsal horn neurons, we visualized the positions of these cells in the Xenium image and hypothesized that relative shallowness of these cell types would correspond to those previously found with macaque RNAscope. To quantify this, we estimated the vertical position of the image of each dorsal horn neuron relative to the average vertical position of all other dorsal horn neurons within the same dorsal horn section (two dorsal sections per spinal cord section = 16). By convention dorsal horns were aligned at the top of the image so that higher vertical position corresponded to a shallower laminar position. We calculated the following z-score metric z = x - u / s, where x is the vertical position of an individual cell, u is the average vertical position of dorsal horn neurons in the same slice, and s is the standard deviation of the positions of the dorsal horn neurons in each slice. After calculating the per-cell z-score for each cell, we tested if the distribution of each cell type’s position was significantly different from average in either the shallow or deep direction, using the Wilcoxon signed-ranked test and multiple hypothesis correction using Holm’s method. This was done with the distributions of individual cells aggregated across slices, and the per-slice distribution of the average z-scores of each cell type in each slice, with similar results. Visualization of results are shown in the main figure for the per-slice distributions, and supplemental figure for the per-cell distributions.

### Mouse Gene-Based Peaks from Introns and Flanking Regions

For each cell type, we identified non-coding regions nearby the mouse-specific marker genes previously found with Seurat. Specifically, we aggregated the intronic regions and flanking regions 20KB on either side of each marker gene, which defines the gene-based peak foreground for that cell type. The background for all cell types was defined as the intronic and flanking regions (same distance) for all coding genes. The foregrounds and backgrounds as defined were first mapped to human orthologous regions (see **Cross-Species Peak Mapping)**, and then used as the inputs to LDSC, described in **Cell-type-specific GWAS SNP enrichment**.

### Cross-Species Peak Mapping

Peaks from above-described mouse datasets were originally in mm10 genome coordinates, while SNPs from GWAS are in human (hg38) coordinates. Non-coding regions such as enhancers generally exhibit less sequence conservation than protein-coding genes, and identifying orthologous regions requires a more flexible mapping technique. Thus, we mapped mouse mm10 peaks to human hg38 orthologs using the mapping tool HalLiftover in conjunction with HALPER, which constructs contiguous ortholog regions from HalLiftover outputs. Further details are described in the HALPER publication and ^67^.

### Mouse single nucleus ATAC sequencing

Twenty sex-matched B6 mice were deeply anesthetized with Isoflurane, decapitated and the spinal cord was dissected out. Then, the lumbar spinal cord from L3-5 was mounted on an agarose mold and the dorsal halve was micro dissected on a vibratome (Leica Microsystems). Next, the tissue was Dounce homogenized with a loose pestle in 2 ml of Lysis Buffer containing in mM: 10 Tris-Cl, 10 NaCl, 3 MgCl2; ∼25 strokes, and filtered through a 70mm cell strainer, followed by a centrifugation step at 400g for 5min at 4°C. After that, the pellet was resuspended in 2 ml of Nuclei Wash and Resuspension Buffer (1X PBS, 1% BSA and RNase inhibitor 0.2U/mL) and filtered-through a 40mm cell strainer to optimize myelin and debris removal. Finally, nuclei viability was assessed by trypan blue staining and sequenced at 10X Genomics Core at the University of Pittsburgh. The 10x libraries were processed according to the manufacturer’s instructions. Completed libraries were run on the Novaseq 6000 (Illumina).

### Mouse snATAC-seq Pre-Processing, Cell Labeling, and Peak Calling

Bcl files were converted to fastq format using the *cellranger mkfastq* command line tool. Fastq reads were then aligned to the mm10 genome using Chromap. Subsequent steps were done with the software suite ArchRv1.0. Arrow files were generated from the aligned fragments and doublet scores were generated using ArchR’s built-in doublet algorithm. An ArchR project was then generated from the Arrow files and sample metadata from the sequencing run. Cells were filtered with minimum transcription start site (TSS) enrichment = 8, minimum number of unique fragments 10^3.5, and doublets filtered out with cutoff enrichment threshold of 0.5. Fragment periodicity plots were also inspected for intact periodicity from nucleosomes.

Un-biased clustering of cells was done using Iterative LSI, with variable features looped between 30 and 230. 200 variable features was selected having a favorable resolution of clustering with minimal batch effects. Batch correction was performed with Harmony. Enrichment of marker genes for major cell types was visualized based on gene score, which is a proxy for gene expression based on accessibility at and around the TSS of the gene. Established marker genes used to identify major cell types are provided in Supplementary Table S13.

Once neuron clusters were identified by marker genes, two separate neuron and glia ArchR projects were generated and we re-clustered cells again with iterative LSI. The purpose of separating neurons from glia is for the clustering algorithm to learn variability in accessibility between neural subtypes, which would otherwise be dominated by the differences among neurons and glia cell types.

Clusters within the neuron sub-project were labeled based on ATAC-RNA integration. Originally we attempted using the Satyamurthy mouse snRNA-seq dataset for integration, but found issues with the generated co-embeddings, and switched to using macaque. This is possibly because the macaque dataset has higher sequencing depth. We used the macaque snRNA-seq dataset with conserved dorsal horn cell type labels (along with unlabeled Mid and Ventral neurons) and we named the macaque genes the orthologous mouse gene names, in order to match with annotated mouse gene scores. We then performed unconstrained integration using ArchR’s implementation of Seurat’s label transfer algorithm using the command addGeneIntegrationMatrix. RNA-ATAC co-embeddings were generated and shown in **Figure S26**. After integration and putative cell labeling, we also visualized in UMAPs the marker gene expression in clusters now labeled as dorsal horn neuron subtypes. We used the macaque and conserved marker genes previously discussed.

We re-combined the neuron and glia projects and generated reproducible peaks for dorsal horn neuron subtypes and glia major cell types using ArchR’s iterative overlap method, and determined peak signals (peak calling) with the statistical method MACS2^85^. For GWAS enrichment analysis, we exported called peaks to narrowpeak files with summit-centered peaks of fixed 501bp length, which defines our neuron subtype-specific foregrounds. The background for each subtype was the union of all foregrounds with the ENCODE Mouse UCSC DNAse Hypersensitivity Sites: https://storage.googleapis.com/encode-pipeline-genome-data/mm10/ataqc/mm10_univ_dhs_ucsc.bed.gz

The foregrounds and backgrounds as defined were first mapped to human hg38 (see **Cross-Species Peak Mapping)**, and then used as the inputs to LDSC, described in **Cell-type-specific GWAS SNP enrichment**.

### Human Immune Cell snATAC-seq, Human Brain Tissue-specific ATAC-seq, and Mouse Bulk Liver ATAC-seq Pre-Processing

#### Human Immune

We downloaded publicly available ATAC-Seq datasets of Naive, IFNγ-treated, and IFNβ-treated CD14+ human monocyte-derived macrophages from Cheng et al. (2019) available at NIH SRA, https://www.ncbi.nlm.nih.gov/sra (Accession number: SRP145626). We reprocessed these datasets, identifying reproducible peaks with the irreproducible discovery rate (IDR) method^86^ using the ENCODE ATAC-seq pipeline (https://github.com/ENCODE-DCC/atac-seq-pipeline) with default parameters. This pipeline filters low-quality reads, aligns reads to the hg38 reference genome via bowtie2^87^, calls peaks with MACS2^85^, excludes peaks in regions with unreliable read mapping^88^, and ^88^sesses peak reproducibility using IDR. The IDR reproducible peak sets were called across pooled pseudo-replicates, and we obtained 31,497 peaks for Naive macrophages, 34522 peaks for IFNβ-treated macrophages, and 74,014 peaks for IFNβ-treated macrophages. The set of foregrounds was the set of peaks for each of the three cell types. The background for all cell types was the union of the merged foregrounds along with all merged annotations from the Roadmap Epigenomics Projects.

#### Human Brain Tissue

We downloaded publicly available NeuN-sorted ATAC-seq^52^ of human postmortem brain from the Read Archive through Gene Expression Omnibus (GEO) accession #GSE96949. We re-processed the dataset using the ENCODE pipeline as described above for *Human Immune.* The two foregrounds were the putamen and hippocampus bulk peak sets, and background for both tissues was the union of the merged foregrounds along with all merged annotations from the Roadmap Epigenomics Projects.

#### Mouse human liver

The experimental methods, data generation and pre-processing of this dataset are described in ^51^. Peaks from this study were then mapped from mm10 to hg38 with HALPER (see **Cross-Species Peak Mapping** below). There was a single foreground, the set of peaks from the bulk ATAC-seq. The background was the union of the foreground with the ENCODE Mouse UCSC DNAse Hypersensitivity Sites. For each of these datasets, the foregrounds and backgrounds as defined for each, already in hg38 coordinates, were used as the inputs to LDSC, described in **Cell-type-specific GWAS SNP enrichment**.

### Cell-type-specific GWAS SNP enrichment (Linkage-Disequilibrium Score Regression)

We received the summary statistics from the corresponding authors of Kupari et al. and Khoury et al. We munged the raw summary statistics as described for linkage-disequilibrium score regression analysis https://github.com/bulik/ldsc. For all peak datasets described in above sections (human orthologs if applicable: mouse gene-based, mouse bulk liver ATAC-seq, mouse immune, human brain), we estimated enrichment of chronic pain GWAS SNPs using stratified linkage-disequilibrium score regression (S-LDSC). S-LDSC estimates the contribution of an annotation (in this case, tissue or cell-type-specific non-coding peaks) to per-SNP heritability in a joint model versus background. Further details are described by which used the same procedure. Briefly, we annotated the LD scores for each foreground and background set using the hg38 1000 Genomes European Phase 3 European super-population (1000G EUR) cohort. Then we estimated conditioned heritability enrichment. This was done separately for each combination of tissue or cell type along with each chronic pain trait. To adjust for multiple hypotheses, we used false discovery rate (FDR) correction (0.05 threshold) because of the inherent correlations between hypothesis tests. Resulting adjusted p-values and false-discovery-corrected significance were visualized.

### Identification of Enriched Peaks Nearby Spinal Injury-Associated Genes

Browser visualizations were first done with the Interactive Genome Viewer (IGV)^89^ and LocusZoom^49^, and figures were generated with the package Plotgardener^90^. First, marginally significant chronic-pain-associated SNPs from ^18^ and ^26^ were examined in single-base-pair coordinates for each study. SNPS at top loci were further scrutinized if they were estimated to be plausibly causal based on fine-mapping and generation of a credible set of SNPs by LocusZoom. We also verified that candidate SNPs were not in strong LD with exonic SNPs. These credible SNPs were intersected with mouse dorsal horn subtype-specific snATAC-seq peaks (see **Mouse snATAC-seq Pre-Processing, Cell Labeling, and Peak Calling**) using the SubsetByOverlaps function for GRanges objects. We examined if SNPs were eQTLs of nearby genes using the GTEx portal (https://www.gtexportal.org/home/snp/rs2287425), and searched the literature to see if nearby genes were linked to dorsal horn development or chronic pain. LANCL1 and FOXP2 were the two candidate loci highlighted here that were identified by this procedure.

### Transcription Factor Regulator Candidates in mouse snATAC-seq

We identified transcription factor (TF) regulator candidates in accordance with the ArchR manual. First, we computed motif enrichment and cell-type-specific variability in enrichment (deviations) using ArchR’s implementation of chromVAR and computed the motif z-scores for all motifs. This generated a cell-by-motif deviations matrix. We computed the correlations of this TF matrix with gene expression of transcription factor genes, using the integrated snRNA-seq gene data for the gene expression matrix. Per the standard ArchR guidelines, TF regulators were identified as having a correlation between motif deviation and TF expression of 0.5 or greater, as well as a high average deviation of at least the 75th percentile. We note that integrated RNA expression is only a proxy for true single-nucleus accessibility and gene expression relationships, because integrated RNA signals are not actually from the same cell or experiment. Thus, we also considered TF motifs that had a non-negative correlation with TF expression and were in the 95th percentile or greater for highest motif deviation. The purpose was to capture strong TF regulator candidates based on particularly strong deviation signals in cases where RNA expression accuracy may be limited.

