## Supplemental Figures for "Spatial, transcriptomic and epigenomic analyses link dorsal horn neurons to chronic pain genetic predisposition"

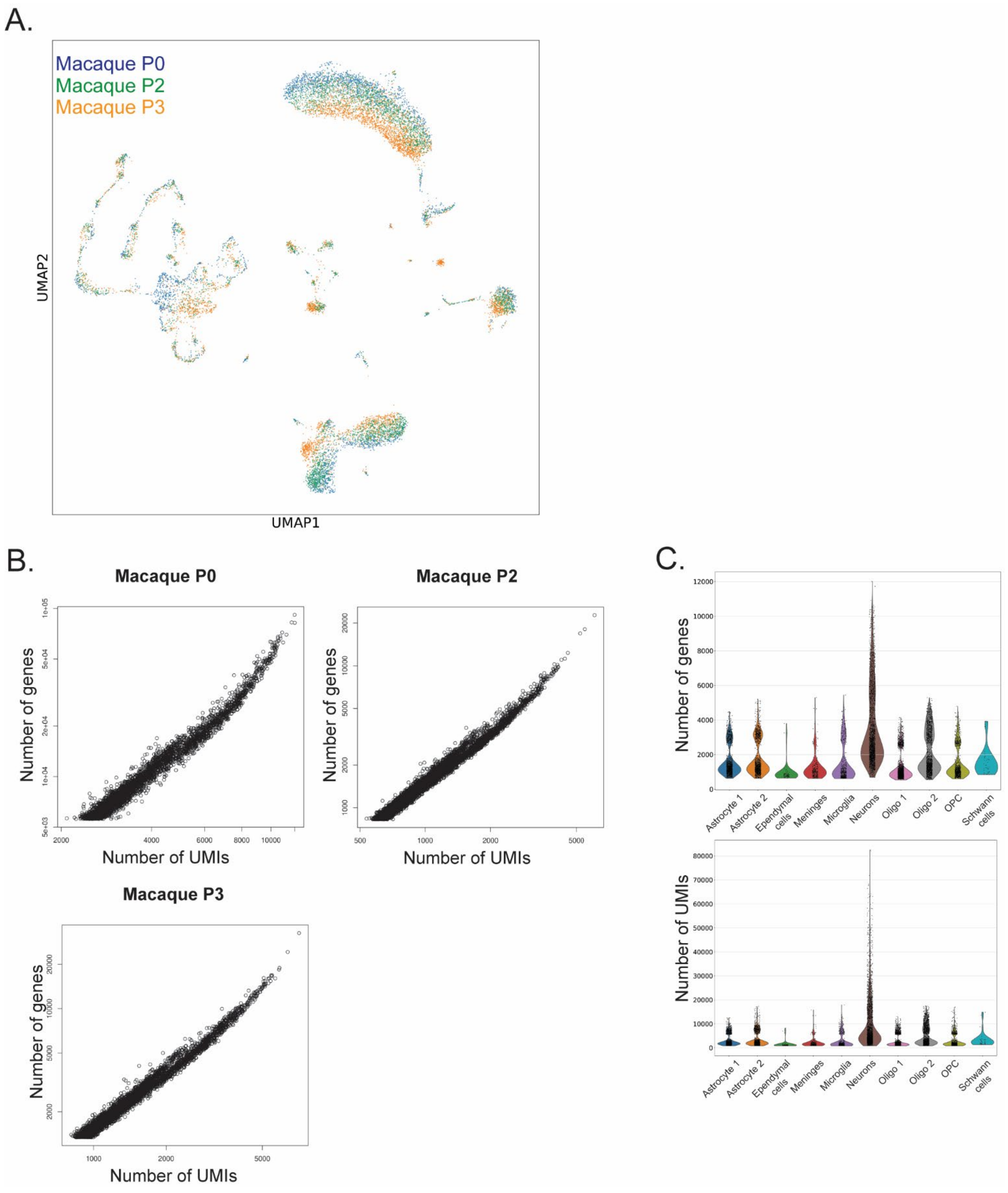


**Supplemental Fig. S1. snRNA-seq quality control metrics for nuclei obtained from three macaque biological replicates, related to Main Figure 1. A,** UMAP visualization of the dorsal horn nuclei from the three biological replicates after co-normalization. **B,** The number of detected unique molecular identifiers (UMIs), which represent unique reads (x-axis) is highly correlated to the number of detected genes (y-axis) for each of the replicates. **C,** Violin plot showing the number of genes (Top) and number of UMIs (Bottom) across the major cell types.

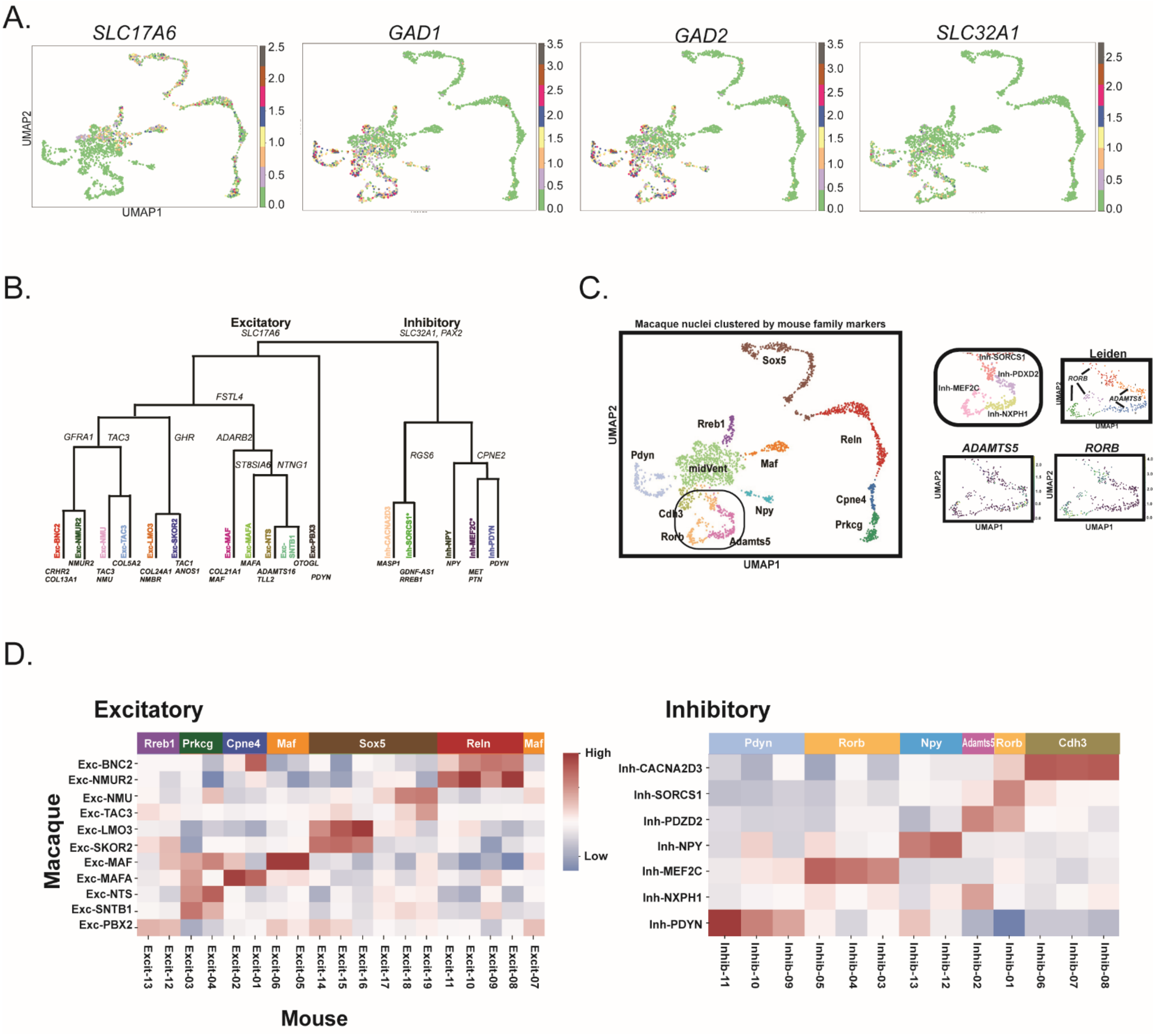

**Supplemental Fig. S2. Macaque snRNA-seq neuronal clusters including the midventral domain, related to Main Figure 1.** UMAP representations showing gene expression of vesicular glutamate transporter 2 (SLC17A6), vesicular inhibitory amino acid transporter (SLC32A1), and GABA synthesizing genes i.e. glutamate decarboxylase 1 and 2 (GAD1 and GAD2) in the macaque clusters. B. Dendrogram of cell clusters divided by excitatory (SLC17A6) and inhibitory (SLC32A/PAX2) showing marker genes for each split point. Unique markers for each of the macaque neuron subtypes are also shown. C. Leiden map of inhibitory clusters Inh-SORCS1, Inh-PDZD2, Inh MEF2C, and Inh-NXPH1. Interestingly Inh-SORCS1 and Inh-MEF2C match to the Rorb mouse family and Inh-PDZD2 and Inh-NXPH1 map to the ADAMTS5 mouse family. D. Heatmap of macaque nuclei clustered based on their correlation (red-high; blue- low) to the mouse excitatory (top) and inhibitory clusters (bottom) and families from the Russ et al., 2021 dataset.

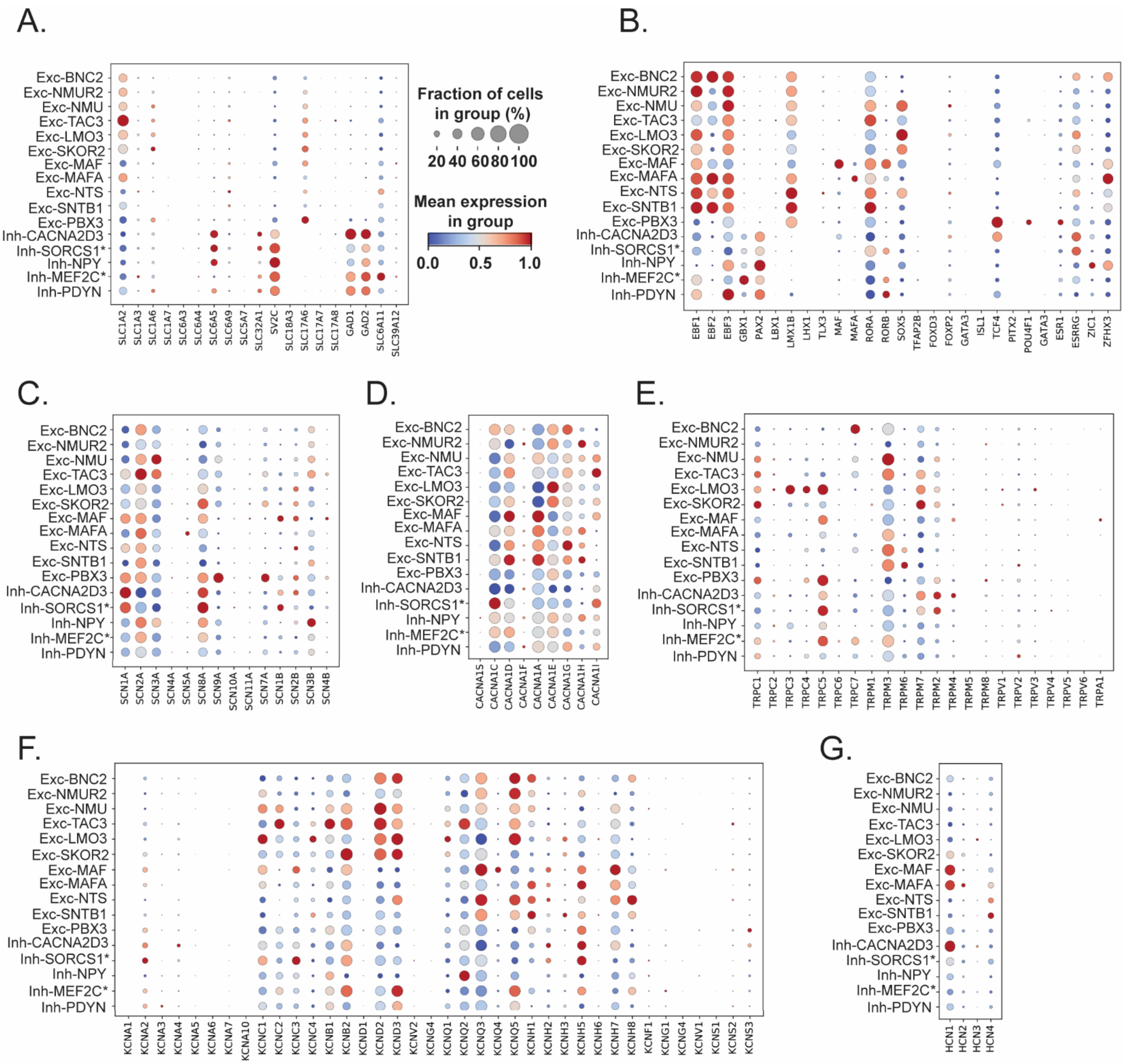


**Supplemental Fig. S3. Macaque snRNA-seq expression of neurotransmitter-related enzymes, transporters, transcription factors and channels across the cell types, related to Main Figure 1.** A, dotpot showing mean expression of neurotransmitter-related enzymes and transporters colored from blue=0 to red=1. B, dotplot showing mean expression of transcription factors. C, dotplot showing mean expression of sodium channels. D, dotplot showing mean expression of calcium channels. E, dotplot showing mean expression of transient receptor potential (TRP) channels. F, dotplot showing mean expression of potassium channels. G, dotplot showing mean expression of hyperpolarization-activated cyclic nucleotide (HCN) channels.

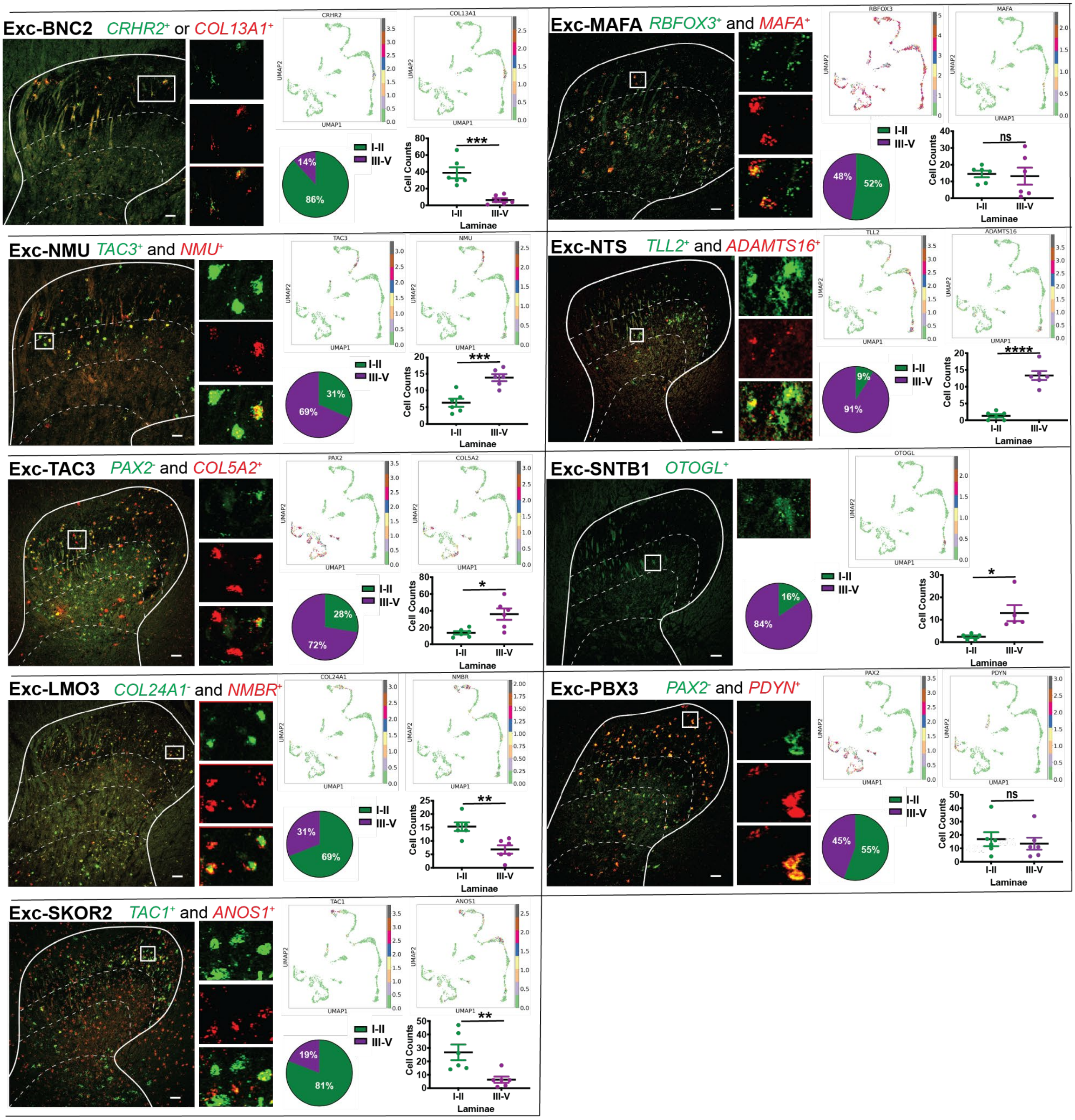


**Supplemental Fig. S4. RNAscope identification of laminar distribution in macaque, additional dorsal horn glutamatergic cell types, related to Main Figure 2.** RNAscope results for the excitatory neuron subtypes. Panels depict the *in situ* hybridization marker gene combinations used to detect each target cell type, with the color of the marker name indicating the fluorescence of the marker in the image. Dorsal horn images are taken at 10x with the smaller insets showing a magnified image (20x) of the individual gene(s) as well as the merged image. Laminar boundaries (dashed lines) are drawn between II/III, III/IV and IV/V. Scale bars in bottom right corners of panels = 100 µm. UMAPs of the marker genes used in the in situ are shown. Pie charts show the percentage of cells expressing the cluster marker genes in superficial (I-II) and deep (III-V) dorsal horn. Histograms show the number of cells positive for the cluster marker gene(s) binned into superficial or deep dorsal horn for each spinal cord section. n=5 or 6 spinal cord sections from N=2-3 macaques. *p<0.05, **p<0.01, ***p<0.001, ****p<0.0001, not significant (ns).

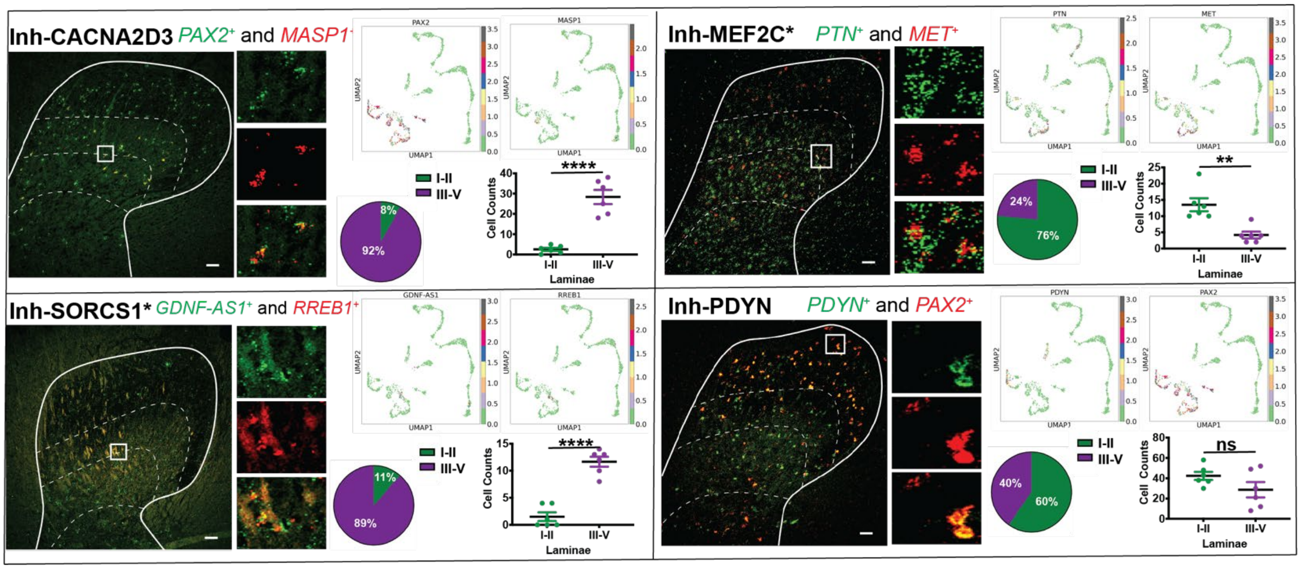


**Supplemental Fig. S5. RNAscope identification of laminar distribution in macaque, additional dorsal horn GABAergic cell types, related to Main Figure 2.** RNAscope results for the inhibitory neuron subtypes. Panels depict the *in situ* hybridization marker gene combinations used to detect each target cell type, with the color of the marker name indicating the fluorescence of the marker in the image. Dorsal horn images are taken at 10x with the smaller insets showing a magnified image (20x) of the individual gene(s) as well as the merged image. Laminar boundaries (dashed lines) are drawn between II/III, III/IV and IV/V. Scale bars in bottom right corners of panels = 100 µm. UMAPs of the marker genes used in the *in situ* are shown. Pie charts show the percentage of cells expressing the cluster marker genes in superficial (I-II) and deep (III-V) dorsal horn. Histograms show the number of cells positive for the cluster marker gene(s) binned into superficial or deep dorsal horn for each spinal cord section. n=5 or 6 spinal cord sections from N=2-3 macaques. *p<0.05, **p<0.01, ***p<0.001, ****p<0.0001, not significant (ns).

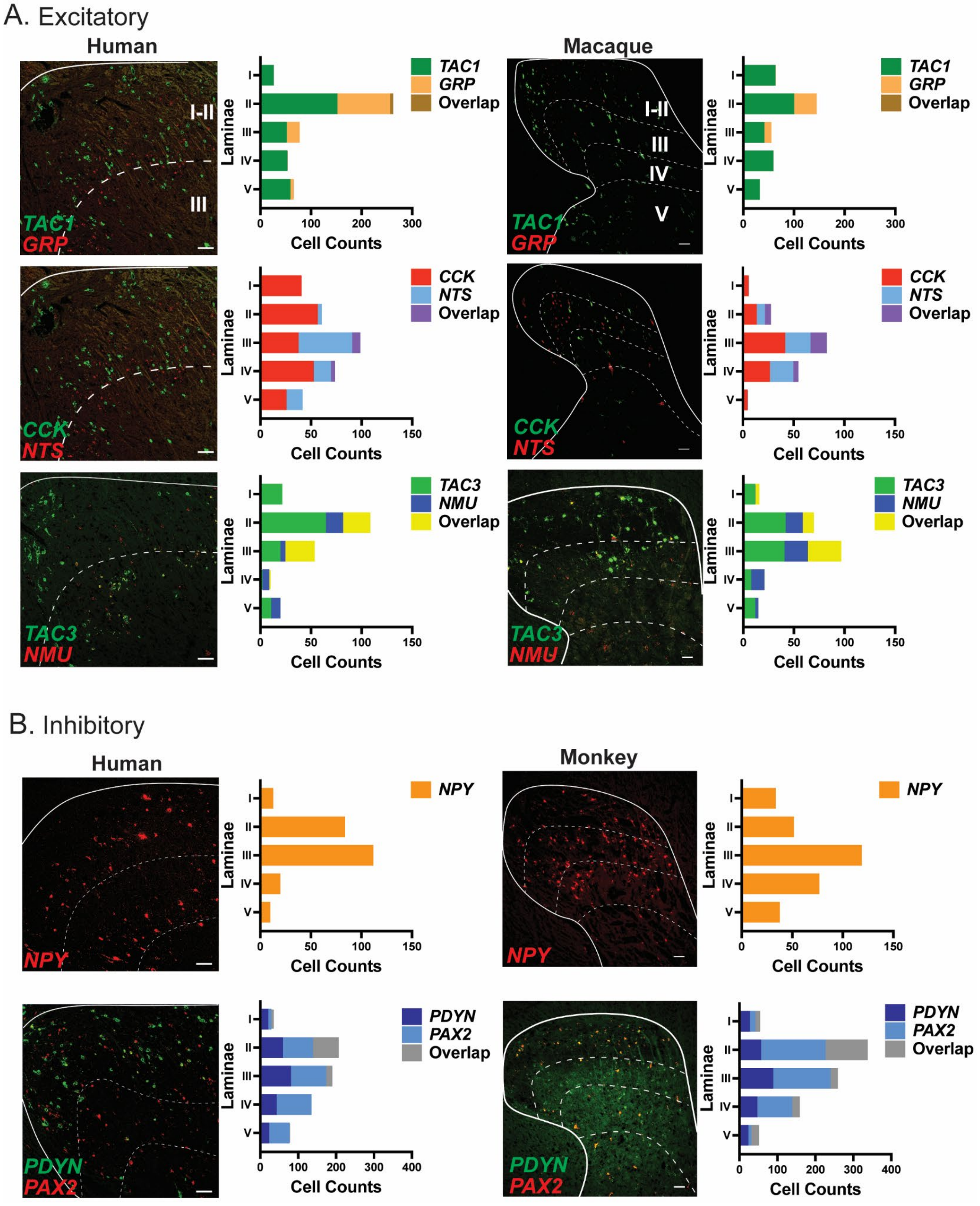


**Supplemental Fig. S6. Cross-species Comparison of Neuropeptides Across Macaque and Human, related to Main Figure 2. A**, Representative human dorsal horn in situ images of excitatory neuropeptides – TAC1, GRP, CCK, NTS, TAC3, and NMU after background subtraction to remove lipofuscin (left). **B**, Representative human dorsal horn in situ images of inhibitory neuropeptides– NPY, PDYN and PAX2 after background subtraction to remove lipofuscin (left). Scale bar = 100 μm. Histograms show total in situ hybridization counts of neuropeptides from macaque and human lumbar dorsal horn. n=6 hemisections from N=2-3 animals/donors.

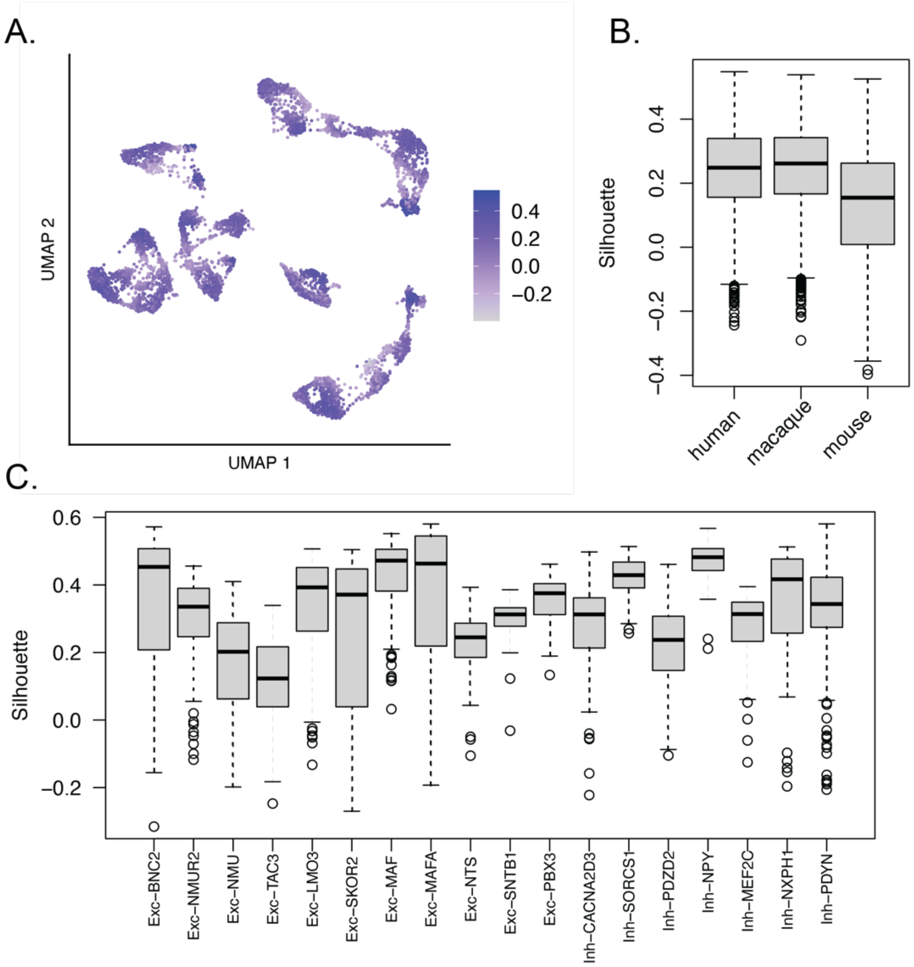


**Supplemental Fig. S7. Silhouette Scores of Cross-Species Integration, related to Main Figure 3.**
**A**, Per-cell silhouette scores in a UMAP representation of the macaque-human-mouse integrated datasets. **B,** Silhouette Score distributions stratified by species. **C,** For macaque, silhouette score distributions stratified by dorsal horn cell type.

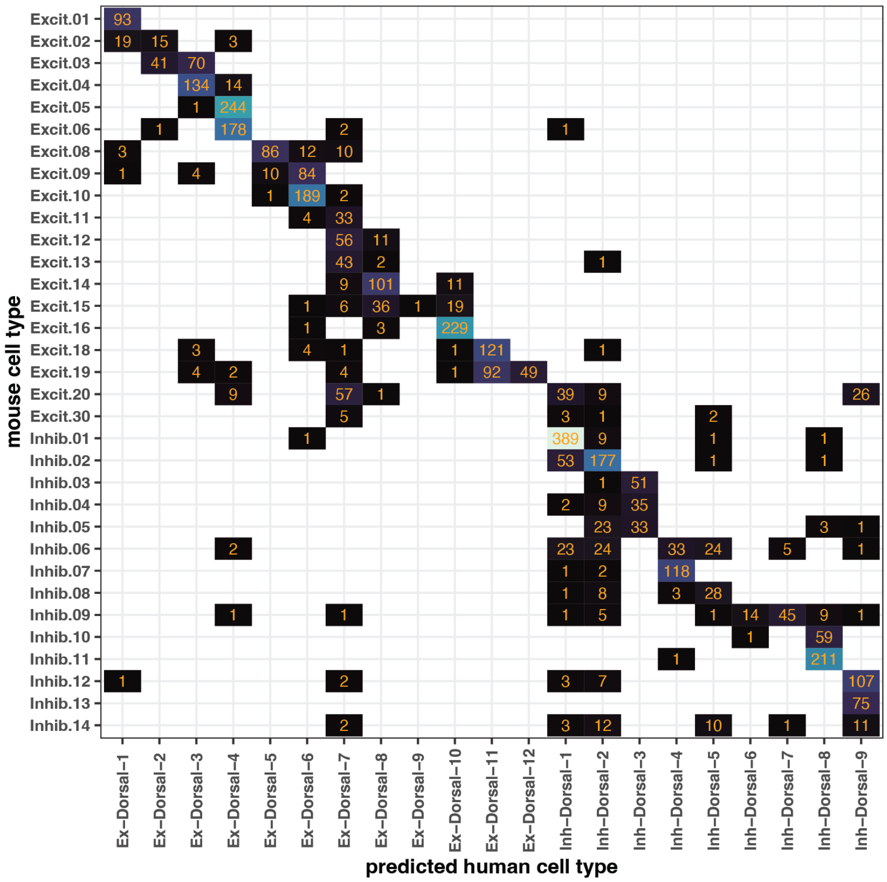


**Supplemental Fig. S8. Transfer of human labels to mouse following cross-species integration, related to Main Figure 3.** Following integration of macaque, human, and mouse: each number in block (X,Y) represents the number of cells of mouse cell type Y that are predicted to be human cell type X based on label transfer. This mouse-human labeling reproduces the correspondence between cell types in Figure 5C of Yadav *et. al.*

A.

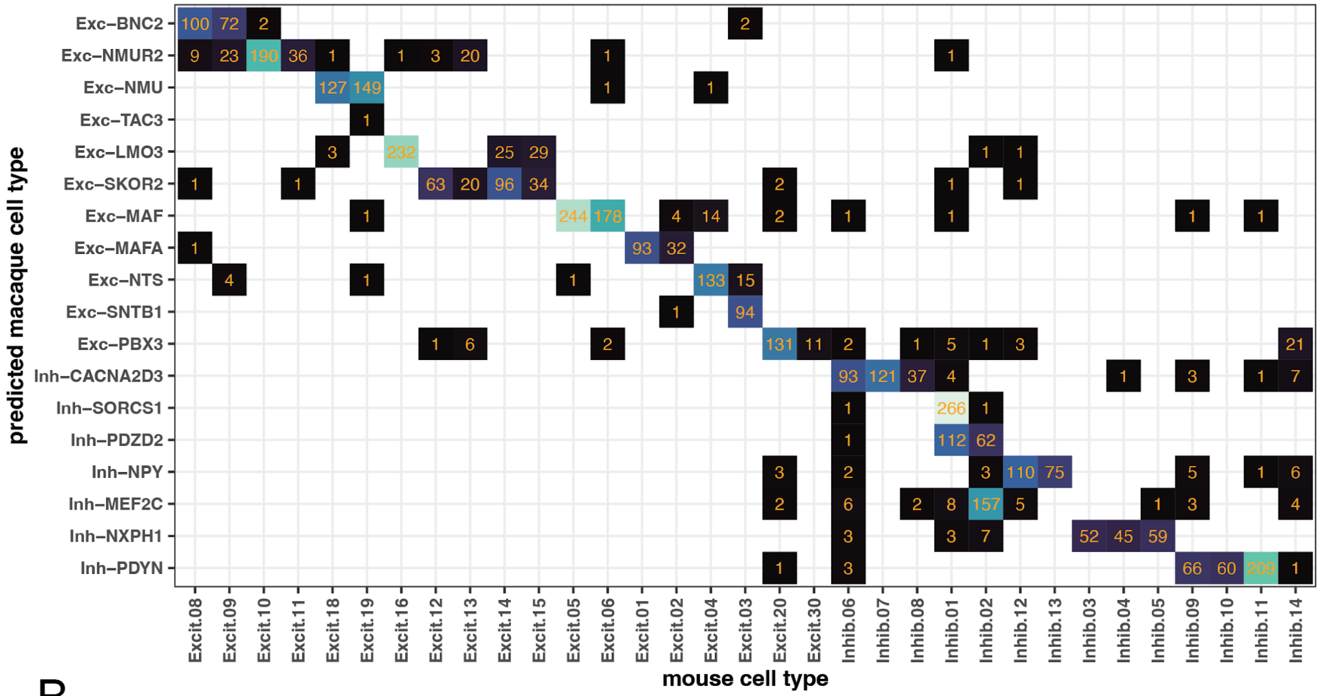


B.

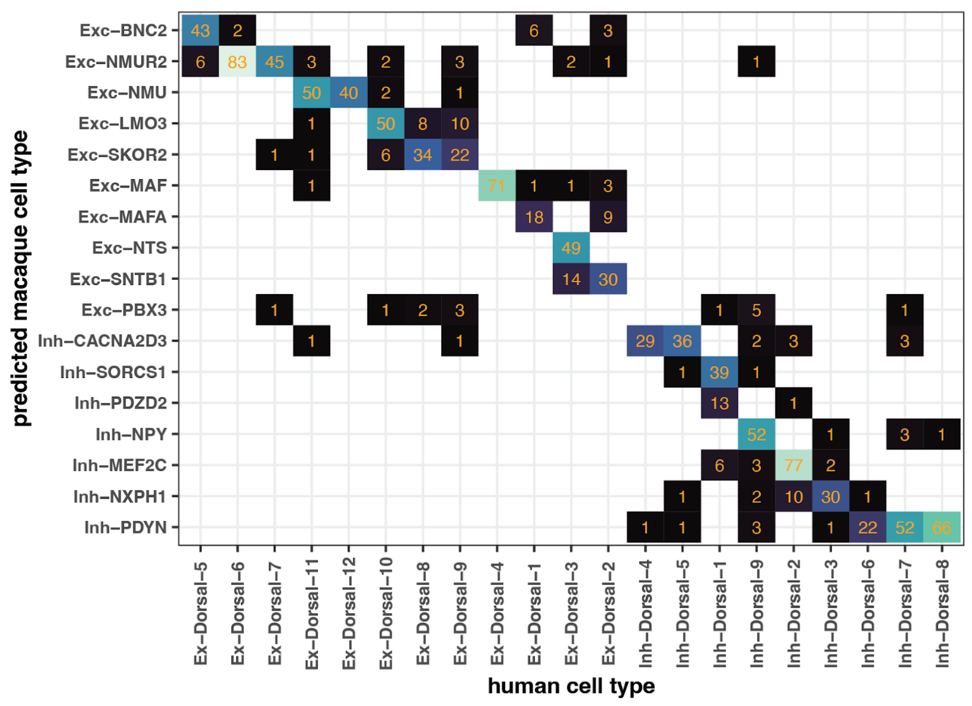


**Supplemental Fig. S9. Transfer of macaque labels to mouse and human cell types following cross-species integration, related to Main Figure 3.** Following integration of macaque, human, and mouse. **A,** each number in block (X,Y) represents the number of cells of mouse cell type X that are predicted to be macaque cell type Y based on label transfer. **B,** each number in block (X,Y) represents the number of cells of human cell type X that are predicted to be macaque cell type Y.

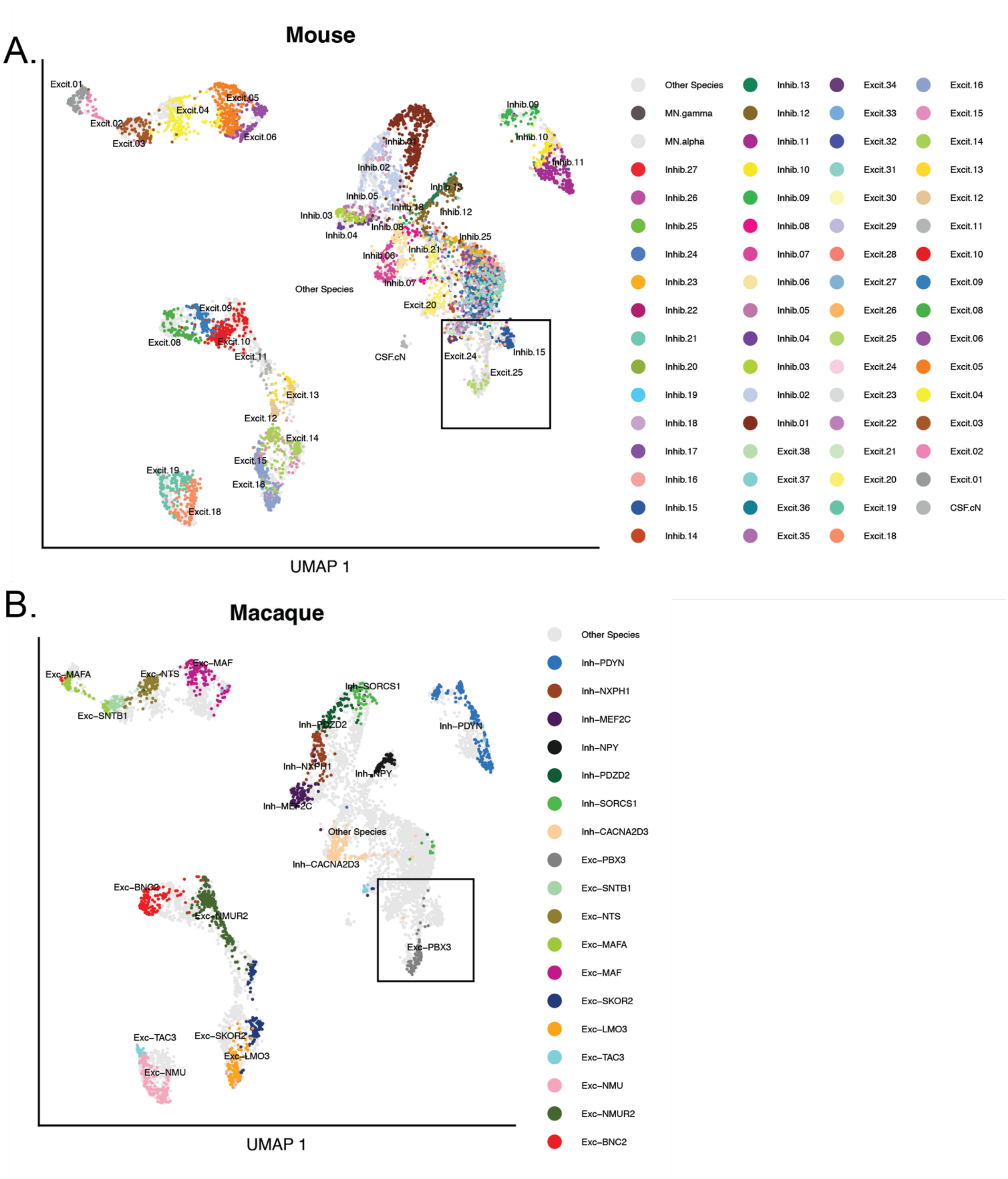


**Supplemental Fig. S10. Mouse-Macaque integration of all neurons to identify mouse ortholog of Exc-PBX-3, related to Main Figure 3.** UMAP representation of mouse and macaque neurons overlayed. Colors and labels represent the cell types for the indicated species. In-line rectangles highlight the location of Exc-PBX3 and nearest mouse ortholog Excit.25. **A**, Mouse cells in color, macaque in light grey. **B,** Macaque cells in color, mouse in light grey.

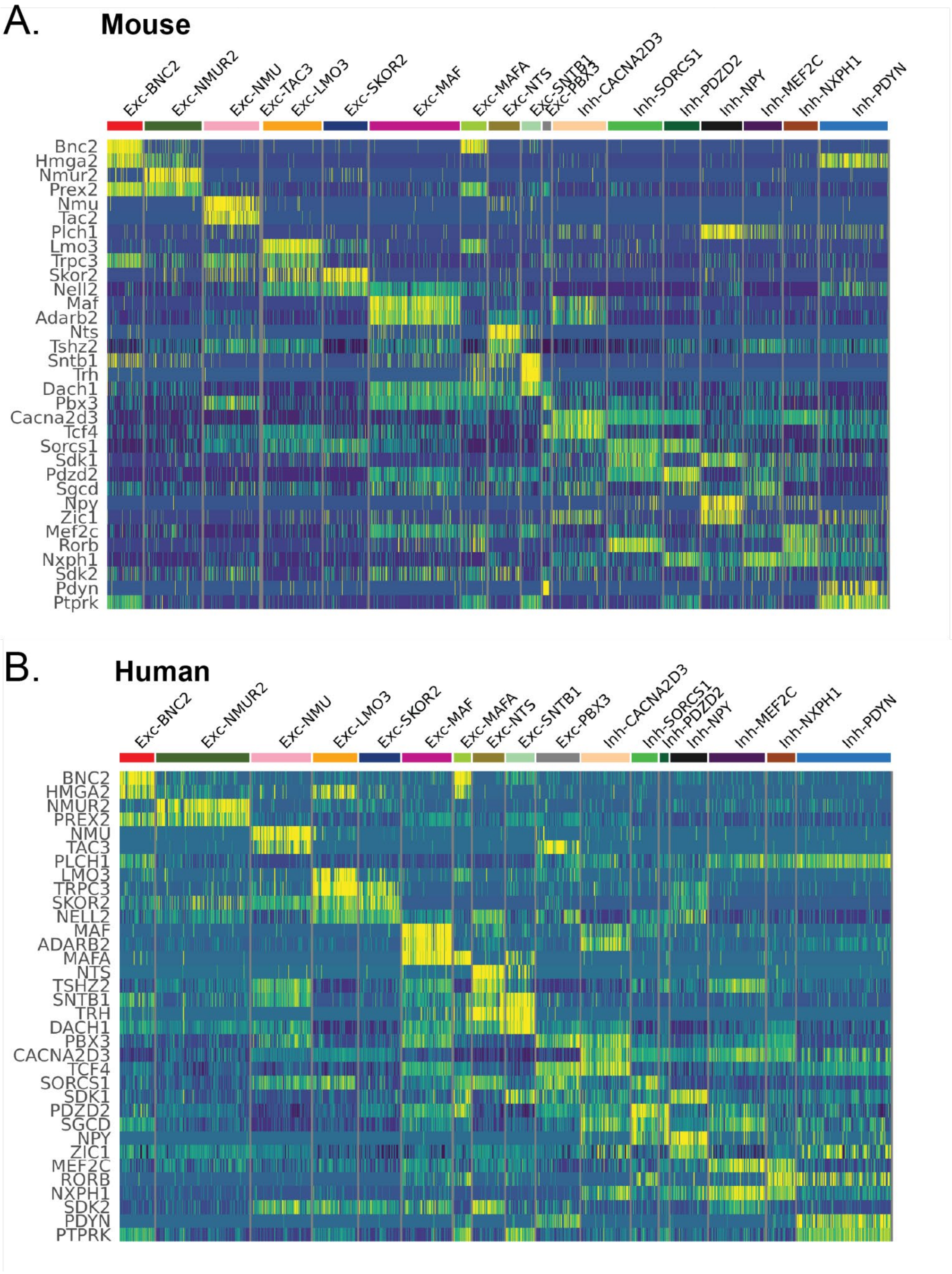


**Supplemental Fig. S11. Gene expression of conserved marker genes in human and mouse, related to Main Figure 3.** Scaled, imputed gene expression for each cell type (x-axis) and marker gene (y-axis). Each vertical tick represents the gene expression level of a single cell. **A,** Human. **B,** Mouse.

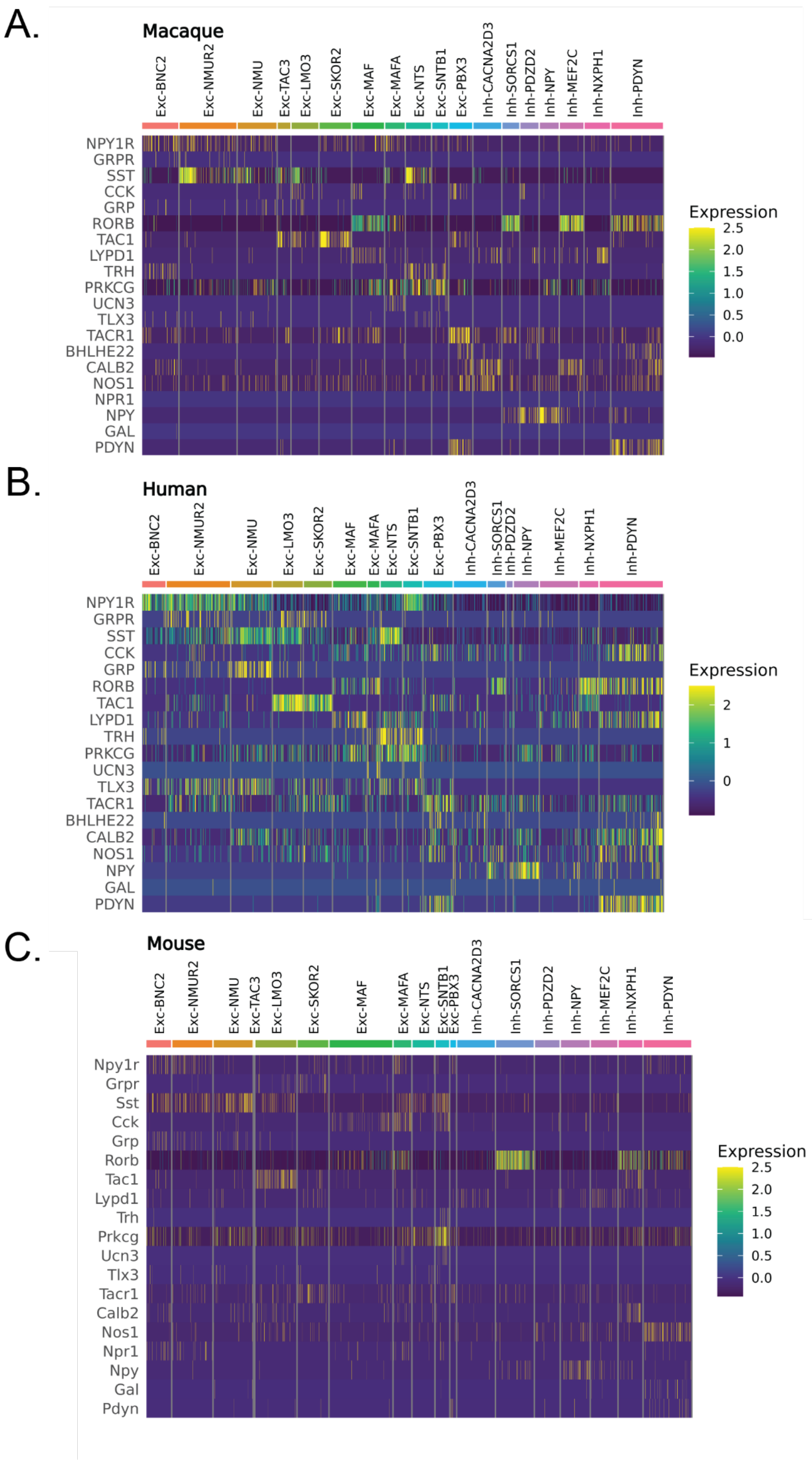


**Supplemental Fig. S12. Gene expression of historically important marker genes in macaque, human and mouse, related to Main Figure 3.** Un-scaled, SCTransformed, non-imputed expression for each cell type (x-axis) and marker gene (y-axis). Each vertical tick represents the gene expression level of a single cell. **A,** Macaque. **B,** Human. **C,** Mouse.

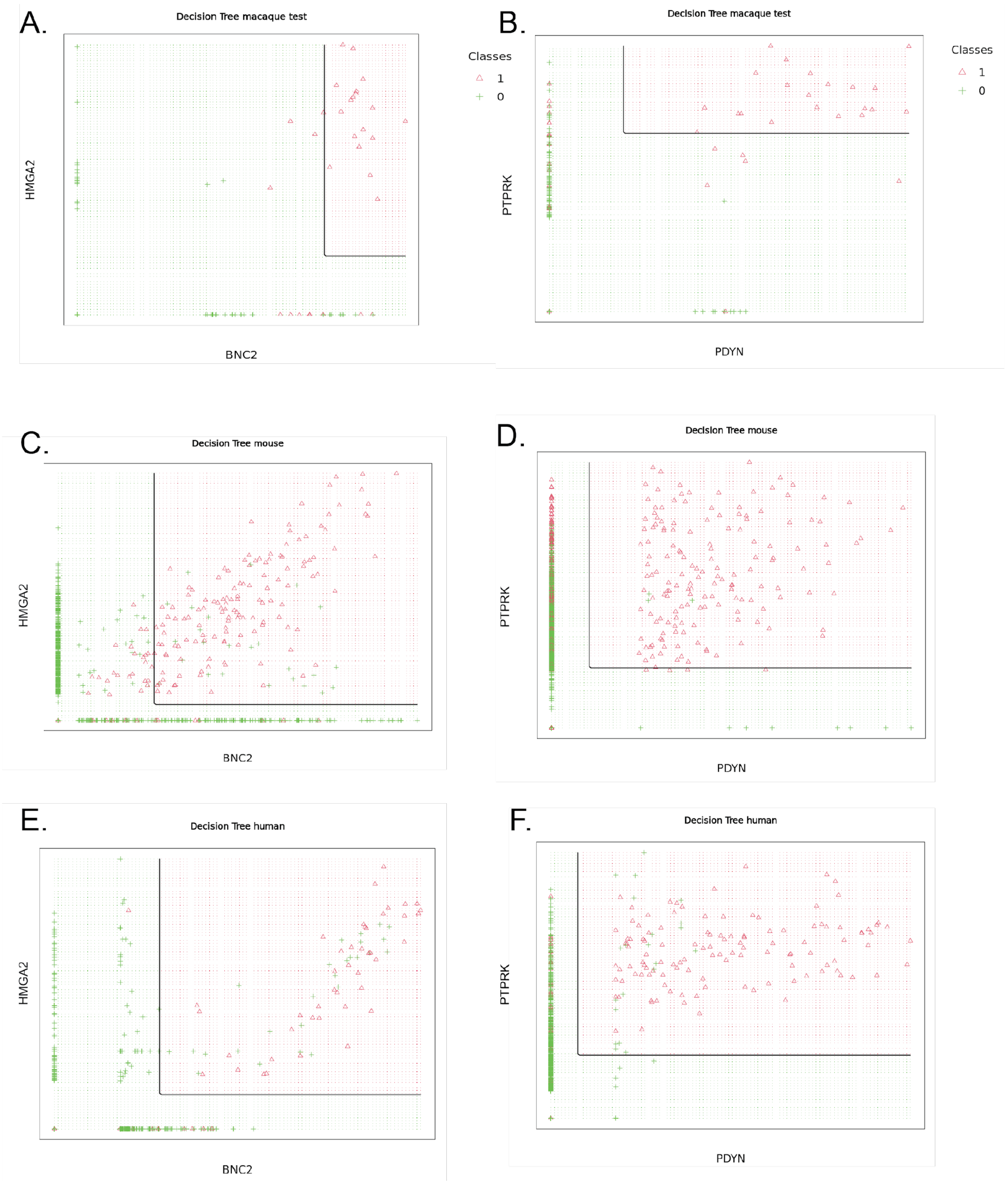


**Supplemental Fig. S13. Example decision trees for the Exc-BNC2 and Inh-PDYN cell types, related to Main Figure 3.** The imputed gene expression of each dorsal horn cell is plotted in a joint two-gene space for the two marker genes of each target cell type. Red triangles represent the target cell type (class 1), and green crosses represent the other off-target cells (class 0). The black lines represent the gene expression decision boundaries previously learned by the decision tree using the macaque training data, and now applied to the test sets. The red and green backgrounds represent the spaces on each side of the decision boundaries predicted to be target (red, class 1) or off-target (green, class 0). **A,C,E** Results for the Exc-BNC2 cell type for Macaque, Mouse, and Human test sets, respectively. **B,D,F** Results for the Inh-PDYN cell type for Macaque, Mouse, and Human test sets, respectively.

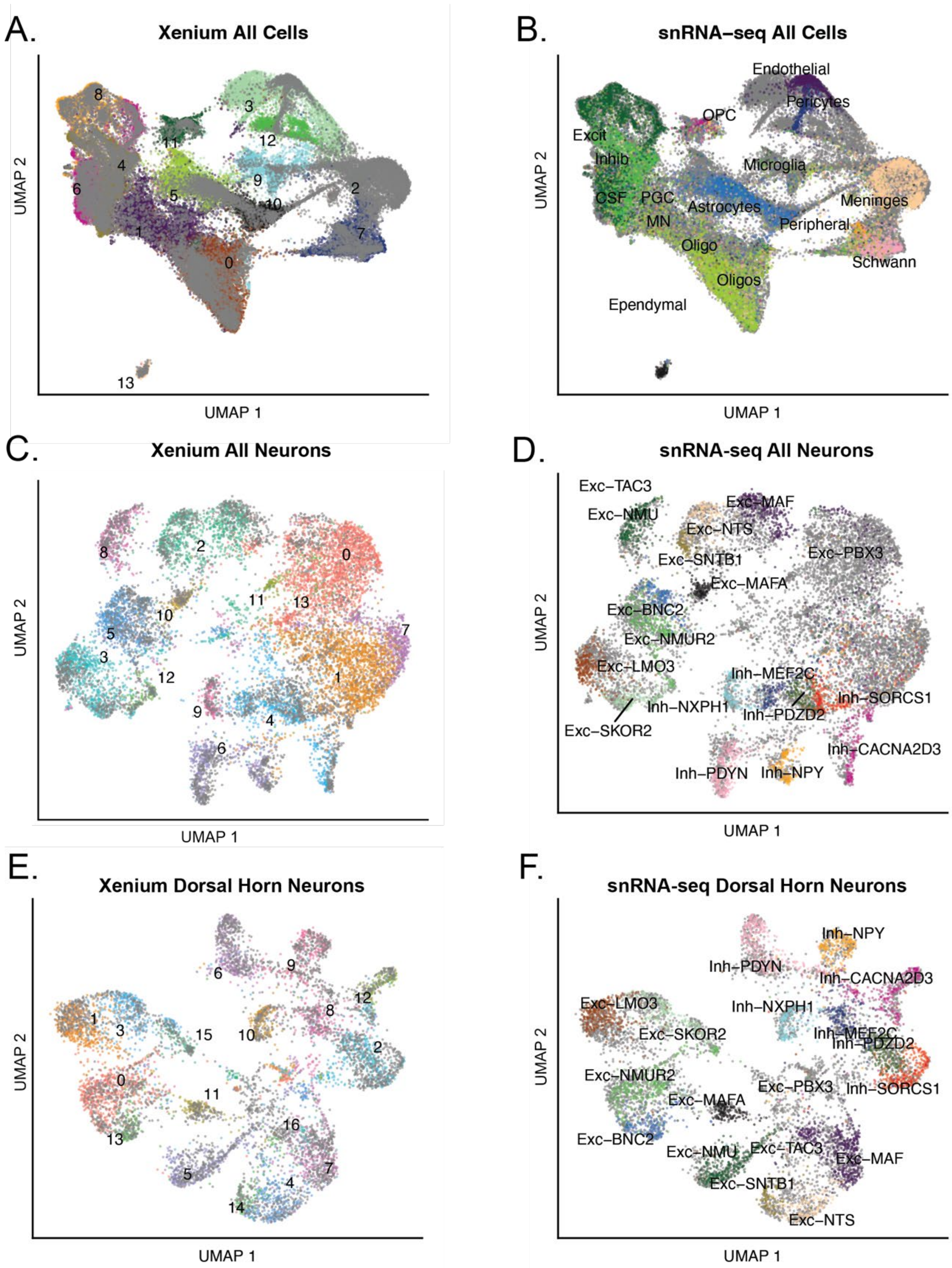


**Supplemental Fig. S14. UMAP representations of Mouse Xenium-snRNA iterative integrations to label cell types, related to Main Figure 4. A,B** Integrated and overlayed UMAP of all spinal cord cells in **A,** Xenium assay and **B,** pre-labeled snRNA-seq. Major Xenium cell types were manually annotated based on relative cluster positioning, and neurons were re-clustered for the next round of integration. **C,D,** Integrated and overlayed UMAP of **C,** all spinal neurons in Xenium, and **D,** pre-labeled dorsal horn neurons from snRNA-seq. Clusters 0,1,7,11, and 13 were labeled as non-dorsal horn, remaining neurons were re-clustered for the third round of integration. **E,F,** Integrated and overlayed UMAP of **E,** putative dorsal horn neurons in Xenium, and **D,** pre-labeled dorsal horn neurons from snRNA-seq. Xenium clusters were manually annotated as dorsal horn subtypes based on relative cluster positions.

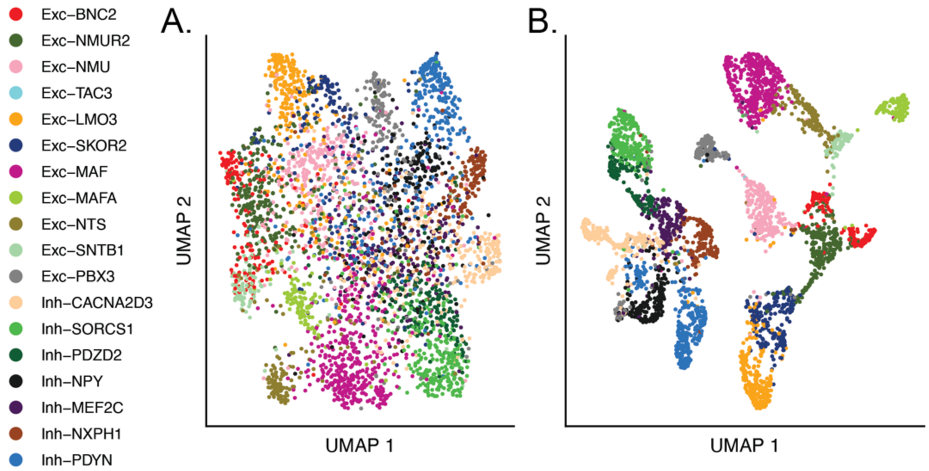


**Supplemental Fig. S15. Mouse snRNA-seq limited to the Xenium gene panel with and without imputed gene expression, related to Main Figure 4. A,** UMAP representation of dorsal horn cell types without gene imputation. **B,** UMAP representation of dorsal horn cell types using gene imputation to simulate the higher sensitivity of Xenium imaging-based detection of transcripts.

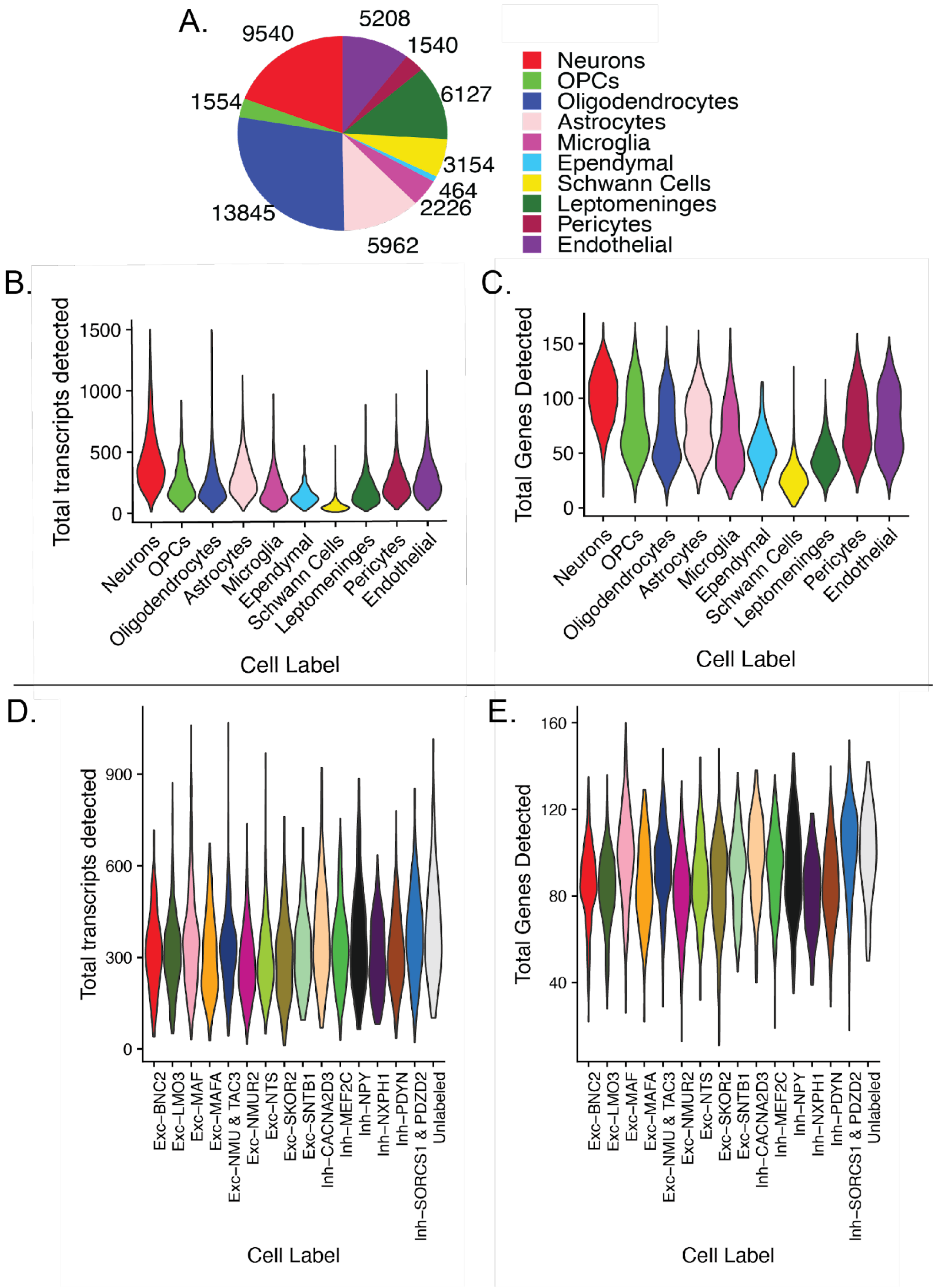


**Supplemental Fig. S16. Cell class proportions and quality control metrics of labeled Xenium clusters, related to Main Figure 4. A,** Pie chart of major cell types after manual annotation of mouse Xenium. Numbers outside of the pie chart indicate total cell counts. **B,C** Violin plot distributions for each major cell class of **B,** total detected transcripts and **C,** total detected unique genes. **D,E** Violin plot distributions for each dorsal horn subtype of **D,** total detected transcripts and **E,** total detected unique genes.

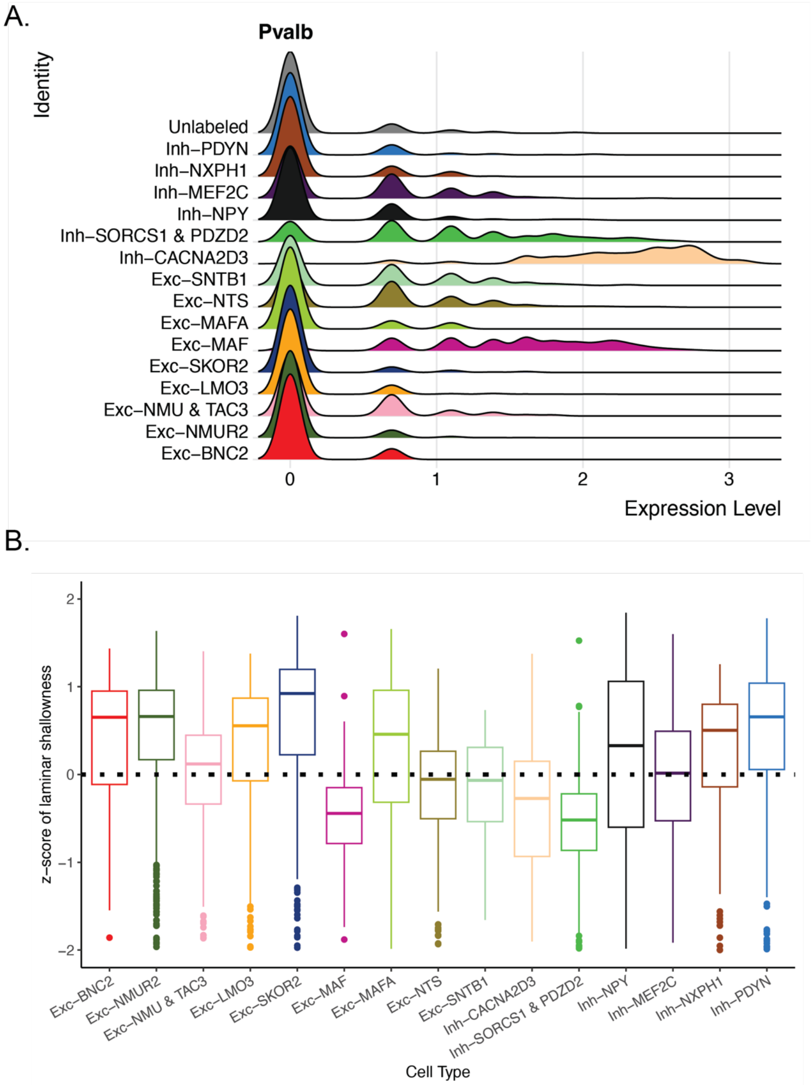


**Supplemental Fig. S17. Xenium dorsal horn parvalbumin expression and per-cell laminar distributions, related to Main Figure 4. A,** Ridge plot of parvalbumin (Pvalb) expression in labeled Xenium dorsal horn subtypes, with highest levels in the Inh-CACNA2D3 cell type. **B,** Boxplot of per-cell distributions of laminar distribution stratified by dorsal horn subtype. Alternative to **Figure 4F**, which shows the distribution of the average laminar deviations per-slide. z-score of laminar shallowness defined in Methods:

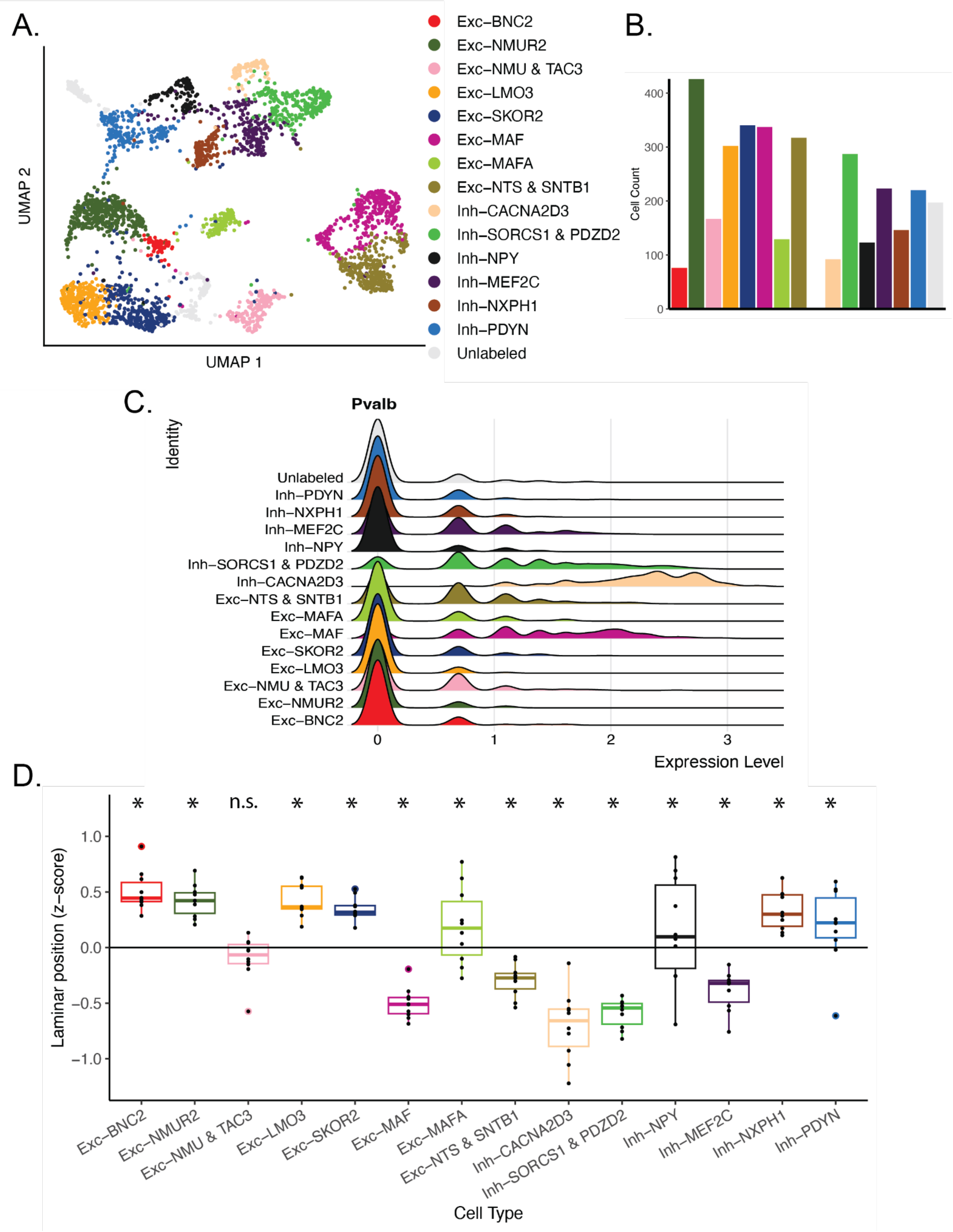
A.
UMAP 2
UMAP 1
Exc–BNC2
Exc–NMUR2
Exc–NMU & TAC3
Exc–LMO3
Exc–SKOR2
Exc–MAF
Exc–MAFA
Exc–NTS & SNTB1
Inh–CACNA2D3
Inh–SORCS1 & PDZD2
Inh–NPY
Inh–MEF2C
Inh–NXPH1
Inh–PDYN
Unlabeled
B.
Cell Count
0
100
200
300
400
C.
Pvalb
Identity
Unlabeled
Inh–PDYN
Inh–NXPH1
Inh–MEF2C
Inh–NPY
Inh–SORCS1 & PDZD2
Inh–CACNA2D3
Exc–NTS & SNTB1
Exc–MAFA
Exc–MAF
Exc–SKOR2
Exc–LMO3
Exc–NMU & TAC3
Exc–NMUR2
Exc–BNC2
0
1
2
3
Expression Level
D.
Laminar position (z–score)
1.0
0.5
0.0
−0.5
−1.0
* * n.s. * * * * * * * * * * *
Exc–BNC2
Exc–NMUR2
Exc–NMU & TAC3
Exc–LMO3
Exc–SKOR2
Exc–MAF
Exc–MAFA
Exc–NTS & SNTB1
Inh–CACNA2D3
Inh–SORCS1 & PDZD2
Inh–NPY
Inh–MEF2C
Inh–NXPH1
Inh–PDYN
Cell Type

**Supplemental Fig. S18. Replication of Xenium Laminar distributions and parvalbumin expression in second mouse, related to Main Figure 4.** The following results were based on the same analysis steps as Figure 4. **A,** UMAP representation of dorsal horn neuron subtypes identified in Xenium assay based on integration with adult mouse snRNA-seq dataset with conserved labels. Inh-SORCS1 and Inh-PDZD2, and Exc-NTS and Exc-SNTB1, were each combined, as cell types were not clearly distinguishable in Xenium. Two clusters indicated as Unlabeled did not clearly correspond to any snRNA-seq dorsal horn cell type. **B,** Bar chart of cell counts per neuron subtype in Xenium. **C,** Ridge plot showing distributions of parvalbumin (Pvalb) expression in each neuron subtype, with highest expression in Inh-CACNA2D3. **D,** Box plot of laminar position per cell type. z-score of laminar position was the vertical position of an individual cell relative to the average vertical position of its slice and normalized by the variability in vertical position of the slice. By convention of image orientation, a higher z-score indicates a more superficial position. Individual points of each box plot represent the average z-score for the cells of that cell type in one slice. * significant p-value indicating difference from zero (either significantly deep or superficial), after false discovery rate correction of 0.05.

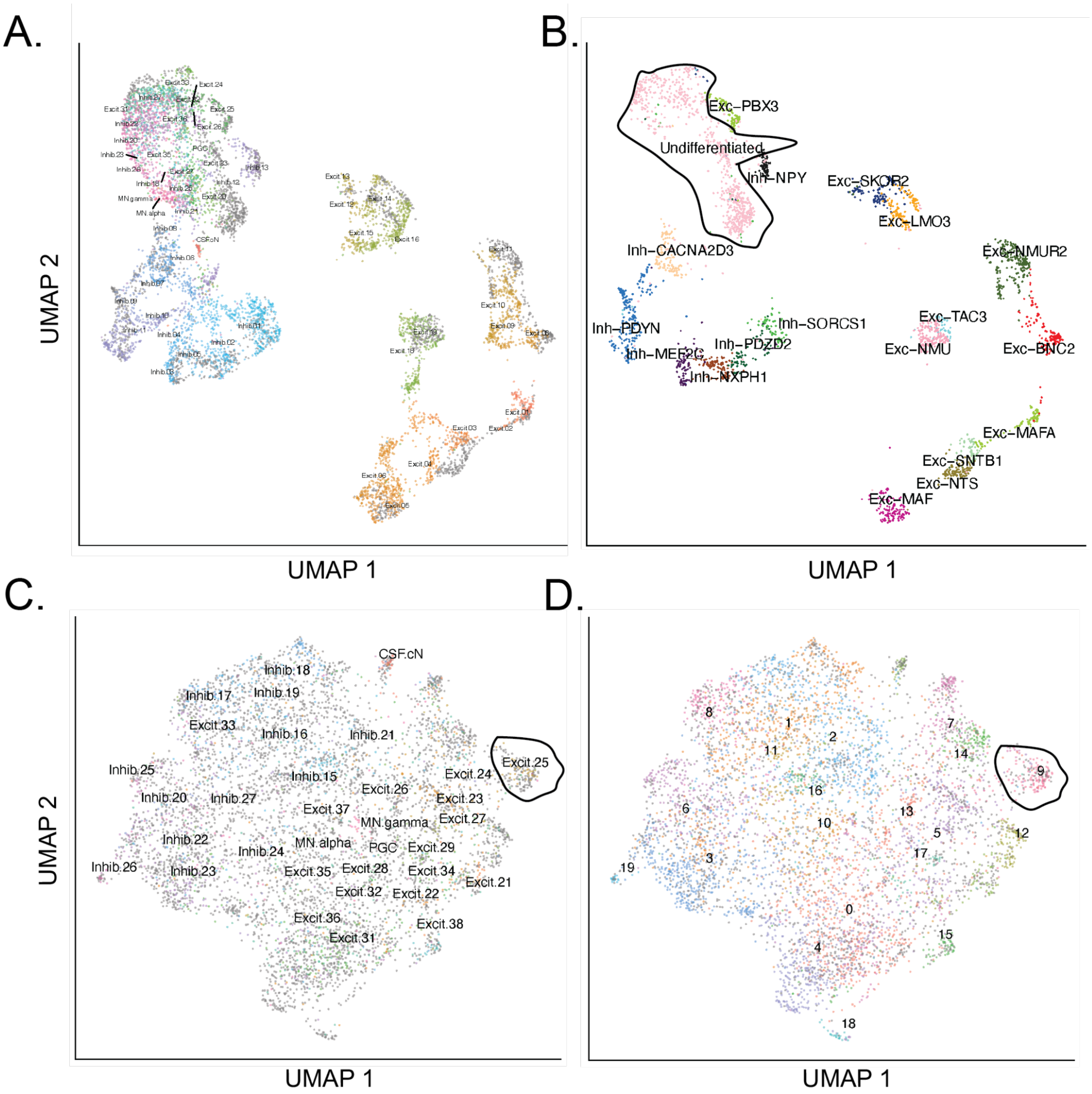


**Supplemental Fig. S19. Further Characterization of SPP1 family neurons, previously called MidVent, Part 1, related to Main Figure 5.** In **A,B**, the full set of macaque snRNA-seq nuclei (**B**) is integrated with the adult mouse subset of the mouse harmonized atlas (**A**), and the undifferentiated cells in macaque are aligned with several populations referred to as MidVent in the harmonized atlas. **C,D,** the Xenium clusters that did not correspond to a conserved subtype (**D**) are integrated with the MidVent neurons from the harmonized atlas (**C**). Here we see that clear separations of distinct clusters are not readily apparent, although cluster xen.9 in Xenium closely is aligned with Excit.25 (previously shown to be homologous to Exc-PBX3/PDYN) of the harmonized atlas.

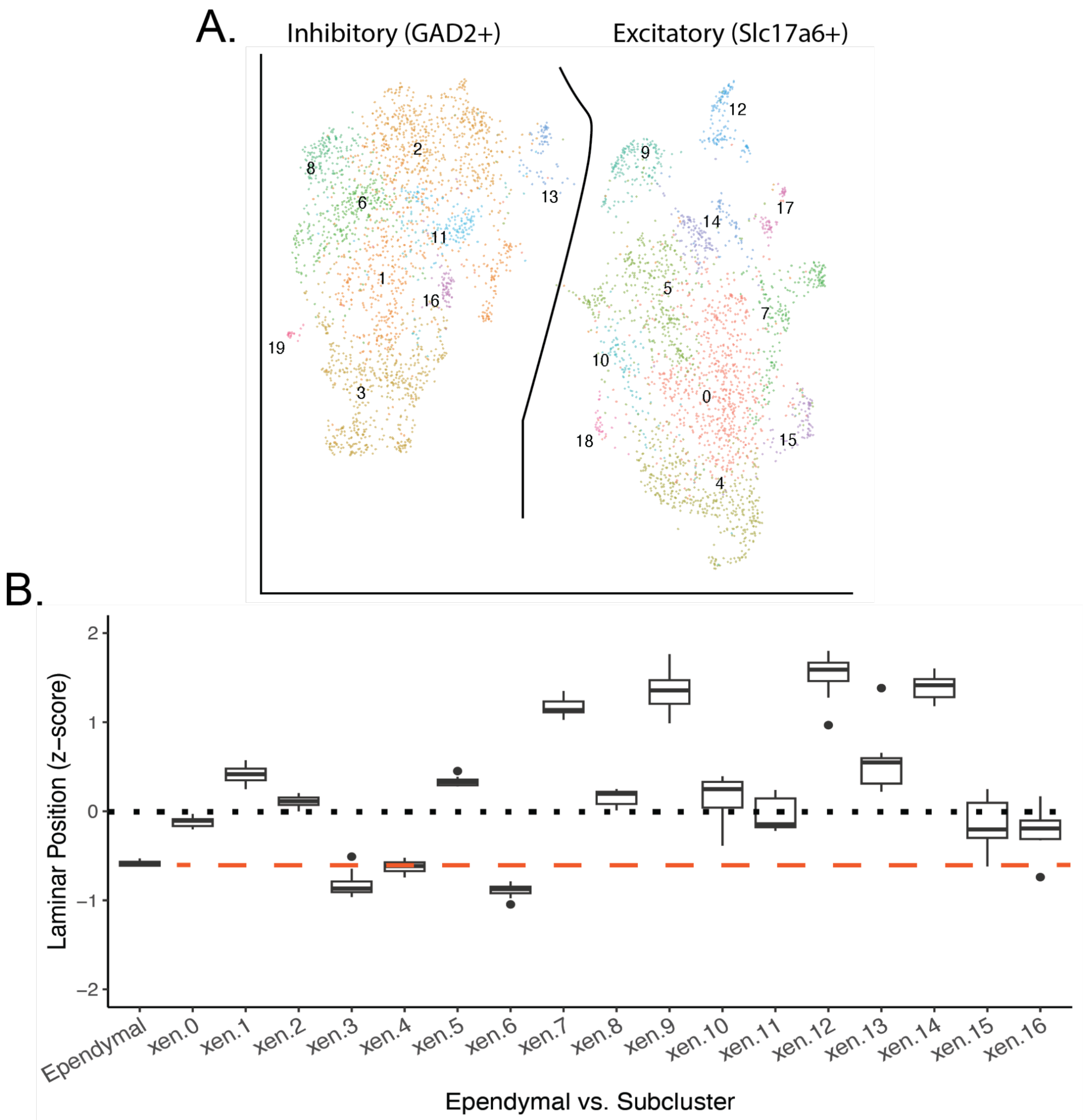


**Supplemental Fig. S20. Further Characterization of SPP1 family neurons, previously called MidVent, Part 2, related to Main Figure 5. A**. Unbiased clustering of the Xenium neurons that did not correspond to conserved neuron subtypes. Left-most clusters were enriched for the inhibitory marker Gad2, while the right-most clusters were enriched for the excitatory marker Slc17a6. **B**. Laminar distributions were calculated as in Figure 4, with ependymal cells (Ependymal) as the marker for the central canal. Several sub-clusters have higher laminar z-score than ependymal scores indicating that they are within the traditional boundary of the dorsal horn.

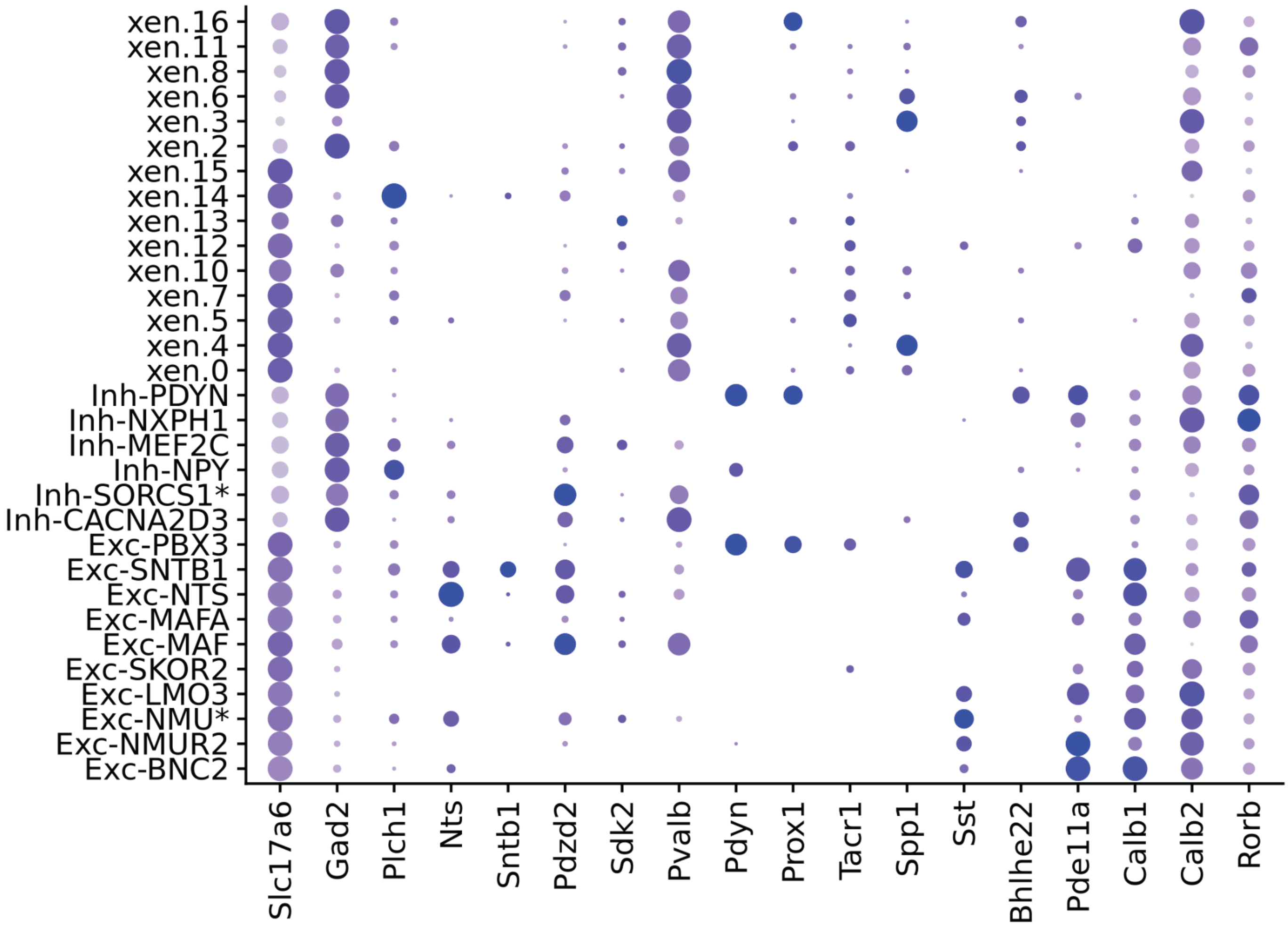


**Supplemental Fig. S21. Expression of selected genes in Xenium subtypes and subclusters, related to Main Figure 5.** Dot Plot of gene expression patterns for excitatory/inhibitory markers (Slc17a6/Gad2), the conserved gene markers available in the Xenium panel (Plch1, Nts, Sntb1, Pdzd2, Sdk2, Pdyn), and select other genes of interest. The size of the dot indicates the relative proportion of cells expressing the gene. Expression shown for Spp1 family sub-clusters (xen.0-16) and conserved neuron subtypes.

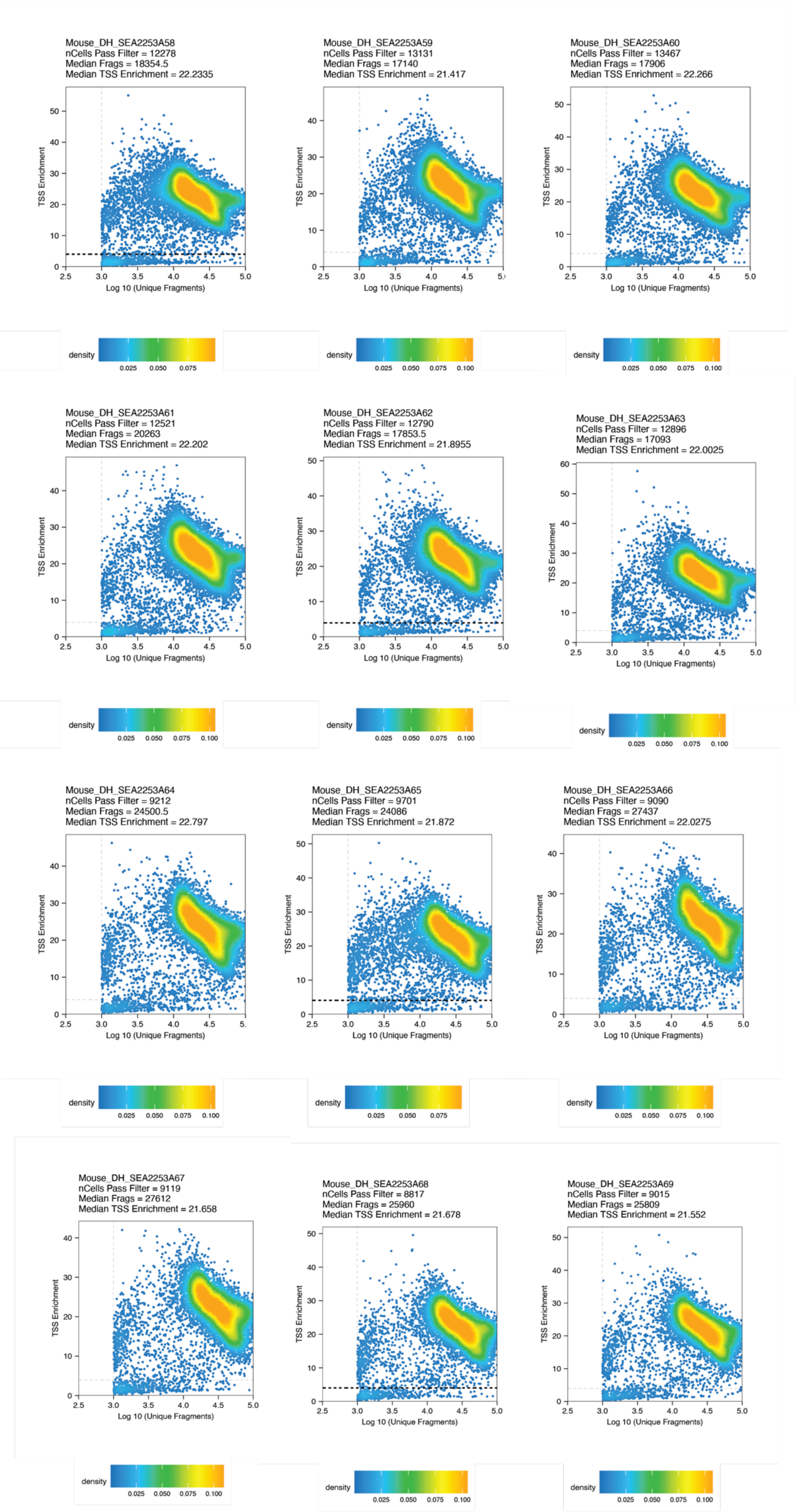
Mouse_DH_SEA2253A58
nCells Pass Filter = 12278
Median Frags = 18354.5
Median TSS Enrichment = 22.2335
Mouse_DH_SEA2253A59
nCells Pass Filter = 13131
Median Frags = 17140
Median TSS Enrichment = 21.417
Mouse_DH_SEA2253A60
nCells Pass Filter = 13467
Median Frags = 17906
Median TSS Enrichment = 22.266
Mouse_DH_SEA2253A61
nCells Pass Filter = 12521
Median Frags = 20263
Median TSS Enrichment = 22.202
Mouse_DH_SEA2253A62
nCells Pass Filter = 12790
Median Frags = 17853.5
Median TSS Enrichment = 21.8955
Mouse_DH_SEA2253A63
nCells Pass Filter = 12896
Median Frags = 17093
Median TSS Enrichment = 22.0025
Mouse_DH_SEA2253A64
nCells Pass Filter = 9212
Median Frags = 24500.5
Median TSS Enrichment = 22.797
Mouse_DH_SEA2253A65
nCells Pass Filter = 9701
Median Frags = 24086
Median TSS Enrichment = 21.872
Mouse_DH_SEA2253A66
nCells Pass Filter = 9090
Median Frags = 27437
Median TSS Enrichment = 22.0275
Mouse_DH_SEA2253A67
nCells Pass Filter = 9119
Median Frags = 27612
Median TSS Enrichment = 21.658
Mouse_DH_SEA2253A68
nCells Pass Filter = 8817
Median Frags = 25960
Median TSS Enrichment = 21.678
Mouse_DH_SEA2253A69
nCells Pass Filter = 9015
Median Frags = 25809
Median TSS Enrichment = 21.552
TSS Enrichment
Log 10 (Unique Fragments)
density

**Supplemental Fig. S22-23. Quality control (TSS enrichment and unique fragments) of mouse snATAC-seq by sample, related to Main Figure 5.** Each panel corresponds to a technical replicate. For each panel, dots correspond to individual nuclei, with corresponding log10 of unique fragments (x-position) and average TSS enrichment (y-position). Cells located in the top right quadrant of the dotted lines passed minimum thresholds of TSS enrichment = 3.5 and log10 fragments = 3.0 and were kept for further analysis. As shown by the high density in the top right sections of the figures, the vast majority of cells had TSS enrichment > 10. TSS = transcription start site.

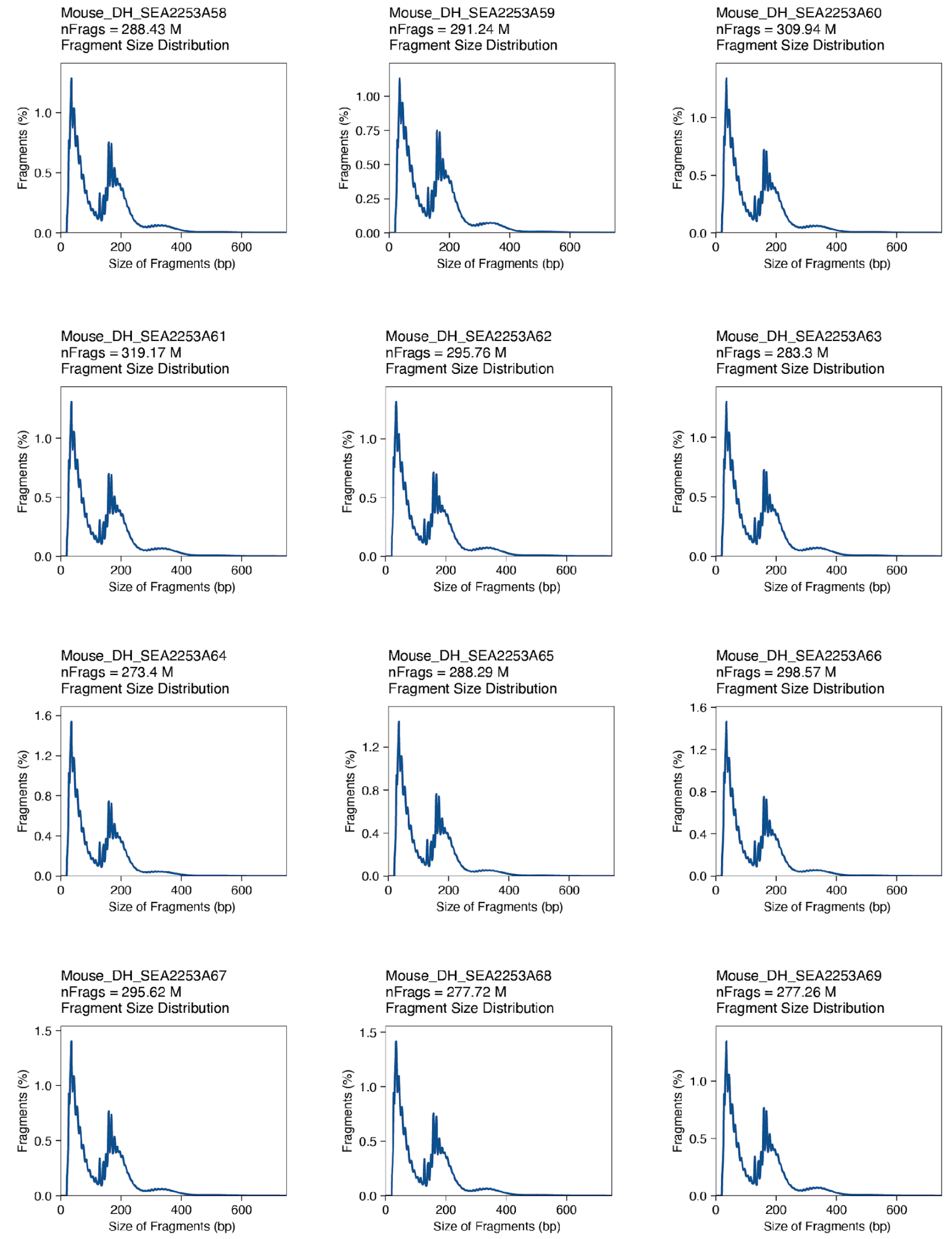


**Supplemental Fig. S24. Quality control (fragment size distributions) of mouse snATAC-seq by sample, related to Main Figure 6.** Each panel corresponds to a technical replicate. For each panel, normalized histograms of fragment sizes per replicate are shown. All samples show a characteristic periodicity of relatively low abundance of fragments every multiple of 140 base pairs, corresponding to the length of nucleosome(s) inaccessible to the Tn5 transposase, an indicator of intact regulation of open and closed chromatin in the cell.
nFrags = the total number of fragments per technical replicate. bp= base pairs.

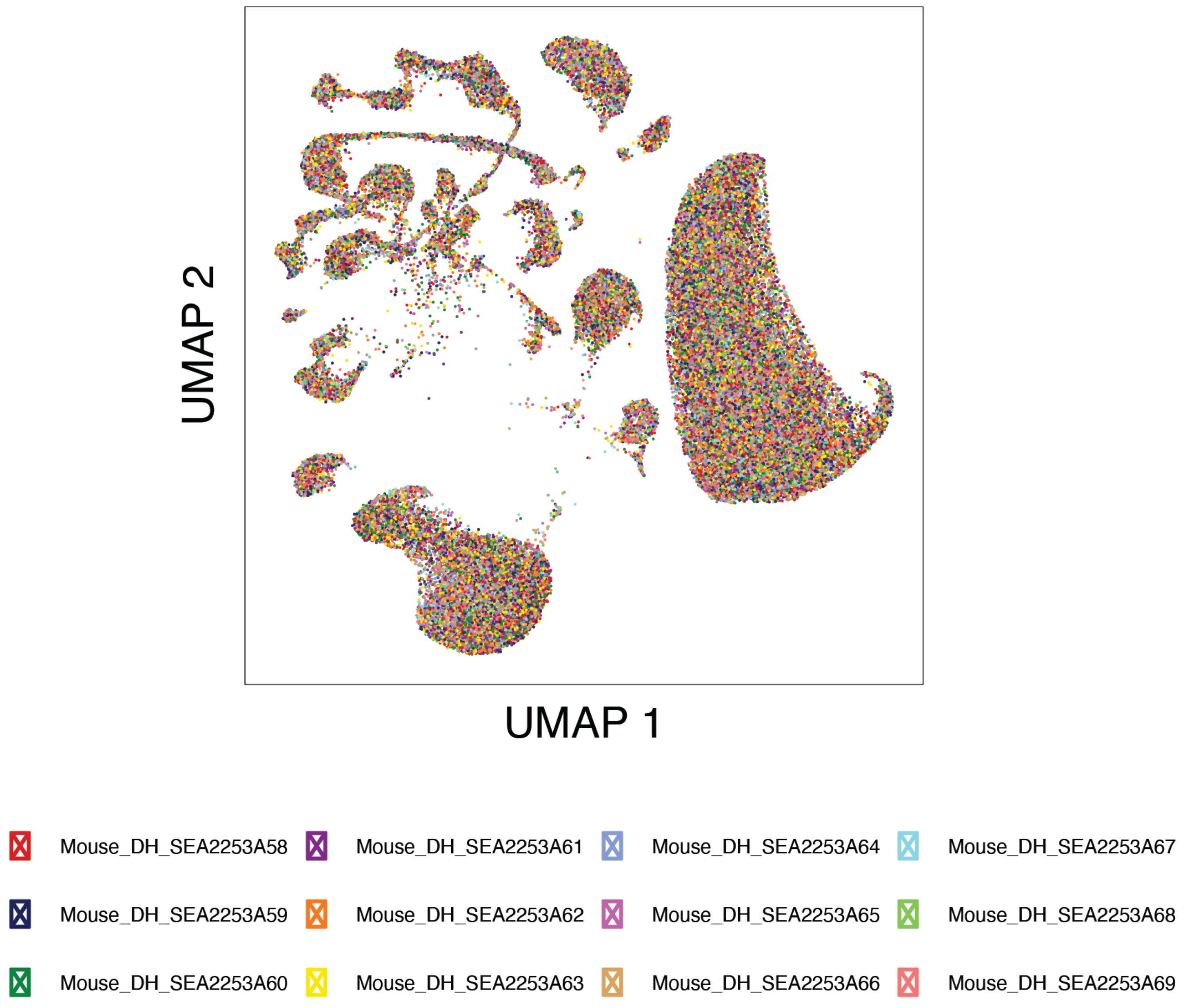


**Supplemental Fig. S25, Successful Batch Correction of Mouse snATAC-seq, related to Main Figure 6.** UMAP visualization of single nuclei, colored by sample. All clusters show mixtures of nuclei from each sample, without clear differences sample-wise differences, demonstrating the results of batch correction.

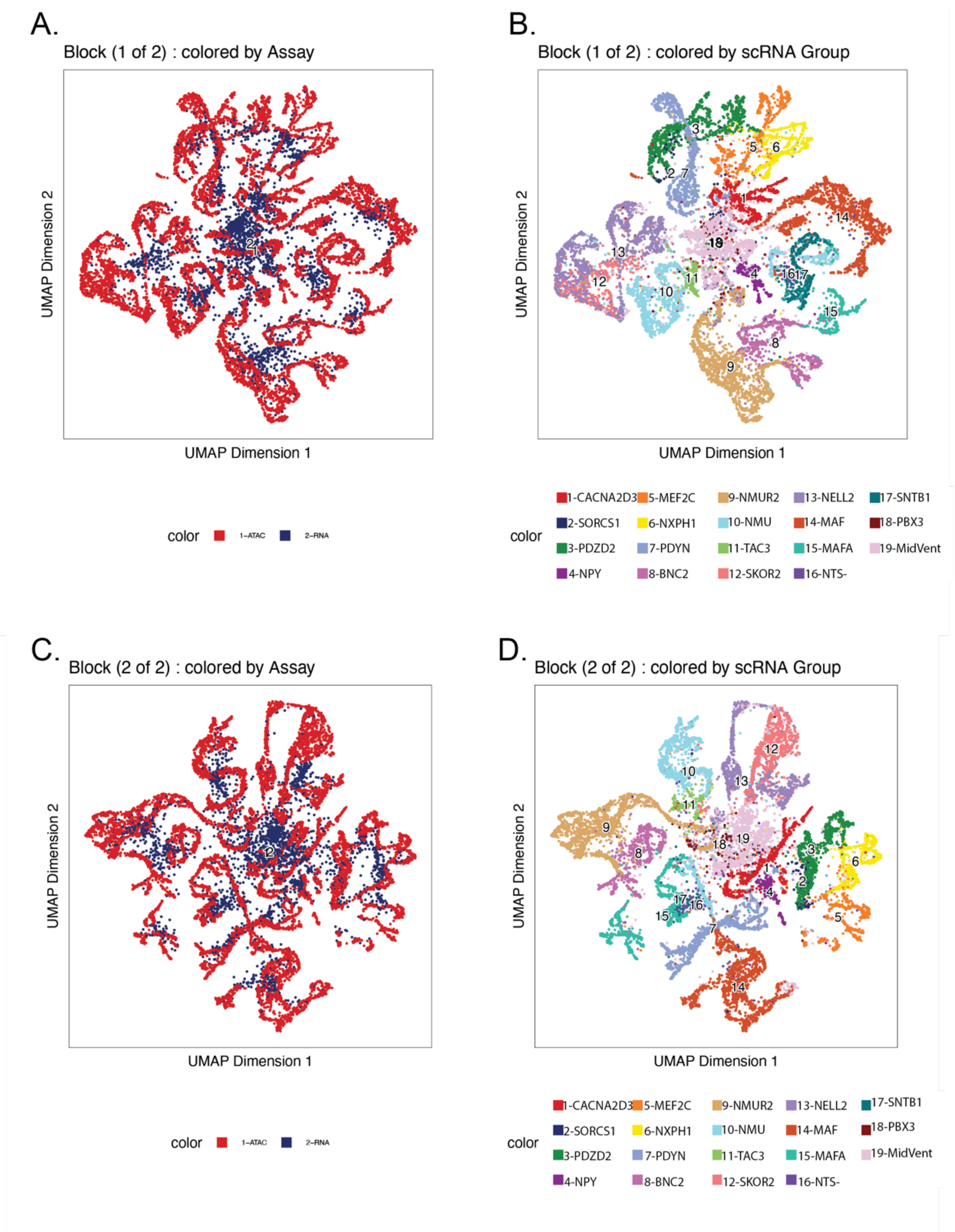


**Supplemental Fig. S26. ATAC-RNA Co-Clustering, related to Main Figure 6.** ATAC-RNA integration creates a shared co-embedding of RNA and ATAC nuclei, from which nearest neighbors are calculated for label transfer. Two blocks of co-embedding were generated for parallelism. UMAP visualizations are shown for each block: A,C (left): co-embedding of ATAC nuclei (red) and RNA nuclei (blue). B,C (right): ATAC labels after nearest neighbors labeling.

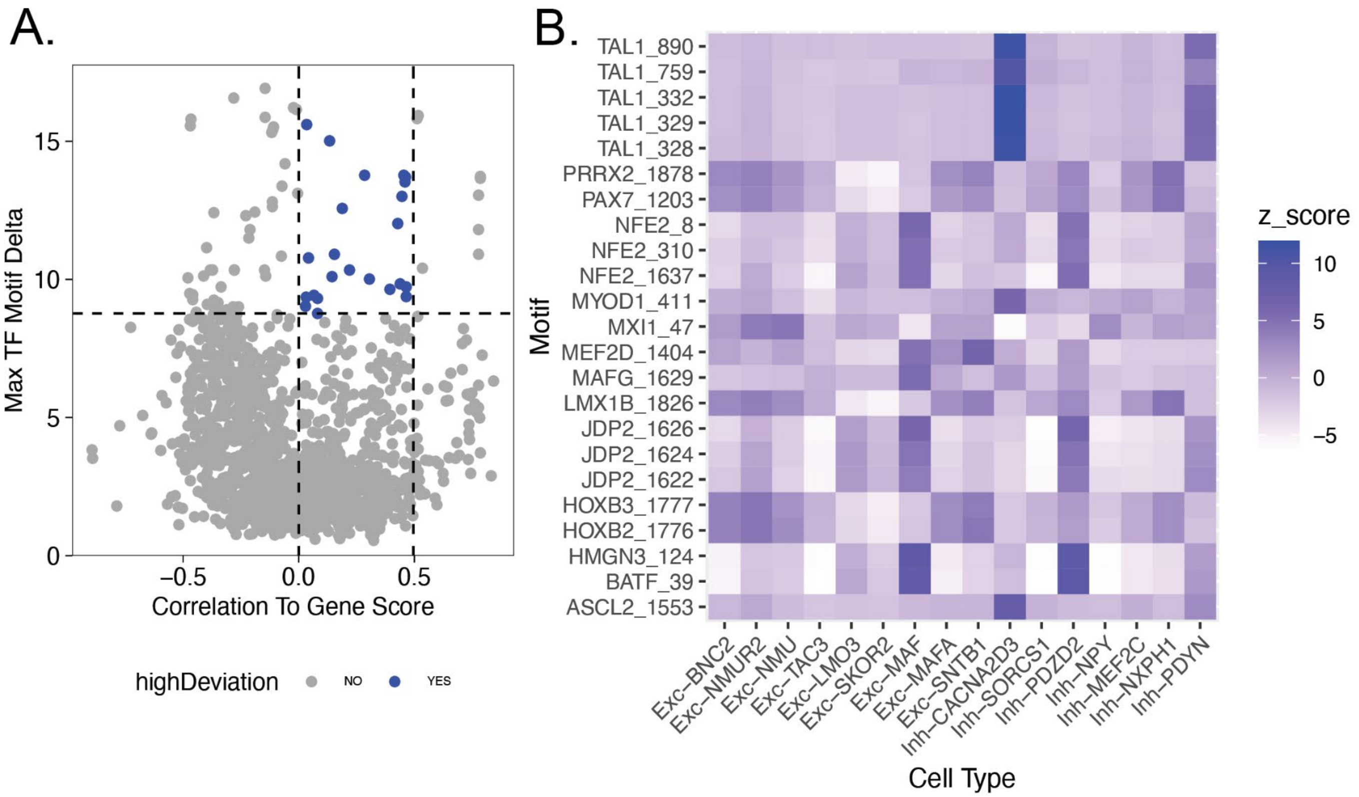


**Supplemental Fig. S27. Transcription factor high-deviation candidates, related to Main Figure 6. A,C,** Each dot corresponds to an individual motif. The y-position is the highest chromVar deviation of a motif (motif delta) across cell types. Higher motif delta indicates higher cell-type-specificity. The x-position is the per-cell correlation of gene expression of the TF with its motif enrichment within the cell. Higher correlation suggests a relationship between production of the TF gene product and its function as a trans-acting regulator in cell open chromatin. **A,** blue indicates motifs selected as high deviation candidates of interest based on the 95$^{th}$ or greater quantile of motif delta, and gene expression correlation > 0, and not already identified as a TF regulator as in **A**. **B,** Heatmap of motifs selected in **A** and their motif delta (z-score) across dorsal horn subtypes.
TF = transcription factor.

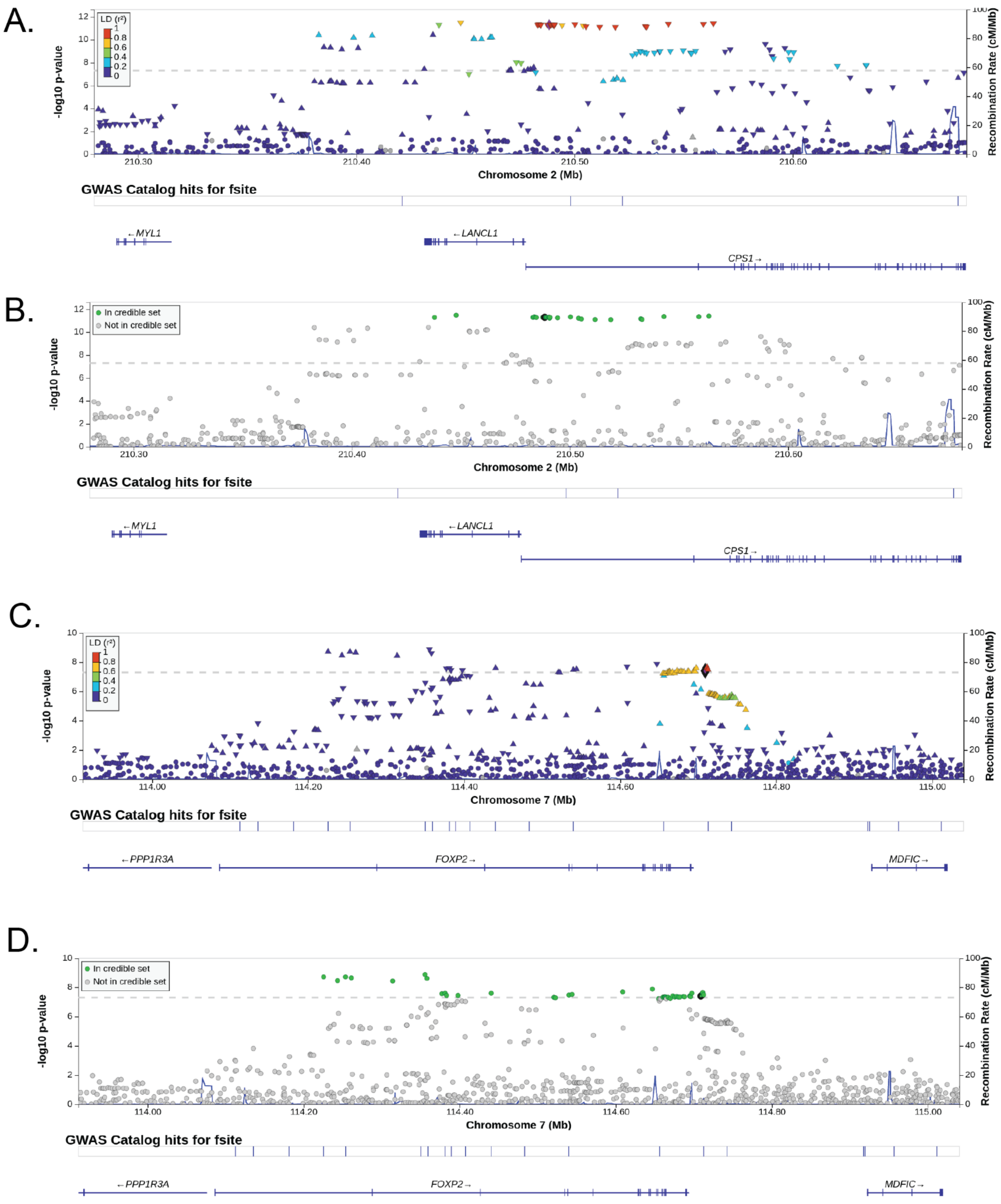


**Supplemental Fig. S28. LocusZoom tracks for SNP candidates nearby LANCL1 and FOXP2 , related to Main Figure 6.** Manhattan plots of SNP nearby the LANCL1 locus (A,B) and FOXP2 (**C,D**). In **A,C**, linkage disequilibrium (LD) statistics of SNPs near the SNP of interest are shown, demonstrating that the SNP of interest is at least similarly significant or more significant than SNPs with which it is in high LD. In B,D, fine-mapping estimates by LocusZoom are shown, where green dots indicate the SNP is part of the credible set of potentially causal SNPs. In both cases, the SNP of interest is part of the credible set.
